# A Split-Luciferin Assay for Real-Time Measurement of Cytosolic Accumulation in Live Mycobacteria

**DOI:** 10.1101/2025.11.02.686125

**Authors:** Rachita Dash, Andrew Spira, Irene Lepori, Sarah E. Newkirk, Sobika Bhandari, Noah Lucas, Mahendra D. Chordia, M. Sloan Siegrist, Marcos M. Pires

## Abstract

Antibiotics against nontuberculous mycobacteria must cross a multilayered cell envelope before reaching their cytosolic targets, yet how much of a given compound arrives there is largely unmeasured. This gap matters because nontuberculous mycobacterial infections are rising among immunocompromised individuals and those with structural lung disease, and intrinsic and acquired multidrug resistance creates an urgent need for new agents. Existing methods quantify whole-cell association without resolving subcellular localization, and none report accumulation in live mycobacteria, in real time. Here we adapt a split-luciferin approach, the D-cysteine-linked accumulation profiling (D-CLAP), to luciferase-expressing *Mycobacterium smegmatis* and *Mycobacterium avium*. Compounds bearing a disulfide-linked D-cysteine tag release the tag upon entry into the reducing cytosol, where it condenses with an intracellular cyanobenzothiazole to form D-luciferin and generate a time-resolved luminescent signal. We applied D-CLAP to small-molecule antibiotics, polyarginine peptides, and griselimycin analogs, and found that masking carboxylates as methyl esters increased both cytosolic accumulation and antimycobacterial potency. A fluorescent polyarginine peptide also associated with *M. smegmatis* residing within macrophage phagosomes. D-CLAP provides a real-time readout of cytosolic entry in live mycobacteria and a route to structure-accumulation relationships.

## INTRODUCTION

Nontuberculous mycobacteria (NTM) cause chronic pulmonary, skin, and disseminated infections, and their incidence has risen steadily worldwide in recent decades.^1,2^ Despite this growing burden, effective therapeutic options remain limited, especially given the intrinsic and acquired multidrug resistance of these organisms.^3^ Antibiotic development against NTM is particularly challenging because of the complex, multilayered cell envelope these organisms share with *Mycobacterium tuberculosis* (*Mtb*).^4,5^ The cell envelope includes an outer mycomembrane, an arabinogalactan layer, a peptidoglycan (PG) layer, and an inner cytoplasmic membrane.^6^ To reach intracellular targets, antibiotics must traverse all of these barriers (**Fig. 1a**). Given that most antimycobacterial agents act on intracellular processes,^7,8^ successful drug development requires not only target engagement but also a thorough understanding of a compound’s ability to penetrate the cell envelope and access its site of action.

**Figure 1.**
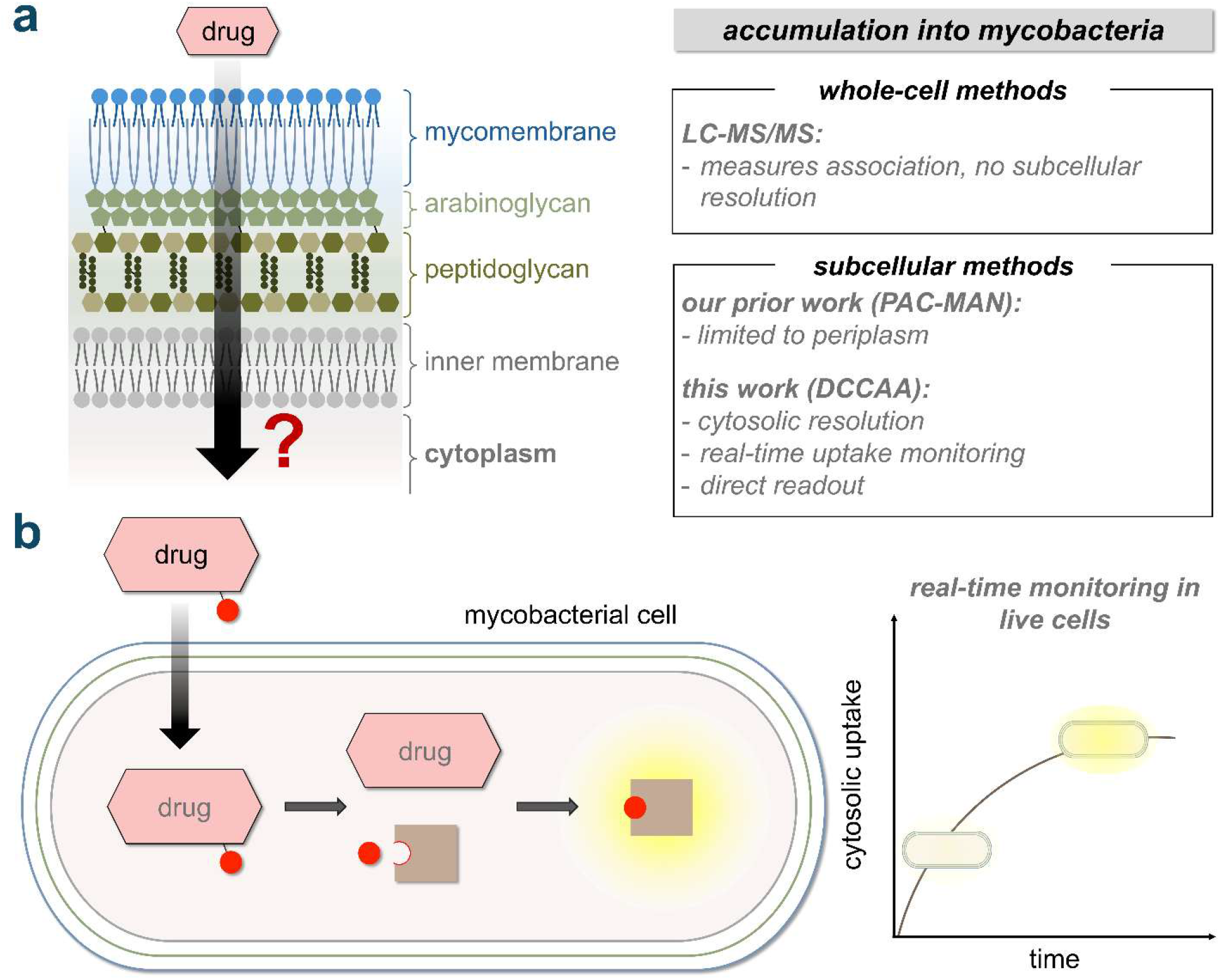
**(a)** Structure of the mycobacterial cell envelope and previous work done to assess molecular accumulation in mycobacteria. **(b)** Conceptual overview of D-CLAP and the barriers it resolves.

Advancing antimycobacterial drug development requires improved methodologies to assess the intracellular accumulation of compounds within mycobacterial cells. Such approaches not only provide fundamental insights into the molecular features that govern intracellular accumulation, but also serve as practical tools for screening during high-throughput drug discovery campaigns. In pursuit of this goal, our laboratory has developed and implemented several methods to investigate molecular accumulation in bacteria,^9–14^ including the Peptidoglycan Accessibility Click-Mediated AssessmeNt (PAC-MAN) assay, which employs click chemistry to quantify molecular accumulation across the mycomembrane in live mycobacteria.^15,16^ While informative, PAC-MAN is limited to assessing periplasmic accumulation and does not resolve cytosolic penetration (**Fig. 1a**).

Liquid chromatography-tandem mass spectrometry (LC-MS/MS) remains the benchmark for quantifying whole-cell-associated molecules without chemical modification.^17,18^ LC-MS/MS has also been used to study accumulation in mycobacteria.^19,20^ However, LC-MS/MS lacks subcellular resolution, providing only bulk measurements across the entire cell (**Fig. 1a**). Moreover, it is limited to end-point analyses and cannot capture the temporal dynamics of accumulation. These constraints underscore the need for methodologies capable of directly assessing real-time cytosolic accumulation.

We previously developed a bioluminescence-based D-cysteine (D-cys) cytosolic accumulation assay to measure cytosolic accumulation in Gram-negative bacteria.^11^ Here, we adapt this approach to mycobacteria, enabling real-time quantification of cytosolic uptake (**Fig. 1b**).^21^ The assay employs luciferase-expressing mycobacterial cells treated with a 2-cyanobenzothiazole (CBT) probe alongside test compounds tagged with a D-cys moiety *via* a disulfide linkage (**Fig. 2a**). Upon cytosolic entry, the reducing environment cleaves the disulfide bond, releasing D-cys, which rapidly undergoes bioorthogonal condensation with CBT in cyto to form D-luciferin. D-luciferin is then processed by luciferase, generating quantifiable bioluminescence (**Fig. S1a**).^22^ In this study, we used D-CLAP to monitor the real-time cytosolic accumulation of a range of small molecules and peptides in live mycobacteria. This represents the first assay capable of resolving cytosolic drug uptake in mycobacteria, addressing a critical gap in current methodologies.

**Figure 2.**
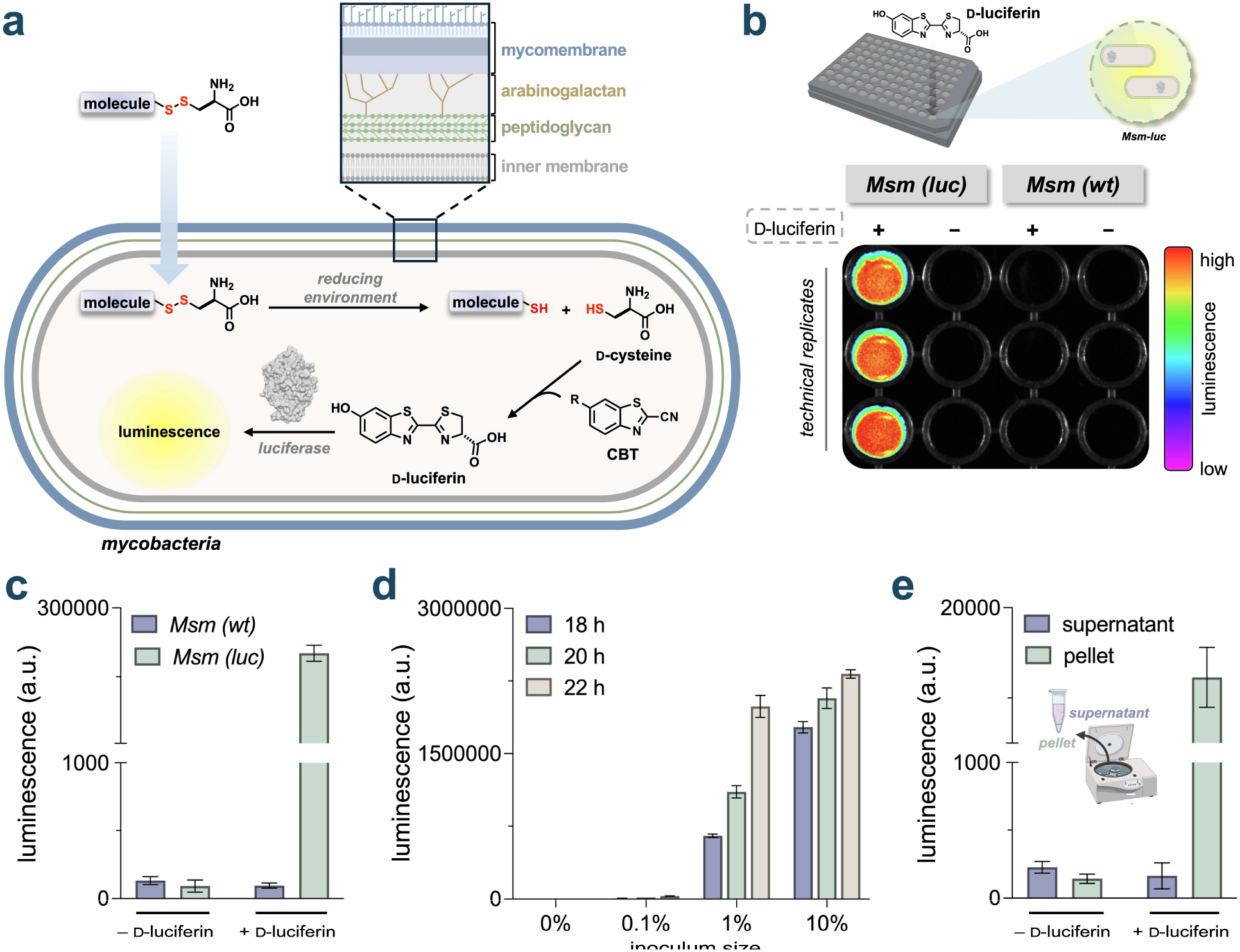
**(a)** Chemical basis of D-CLAP. **(b)** Luminescent imaging of *Msm (luc)* and *Msm (wt)* cells in the presence or absence of 100 μM D-luciferin. **(c)** Quantification of the luminescence depicted in (b). **(d)** Luminescence of *Msm (luc)* with varying inoculum sizes and growth durations after treatment with 25 μM D-luciferin. **(e)** Luminescence of *Msm (luc)* cell pellet and supernatant after centrifugation with subsequent addition or absence of 25 μM D-luciferin. Bars in (c – e) are represented as mean ± SD (*n* = 3).

## RESULTS

### Assay Development in Mycobacteria

We initially sought to build upon our prior assay parameters by employing *Mycobacterium smegmatis* (*Msm*), a model organism that recapitulates essential characteristics of pathogenic mycobacteria, including NTM species.^23^ Firefly luciferase was introduced into *Msm* by transformation with a previously optimized, constitutively active plasmid.^24^ To confirm expression, luciferase-expressing *Msm* (*Msm (luc)*) cells were grown to mid-log phase, incubated with D-luciferin, and subsequently imaged in a multiwell plate using a gel imager. Cells harboring the plasmid exhibited strong luminescence relative to the wild-type control (*Msm (wt)*), confirming luciferase expression (**Fig. 2b**). Luminescence was also quantified after a one-hour incubation with D-luciferin using a plate reader, which revealed a robust signal-to-noise ratio (**Fig. 2c**). These results confirmed that *Msm (luc)* supports luciferase-based detection of luciferin.

To optimize the intensity of the luminescence signal, we evaluated luciferase activity across varying growth durations and stationary phase inoculum sizes of *Msm (luc)* cells (**Fig. 2d**). A 22-hour growth period following a 1% inoculum was found to yield a strong signal-to-background ratio and was selected as the standard condition for the assay, with cultures reaching an OD_600_ of approximately 1. We also observed a correlation of luminescence with cell density, which is expected since higher cell numbers result in increased photon emission across the analyzed surface area (**Fig. S2**). To confirm that the luminescence signal originated from within the cells and was not due to an unrecognized source of extracellular luciferase activity, we separated the cells from the supernatant. *Msm (luc)* cells were incubated with D-luciferin for 30 minutes, followed by centrifugation to isolate the supernatant from the cellular pellet. After resuspension in buffer and an additional 30-minute incubation, luminescence measurements revealed near-background signal in the supernatant and a markedly stronger signal in the resuspended pellet. These results confirm that luciferin processing occurs exclusively within the cellular compartment, effectively ruling out contributions from extracellular luciferase activity (**Fig. 2e**).

We then modeled the accumulation assay using the split-luciferin system. The cytosol of mycobacteria is a reducing environment, owing primarily to the high intracellular concentration of mycothiol,^25,26^ which reduces the disulfide bond of a tagged compound and releases free D-cys for condensation with intracellular 6-hydroxy-2-cyanobenzothiazole (OH-CBT) to form D-luciferin. As a simple proxy for our test compound conjugates, we used D-cystine, the disulfide-linked dimer of D-cys (**Fig. 3a**). Mycobacterial cells were co-incubated with OH-CBT and D-cystine, and we monitored luminescence over time. Satisfyingly, we observed a robust luminescence level in *Msm (luc)* cells and no signal in *Msm (wt)*, confirming the luciferase dependence of the signal observed. This enabled monitoring of accumulation over time, which could be leveraged to elucidate how cellular factors might influence the kinetics of not only accumulation but also the cellular redox state.

**Figure 3.**
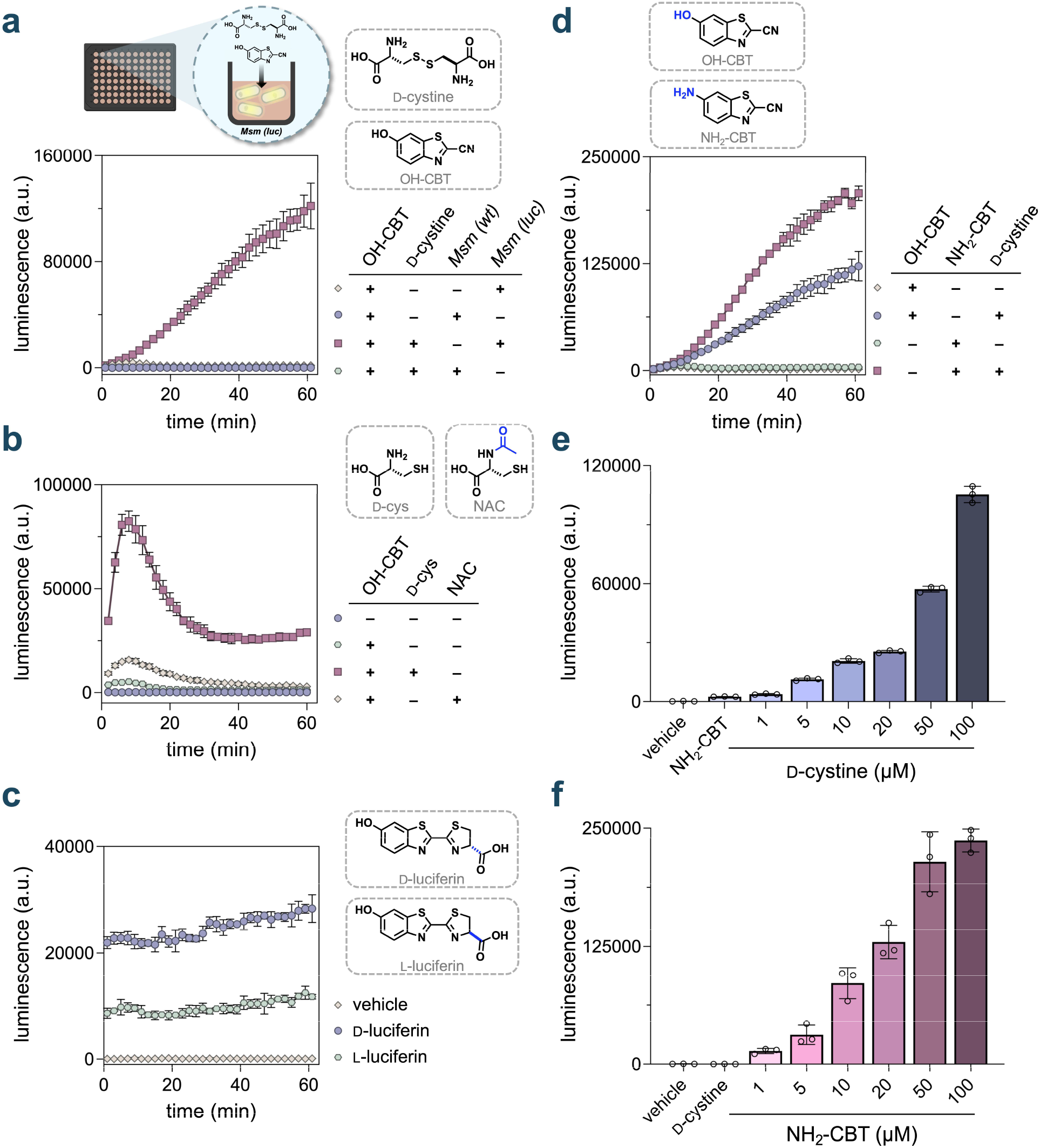
**(a)** Luminescence of *Msm (wt)* and *Msm (luc)* cells in the presence or absence of OH-CBT and D-cystine (both 50 μM). **(b – f)** Luminescence of *Msm (luc)* cells in the presence or absence of **(b)** OH-CBT, D-cys, and NAC (all 50 μM); **(c)** D- and L-luciferin (both 25 μM); **(d)** OH-CBT, NH_2_-CBT, and D-cystine (all 50 μM); **(e)** D-cystine (varying concentrations) and NH_2_-CBT (50 μM); **(f)** NH_2_-CBT (varying concentrations) and D-cystine (50 μM). Luminescence was measured over 60 min in (a – d) and at 60 min in (e – f) after addition of the indicated compounds. Data are represented as mean ± SD (*n* = 3).

We also observed a signal in the presence of the free amino acid D-cys. A key caveat in using free cysteine, or any thiol-containing molecule, in the oxidizing conditions of cell culture media is its rapid conversion to D-cystine, complicating attribution of observed effects to either the free thiol or its oxidized dimer (**Fig. 3b**). To test whether the amino group of D-cys is essential for its condensation with OH-CBT, we treated cells with *N*-acetyl-D-cysteine (NAC), which lacks a free amino group. The acetylation of the *N*-terminus of this amino acid effectively prevents participation in the OH-CBT condensation (**Fig. 3b**). Our results showed significantly reduced luminescence for NAC compared to D-cys. Interestingly, signals from NAC were still above background, suggesting possible endogenous *N*-deacetylase activity in mycobacterial cells. Overall, these findings highlight the requirement for free *N*-terminal cysteines in our workflow and confirm that luminescence arises from the bioorthogonal reaction between OH-CBT and D-cys. Incubation of *Msm* (luc) cells with OH-CBT alone also produced a detectable, albeit substantially lower, signal. We attribute this signal to the formation of L-luciferin in cyto, a phenomenon we previously observed in our analogous *E. coli* system.^11^ Endogenous cytosolic L-cysteine (L-cys) can undergo a condensation reaction with OH-CBT to yield L-luciferin,^21,27,28^ which has been reported to epimerize in the presence of firefly luciferase and subsequently serve as a substrate for the enzyme.^29^ Indeed, treatment of *Msm (luc)* cells with L-luciferin produced luminescence signals well above background levels, indicating substantial epimerase activity when the substrate is readily available (**Fig. 3c**). We propose that, within the context of our assay workflow, where the intracellular pool of endogenous L-cys is finite and limited, the formation of L-luciferin occurs rapidly following internalization of OH-CBT, but this pool is quickly depleted. Over time, the predominant source of cysteine for OH-CBT condensation shifts to the D-cys tag present on molecules arriving in the cytosol.

It has been well established that the chemical nature of the substituent at the 6-position of the benzothiazole ring can significantly influence luminescence output.^30,31^ Likewise, we have also previously noted this factor in luciferase-based analysis.^11^ Furthermore, we recognize that chemical modifications to OH-CBT may influence its intracellular accumulation and retention within target mycobacterial cells. In light of this, we compared the luminescence signals from mycobacterial cells treated with either OH-CBT or its amino analogue, 6-amino-2-cyanobenzothiazole (NH_2_-CBT) (**Fig. 3d**). Notably, these compounds are known to have differences in their reaction rate constants with L-cys, with OH-CBT reacting marginally faster.^32^ Consistent with our observations in *E. coli*, NH_2_-CBT yielded a higher luminescence signal in *Msm (luc)* cells. Given the faster chemical reactivity of OH-CBT, we propose that the increased luminescence observed in cells treated with NH_2_-CBT stems from differences in the intracellular accumulation and retention of these compounds within mycobacterial cells. For subsequent assays, NH_2_-CBT was used due to its higher luminescence signal. To establish the requirement for a reduction step in signal generation, we conducted a cell-free assay to better control and define the system’s redox conditions. Specifically, we analyzed NH_2_-CBT in a cell-free setup containing only the luciferase enzyme (**Fig. S3**). Luminescence was observed only when both D-cystine and the reducing agent TCEP were present, underscoring the necessity of a reducing environment, such as that found in the cytosol, for the reaction to proceed.

We next sought to determine the optimal concentrations of assay components for use in our experiments. To this end, cells were titrated with varying concentrations of D-cystine while maintaining a constant concentration of NH_2_-CBT (**Fig. 3e**). The data revealed a clear concentration-dependent increase in luminescence. Based on the robust signal-to-noise ratios observed, we selected 50 µM as the working concentration of D-cystine for all subsequent assays. We then held D-cystine constant at 50 µM and performed a dose-response analysis of NH_2_-CBT (**Fig. 3f**). This revealed that a labeling concentration of 50 µM NH_2_-CBT provided optimal signal-to-noise levels. Accordingly, this concentration was selected for all subsequent assays. To account for background signal arising from the endogenous cysteine pool, the luminescence measured with NH_2_-CBT alone was subtracted from that of D-cys-tagged compounds hereafter, enabling a more accurate assessment of compound accumulation patterns.

Next, we confirmed that the cell envelope remained intact upon treatment with assay components. To this end, we evaluated whether incubation with NH_2_-CBT and D-cystine affected the integrity of the *Msm* cell envelope, as assessed using Nile red, a dye previously employed to monitor mycomembrane disruption.^33^ We observed that, in our assay, treatment of cells with NH_2_-CBT and D-cystine did not significantly alter fluorescence levels, indicating that these components do not compromise cell envelope integrity and are unlikely to influence the accumulation profiles of tagged molecules (**Fig. S4**). Collectively, these experiments established the optimal conditions for analyzing D-cys-tagged compounds in luciferase-expressing mycobacteria.

### Molecular Determinants of Accumulation in Mycobacteria

Having established the working parameters for D-CLAP in mycobacteria, we next sought to broadly investigate how structural modifications to molecules or changes in cellular phenotypes (including cellular treatments) influence compound accumulation. Our first objective was to directly assess the impact of esterification on accumulation. A common prodrug strategy to mask polar functional groups, such as carboxylic acids and alcohols, involves ester formation, with the ultimate goal of enhancing biological activity in both bacterial^34^ and eukaryotic systems.^35^ Notably, mycobacteria are known to express esterases capable of hydrolyzing such modifications to generate the active form of the molecule in the cytosol.^36,37^ Consequently, prodrug strategies could potentiate anti-mycobacterial drugs containing highly polar functional groups, provided this approach improves intracellular accumulation. Although ester-based modifications of antibiotics have been evaluated for anti-mycobacterial activity, their specific impact on accumulation profiles remains uncharacterized.

To empirically determine whether masking the negative charge of carboxylic acids enhances accumulation within the mycobacterial cytosol, we employed a D-cystine derivative in which both carboxylic acids were masked as methyl esters (D-cystine-ME) and performed D-CLAP in *Msm (luc)* cells (**Fig. 4a and 4b**). While D-cystine-ME can undergo condensation with NH_2_-CBT, the carboxyl group of D-luciferin must be hydrolyzed to its free acid form for proper processing by luciferase (**Fig. S1a** and **S1b**). Therefore, we anticipated that no luciferase-based signal would be detected from the parent D-cystine-ME unless the ester groups were cleaved. Indeed, in a cell-free system, luminescence signal levels in the presence of D-cystine-ME were substantially lower than those observed with the parent D-cystine (**Fig. 4a**). Moreover, luminescence signals were at background levels in the absence of a reducing agent, underscoring the requirement for disulfide bond reduction to liberate free thiols for condensation with NH_2_-CBT. Extending these findings to whole cells, mycobacterial cultures incubated with D-cystine-ME exhibited a marked increase in apparent accumulation over time compared to the carboxylic acid-based D-cystine (**Fig. 4b**). Because D-luciferin methyl ester must be hydrolyzed before luciferase can process it, this readout reports the combination of cytosolic entry and intracellular ester hydrolysis rather than entry alone. Notably, the cell-free signal for D-cystine-ME was lower than that for D-cystine, so the increase measured in whole cells is a conservative estimate of the gain in accumulation. To the best of our knowledge, these results provide the first empirical evidence that masking carboxylic acids as ester groups can specifically enhance molecular accumulation in a manner independent of anti-mycobacterial activity. We are currently leveraging D-CLAP to further investigate the substrate specificity of mycobacterial esterases.

**Figure 4.**
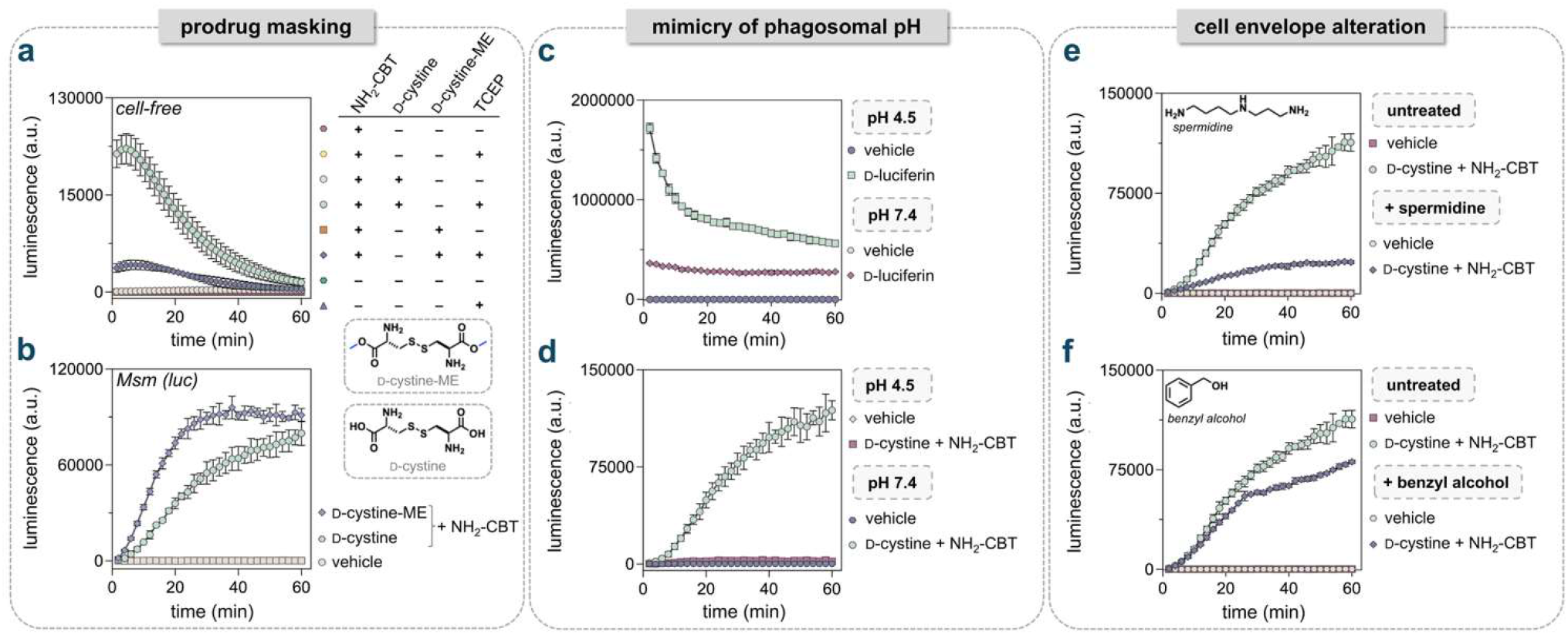
**(a)** Cell-free luminescence of luciferase in the presence or absence of NH_2_-CBT, D-cystine, D-cystine-ME, and TCEP (1 mM). **(b)** Luminescence of *Msm* (luc) cells after addition of NH_2_-CBT and either D-cystine or D-cystine-ME. **(c – d)** Luminescence of *Msm (luc)* cells in acidic (pH 4.5) and neutral (pH 7.4) buffers after addition of **(c)** D-luciferin (25 μM) or **(d)** D-cystine and NH_2_-CBT. **(e – f)** Luminescence of *Msm (luc)* cells after the addition of D-cystine and NH_2_-CBT with and without **(e)** spermidine pretreatment (10 mM, 10 min) or **(f)** benzyl alcohol pretreatment (25 mM, 60 min). Luminescence was measured over 60 min after addition of the indicated compounds (all were at 50 μM unless mentioned otherwise). Data are represented as mean ± SD (*n* = 3).

### Cellular Determinants of Accumulation in Mycobacteria

Next, we sought to broadly assess how perturbations to the cellular environment influence compound uptake. Mycobacteria are intracellular pathogens that survive and replicate within macrophage phagosomes of the host.^38^ The pH of these compartments generally ranges from 4.5 to 6.2, depending on the state of macrophage activation.^39^ The unique architecture of the mycobacterial cell wall is intrinsically linked to its resistance to acidic conditions.^39,40^ Furthermore, mycobacteria have evolved mechanisms to sense acid stress and activate metabolic responses that maintain intracellular pH homeostasis,^41,42^ disruption of which has been explored as an antimycobacterial strategy.^43^ We hypothesized that changes in extracellular pH could influence apparent accumulation by altering the physiological state of the bacterium, including permeability barriers and transport mechanisms. Such changes may contribute to mycobacterial cells restricting the uptake of antimycobacterial agents upon their entry into the phagosome.

To test the impact of acidic pH, we employed D-CLAP in *Msm (luc)* cells under conditions mimicking those of macrophage phagosomes to assess the influence of acidic pH on compound accumulation. Treatment with D-luciferin resulted in a pronounced increase in luminescence signal when the pH was lowered to mimic phagosomal conditions (**Fig. 4c**). This finding was surprising because it runs counter to the expected effect of pH on luciferase activity. Previous reports indicate that the quantum yield of firefly luciferase with D-luciferin generally decreases as pH is lowered; indeed, luminescence has been shown to be quenched around pH 5.6.^44^ Nonetheless, the intracellular pH of mycobacteria is expected to remain relatively constant despite changes in extracellular acidity.^42,45^

Next, we tested how pH changes affect signals in a split-luciferin context using D-cystine. In contrast to D-luciferin, we observed a complete loss of signal in the NH_2_-CBT and D-cystine system when the pH was lowered to mimic phagosomal conditions (**Fig. 4d**). Neither compound is expected to undergo degradation under the pH conditions employed, and our results with the parent D-luciferin confirm that luciferase expression and processing remain functional at lower pH levels. This suggests that the accumulation of certain compounds may be particularly sensitive to changes in the extracellular pH of mycobacterial cells. While some mechanistic aspects remain unresolved, we provide the first direct measurement of the effect of phagosomal-like pH on the apparent accumulation of molecules in mycobacteria.

Previous studies have suggested that polyamine treatment can modulate compound entry through porins in both Gram-negative bacteria^46^ and mycobacteria.^47,48^ In *E. coli*, this effect is thought to result from polyamines altering porin channel dynamics, shifting them toward a less permeable, closed configuration.^49^ To assess the disruption of porin-mediated entry with polyamines, we pretreated *Msm (luc)* cells with spermidine, as previously described.^48^ Optical density measurements taken before and after spermidine treatment confirmed that pretreatment did not affect cell viability, consistent with previous observations in another mycobacterial species.^48^ Interestingly, we observed a significant loss of signal upon incubation with D-luciferin when cells were pretreated with spermidine (**Fig. S5**). A similar effect was seen with D-cystine (**Fig. 4e**), consistent with a porin-mediated entry mechanism for these molecules.

These findings suggest that uptake of D-luciferin and D-cystine may be porin-mediated under the conditions tested. However, this interpretation carries a caveat, because spermidine is incompletely characterized as a porin inhibitor and its effects in this context may extend beyond porin inhibition. For example, spermidine has been reported to influence intracellular redox homeostasis in *E. coli.*^50^ Alteration of the cellular redox state could influence both the reduction of D-cystine and the oxidation of D-luciferin, which ultimately produces a luminescence signal.

There has been growing interest in identifying small-molecule adjuvants that can broadly permeabilize the mycobacterial cell envelope to enhance the uptake of potential antimycobacterial drugs. Such agents aim to overcome the formidable barrier posed by the complex mycobacterial cell wall, thereby improving drug access to intracellular targets. Among these, benzyl alcohol has received particular attention as a candidate that can disrupt envelope integrity and facilitate compound entry.^51,52^ Cells were pretreated with benzyl alcohol as previously described^51^ and the apparent uptake of various molecules was assessed using D-CLAP. As before, optical density measurements remained unchanged following benzyl alcohol exposure, indicating no impact on cell viability. Interestingly, our results showed that benzyl alcohol modestly reduced the accumulation of both D-luciferin (**Fig. S6**) and D-cystine/NH_2_-CBT over time (**Fig. 4f**). This observation was unexpected, as membrane fluidization would typically be anticipated to facilitate compound entry and thereby increase signal. Notably, we previously observed a similar phenomenon upon treatment with polymyxin B nonapeptide (PMBN) in *E. coli*, which, although typically expected to improve accumulation,^10,14^ led to lower luminescence signals, which we hypothesize may result from leakage of other components integral to our assay.^11^

Efflux pumps play a critical role in modulating drug accumulation and retention by actively exporting compounds out of the cell, thereby reducing their intracellular concentration and effectiveness.^53,54^ To probe their impact on compound accumulation, small-molecule efflux pump inhibitors can be employed, providing a strategy to assess the contribution of active efflux to uptake and retention.^55^ We set out to probe the effect of verapamil, which has been described as a modulator of efflux pump activity in mycobacterial cells.^56,57^ Cells were pretreated with verapamil as described earlier,^58^ and optical density measurements confirmed that treatment did not affect cell viability. However, verapamil treatment produced no difference in signal (**Fig. S7**). One explanation is that D-luciferin is not a substrate for the efflux pumps modulated by verapamil.

### Measuring Uptake of Polyarginine Peptides into *Mycobacteria*

One strategy for delivering cargo across cellular membranes and into the cytosol, extensively explored in mammalian systems, involves the use of cell-penetrating peptides (CPPs).^59,60^ A notable class of CPPs is the polyarginines, short polycationic peptides, whose guanidinium-rich side chains have been proposed to facilitate strong electrostatic interactions with negatively charged cell-surface components such as phosphates, promoting efficient translocation across cellular membranes.^61^ The Wender group has demonstrated that conjugating vancomycin to even a single arginine residue significantly enhances its bactericidal activity against mycobacterial cells.^62^ In a similar context, the presence of arginine residues has been shown to be essential for the antimicrobial activity of lactoferricin-derived peptides against certain mycobacterial species.^63^ While some studies have investigated the use of CPPs for delivery into bacterial cells, this remains a relatively underexplored area. Most notably, previous detection methods have primarily relied on whole-cell association – typically assessed via fluorescence microscopy – to infer uptake. However, these approaches lack direct empirical evidence confirming cytosolic localization of the peptides.

To address this gap, we empirically evaluated the uptake of a series of polyarginine peptides into the cytosol of mycobacteria using D-CLAP in *Msm (luc)* cells. For polyarginine-based CPPs, previous studies have demonstrated that the number of arginine residues is a critical determinant of cellular accumulation and association levels.^64,65^ To align with these structural patterns, we synthesized a series of polyarginine peptides (cysR5, cysR7, cysR9, and cysR11), each modified with an *N*-terminal D-cys moiety and a *C*-terminal tryptophan residue to facilitate peptide quantification (**Fig. 5a**). To evaluate potential differences in the release of the D-cys tag, we first tested our polyarginine peptide series in a cell-free system under reducing conditions. As anticipated, no significant differences in luminescence signals were observed among the polyarginine peptides, suggesting that each peptide releases an equivalent amount of D-cys under reducing conditions, in the absence of barriers to intracellular accumulation (**Fig. 5b**). We then evaluated our polyarginine series using D-CLAP by treating *Msm (luc)* cells with each individual peptide and monitoring the luminescence output (**Fig. 5c and S8**). While all four peptides demonstrated noticeable apparent accumulation over the baseline, the accumulation pattern of the peptides differed markedly from that of D-cystine, displaying an initial burst-phase uptake followed by a plateau. Among the series, cysR7 displayed the highest level of accumulation, which was also found to be concentration-dependent (**Fig. S9**). The addition of polyarginine peptides to *Msm (luc)* was also found to have minimal impact on cell envelope integrity, as determined using a Nile red assay (**Fig. S10**). Polyarginine uptake is generally thought to be dependent on the number of guanidinium groups. The accumulation profiles observed in *Msm* deviated from this trend.

**Figure 5.**
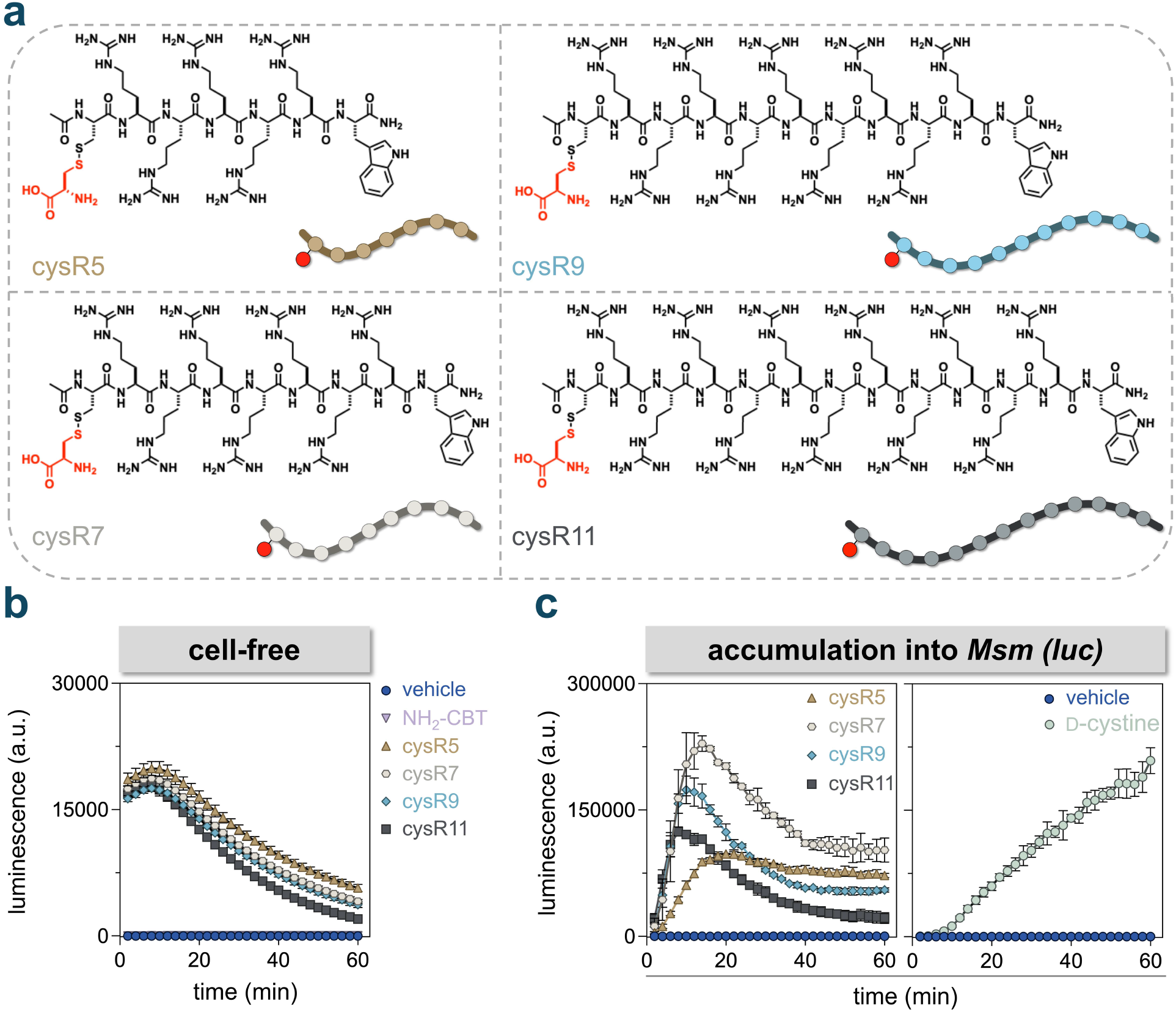
**(a)** Structures of D-cys-tagged polyarginine peptides. **(b)** Cell-free luminescence of luciferase after the addition of the indicated D-cys-tagged polyarginine peptide. **(c)** Luminescence of *Msm (luc)* cells after the addition of either D-cystine or the indicated D-cys-tagged polyarginine peptide. Luminescence was measured over 60 min after addition of the indicated compounds (all were at 50 μM unless mentioned otherwise). Data are represented as mean ± SD (*n* = 3).

Previous work,^66^ including our own,^65^ has demonstrated that the stereochemical configuration of polyarginine residues can play a critical role in determining peptide accumulation in mammalian cells. This effect may stem from chirality-dependent interactions at the lipid bilayer-solvent interface or other less well-defined interactions at the cell surface.^67^ Additionally, D-amino acids exhibit enhanced proteolytic stability^68^ which may be advantageous for certain applications. To investigate the stereochemical influence on apparent accumulation in mycobacterial cells, we synthesized and evaluated a diastereomer of cysR7 (cysr7, in which lowercase denotes D-arginine), inverting the stereochemistry at each arginine residue, in *Msm (luc)* (**Fig. S11a**). While overall accumulation levels did not differ significantly between the two peptides, their entry dynamics varied: cysr7 exhibited a more rapid decline in intracellular signal compared to cysR7 (**Fig. S11b**). These findings suggest that polyarginine uptake in mycobacteria may be largely independent of stereochemical configuration.

Mycobacteria are known to persist within the phagosomal compartment of host macrophages, presenting a formidable barrier to antimicrobial delivery.^38^ Before reaching the mycobacterial cell envelope, drugs must traverse two additional lipid bilayers, the macrophage plasma membrane and the phagosomal membrane, making intracellular mycobacteria substantially more challenging to target than extracellular ones. Building upon our observations here that polyarginine peptides accumulate within *Msm*, we sought to determine whether such peptides also associate with intracellular *Msm* residing in macrophages.

To analyze phagocytosis of mycobacteria by macrophages, we utilized a fluorescein-modified synthetic tetrapeptide PG analog developed in our lab (referred to as **PG-label** hereon), which incorporates into the PG *via* L,D-transpeptidase (Ldt)-mediated crosslinking.^69^ Briefly, *Msm (wt)* were labeled with 25 µM **PG-label** and incubated with J774A.1 macrophages at a multiplicity of infection (MOI) of 100 to promote uptake (**Fig. S12a**). Flow cytometry revealed a marked increase in fluorescence in macrophages infected with PG-labeled *Msm* compared to those infected with unlabeled *Msm*, with events shifting to the right (**Fig. S12b**). Confocal microscopy further revealed that bacteria were taken up by the phagocytic cells, and the fluorescence signal was stable (**Fig. S12c**).

Having benchmarked the infection of macrophages with *Msm*, we also evaluated the association of a fluorescent polyarginine peptide, coumR7, bearing an *N*-terminal coumarin moiety, with intracellular *Msm*. Following phagocytosis of PG-labeled *Msm*, macrophages were treated with 10 µM coumR7 for 30 min (**Fig. S12a**). Quadrant-based flow cytometry analysis demonstrated increased fluorescence in coumR7-treated cells relative to untreated controls, evident as an upward shift of events (**Fig. S12b**). In particular, for both uninfected macrophages (**Fig. S13**) and those infected with unlabeled *Msm* (**Fig. S12b**), events shifted from the bottom-left (fluorescein^-^/coumarin^-^) to the top-left quadrant (fluorescein^-^/coumarin^+^) (**Fig. S12b**). Most notably, macrophages infected with PG-labeled *Msm* exhibited a similar upward shift from the bottom-right (fluorescein^+^/coumarin^-^) to the top-right double-positive quadrant, which contained 84% of events. It is important to note that even in the absence of coumR7, macrophages harboring mycobacteria displayed a slight upward shift toward the coumarin-positive quadrants relative to macrophages alone. This is consistent with reports of *Msm* autofluorescence due to its naturally fluorescent coenzyme F_420_, which emits blue-green light near 475 nm when excited at 405 nm.^70^ Confocal imaging further suggested that the coumarin signal from the CPP colocalized primarily with intracellular mycobacteria, with minimal signal detected in the macrophage cytosol, suggesting preferential association of coumR7 with internalized bacteria (**Fig. S12c**). Collectively, our data on CPPs provide direct evidence that polyarginine-based CPPs can access the cytosol of mycobacterial cells and associate with mycobacteria residing within the phagosomes of macrophages. Together, these findings indicate that the mycobacterial envelope is more permissive to guanidinium-rich transporters than is generally assumed, and they establish a foundation for intracellular delivery strategies in these organisms. Based on these findings, we are currently pursuing the synthesis and functional evaluation of clinically relevant antibiotic conjugates incorporating releasable polyarginine-based transporters to potentially enhance their cellular permeability.

### Measuring Uptake of Antibiotics into *Mycobacteria*

We next evaluated a panel of small-molecule antibiotics with diverse physicochemical properties for their accumulation profiles in mycobacteria as a proof-of-principle. For each compound, a D-cys tag was added to make it compatible with D-CLAP. In total, we synthesized analogs of ciprofloxacin (cys-Cipro), puromycin (cys-Puro), linezolid (cys-Line), and rifamycin B (cys-Rifa) (**Fig. 6a**).^11^ First, we evaluated the antibiotic conjugates cys-Puro, cys-Rifa, cys-Cipro, and cys-Line in a cell-free system. Here, in principle, they should generate equivalent luminescence signals at the same concentration, owing to the release of an identical number of D-cys moieties.^11^ In this cell-free setup, cys-Puro, cys-Rifa, and cys-Line, all generated luminescence signals of comparable magnitude, whereas cys-Cipro produced a markedly reduced signal (**Fig. S14**). We attribute this to residual D-cys that remains covalently attached to the ciprofloxacin conjugate after disulfide bond cleavage, which could undergo a nonproductive condensation with NH_2_-CBT.^11^

**Figure 6.**
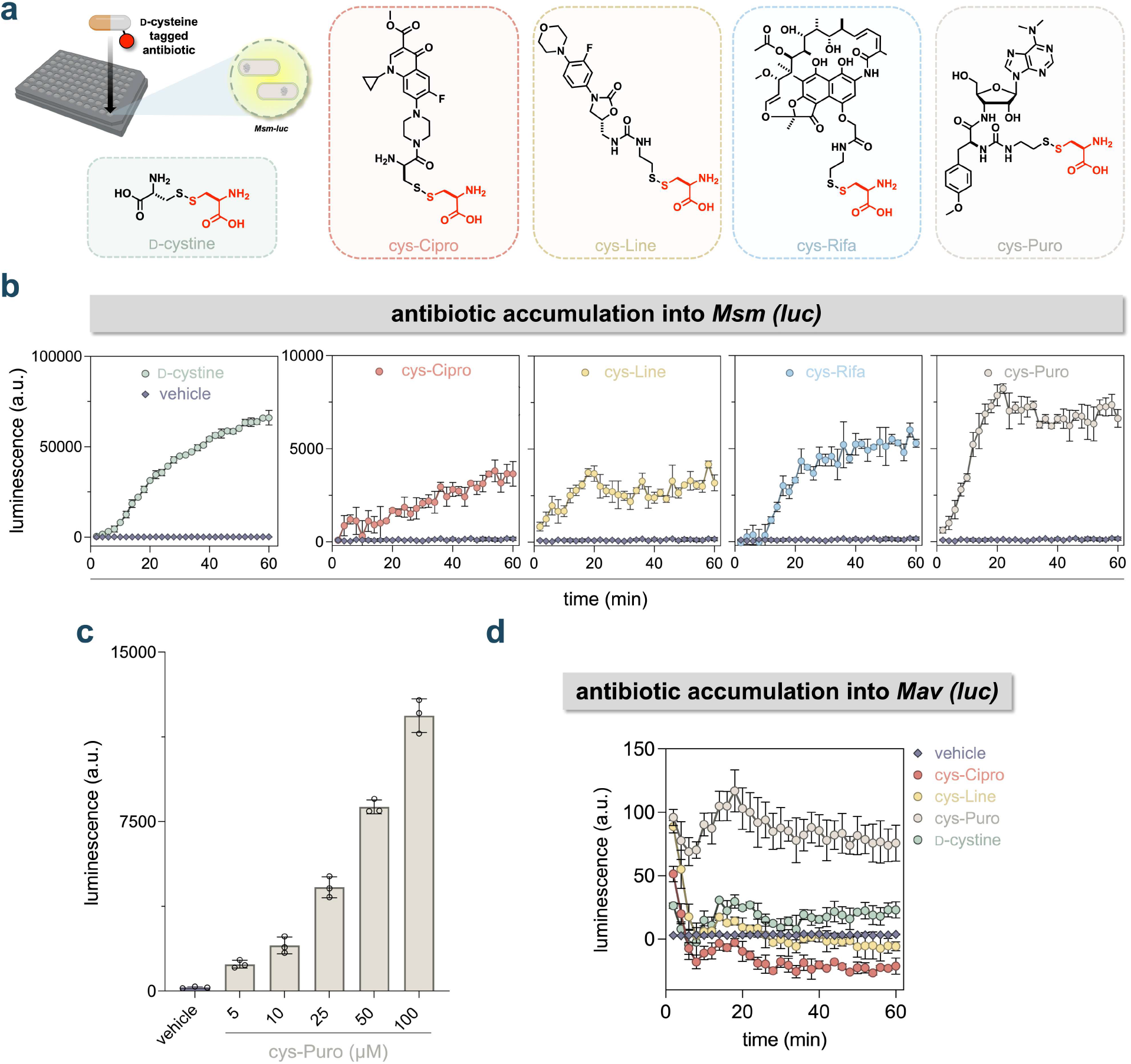
**(a)** Structures of D-cys-tagged antibiotics. **(b)** Luminescence of *Msm (luc)* cells after the addition of either D-cystine or the indicated D-cys-tagged antibiotic measured over 60 minutes. **(c)** Luminescence of *Msm (luc)* cells 60 minutes after the addition of cys-Puro (varying concentrations). **(d)** Luminescence of *Mav (luc)* after the addition of either D-cystine or the indicated D-cys-tagged antibiotic measured over 60 minutes. All assays except those with D-luciferin were done in the presence of NH_2_-CBT (50 μM). All compounds were at 50 μM unless mentioned otherwise. Data are represented as mean ± SD (*n* = 3).

With these results in hand, we next turned our attention to analyzing compound accumulation in live cells. All four antibiotic conjugates produced luminescence signals above background in D-CLAP, indicating measurable intracellular accumulation (**Fig. 6b**). For each antibiotic, the luminescence signal was approximately ten-fold lower than that observed with D-cystine, consistent with the expectation that these larger molecules would exhibit reduced apparent accumulation. At the 60-minute time point, the accumulation levels of cys-Cipro and cys-Line were comparable, whereas cys-Rifa exhibited slightly higher accumulation, and cys-Puro showed the highest accumulation among the tested conjugates (**Fig. S15**). We further analyzed the uptake of the highest accumulator in our panel, cys-Puro, and found its accumulation to be concentration-dependent (**Fig. 6c**). Additionally, treatment with the antibiotic conjugates at the tested concentrations did not perturb *Msm (luc)* cells, as assessed by the Nile red assay (**Fig. S16**).

We next sought to extend our assay to a pathogenic NTM species, *Mycobacterium avium* (*Mav*). *Mav* belongs to the *Mycobacterium avium* complex, a group of NTM species that can cause chronic pulmonary infections. To enable our assay, we engineered these cells to express luciferase, generating *Mav (luc)*. We observed a robust signal above background with D-luciferin (**Fig. S17**). This encouraged us to evaluate our panel of antibiotic conjugates in *Mav (luc)*.

As observed with *Msm (luc)*, cys-Puro exhibited the highest apparent accumulation. However, overall signal intensities in *Mav (luc)* were substantially lower relative to background, including for D-cystine (**Fig. 6d and S18**). This limits our ability to assess diverse molecules in *Mav* (luc), as evidenced by certain antibiotic conjugates exhibiting negative luminescence values after subtraction of the NH_2_-CBT-only background. Therefore, with the aim of improving signal intensities, we hypothesized that masking the two negatively charged carboxylate groups of D-cystine might enhance cellular entry. Accordingly, we evaluated D-cystine-ME and observed a marked increase in signal over D-cystine, supporting its utility as an alternative tag. This observation further demonstrates that prodrug-ester-based approaches that mask negative charge can be helpful in improving accumulation into the cytoplasm of a pathogenic *Mav* species (**Fig. S19**).

To empirically test the use of D-cystine-ME as a tag and motivated by our ability to directly observe, for the first time, antibiotic accumulation into the cytoplasm of a pathogenic mycobacterial species, we sought to apply our assay to an antibiotic scaffold of active preclinical interest. Griselimycin is a cyclic peptide antibiotic that displays strong activity against *Msm* and *Mtb* by inhibiting the bacterial DNA sliding clamp.^71^ As its target protein is cytosolic, we wanted to assess the accumulation of griselimycin in *Msm* using D-CLAP by appending an *N*-terminal disulfide-bound D-cysteine, yielding the analog cysGM (**Fig. 7a**). The disulfide tag was added to the *N*-terminus because we have previously shown that this position tolerates modification without strongly affecting peptide activity.^72^ It should also be noted that our griselimycin scaffold replaces the native 4-methyl-proline residues at positions 2 and 5 with standard prolines (GM), since the 4-methyl-proline residues are synthetically demanding. This substitution increases the MIC, but the analog remains potent. Given our results with D-cystine, we reasoned that the accumulation of GM may be affected by the addition of the negatively charged tag, as GM is a lipophilic and uncharged molecule. To account for this, we synthesized an additional GM analog with a D-cysteine-methyl ester tag, cysMeGM, which maintains the neutrality of the molecule (**Fig. 7a**).

**Figure 7.**
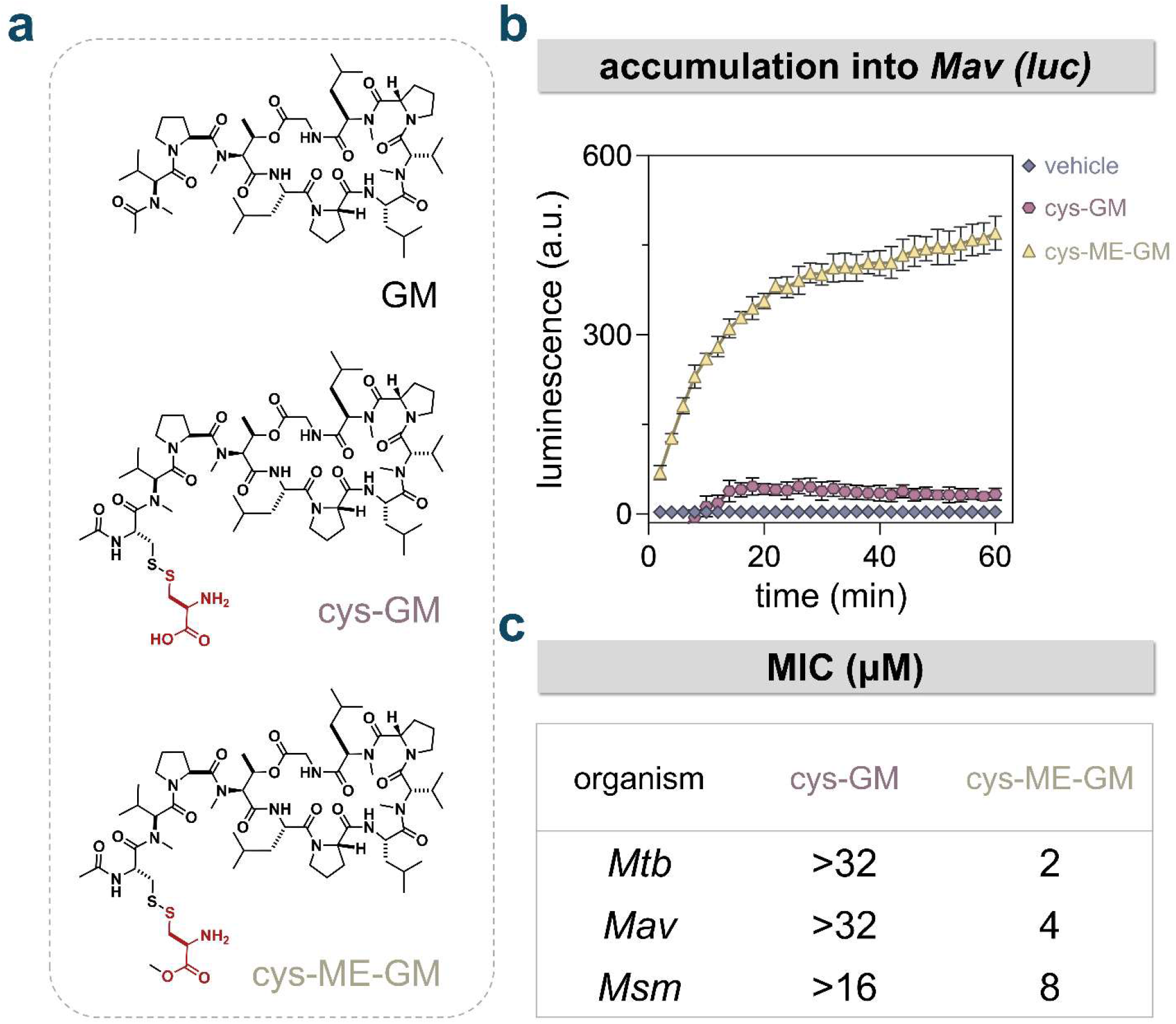
**(a)** Structures of D-cys-tagged griselimycin analogs. **(b)** Luminescence of *Mav (luc)* cells after the addition of NH_2_-CBT and either cysGM or cysMeGM measured over 60 minutes. All compounds were at 50 μM. Data are represented as mean ± SD (*n* = 3). **(c)** MIC (μM) of griselimycin derivatives against *Mtb* strain mc^2^6206, *Mav* strain MAH11, and *Msm* strain mc^2^155.

Accumulation assays in both *Msm (luc)* (**Fig. S20**) and *Mav (luc)* (**Fig. 7b**) showed substantial signal above background for cysMeGM, and as anticipated, markedly improved accumulation compared to cysGM in both species. We further evaluated the minimum inhibitory concentrations of cysGM and cysMeGM in *Msm*, *Mav,* and *Mtb* (**Fig. 7c**). In all three cases, cysMeGM displayed enhanced antimicrobial activity relative to cysGM, which could be related to its ability to accumulate better into the mycobacterial cytosol. Furthermore, cysMeGM (MIC 2 µM for *Mtb* and 8 µM for *Msm*) retains activity comparable to the parent scaffold containing standard prolines (1 µM for *Mtb* and 4 µM for *Msm*), indicating that incorporation of the tag minimally perturbs the molecule’s antibacterial activity. These results underscore the utility of D-cysteine-ME as a tagging strategy in our assay, particularly for net-neutral compounds. More broadly, they highlight the sensitivity of our platform to subtle structural modifications in antibiotic scaffolds under preclinical development. Such an approach could be leveraged for structure-activity relationship (SAR) studies of molecules like griselimycin, guiding the optimization of both antibacterial activity and pharmacokinetic properties.

## DISCUSSION

While antibiotic discovery efforts typically focus on optimizing chemical features that maximize potency against the intended target, less attention is typically paid to the compound’s ability to reach that target, a factor that can be critical when the target resides within the cytosol. This imbalance may in part stem from the lack of robust and accessible tools to quantitatively assess intracellular accumulation in mycobacteria.

In this work, we present an approach to directly assess compound accumulation within the cytosol of mycobacteria, addressing a long-standing gap in the field. The strength of this method lies in its ability to distinguish true cytosolic accumulation from general cell-associated signal, given that our assay relies on a reducing environment that is largely confined to the cytosol. In contrast, a limitation common to LC-MS/MS-based and fluorescence-based assays is that such measurements are likely to report total cellular association rather than compartment-specific entry. Moreover, the real-time luminescent readout of this approach enables time-resolved analyses, offering insight into the kinetics of intracellular uptake. The value of such dynamic monitoring is highlighted by our observations with D-cys-tagged polyarginine peptides, where accumulation patterns were nonlinear over time and would have been missed by endpoint-based measurements. Although we did not exploit this real-time kinetic capability of D-CLAP in the present study, such analyses may prove valuable for applications where the rate of uptake rather than its endpoint magnitude is informative.

However, certain limitations should be considered. First, the method relies on genetic modification, which may restrict its applicability to bacterial strains for which robust genetic tools are available. Second, the tag adds molecular bulk, which could affect how the compound is taken up or distributed within the cell. The tag may also change physicochemical properties and potentially influence target engagement of small-molecule antibiotics. For these reasons, the functionalized compounds may not behave exactly like the parent antibiotics in terms of antibacterial activity. Third, the assay cannot be extended to intracellular bacteria as the cytosolic milieu of the macrophage is reducing, which could lead to premature cleavage of the disulfide bond linking the reporter D-cysteine moiety to our conjugate prior to entry into the mycobacterial cytosol. In such a scenario, the luciferase signal could arise from free D-cysteine entering the cytoplasm of mycobacteria rather than the intact conjugate, thereby confounding the interpretation of cytoplasmic delivery.

Nonetheless, our approach yielded several notable insights. The applications presented here were chosen to span chemically and mechanistically distinct permeants, including small-molecule antibiotics, cationic peptides, and a cyclic peptide natural product, in order to define the operating range of the assay rather than to dissect any single uptake pathway exhaustively. For example, we were able to directly test whether perturbations to the mycobacterium or its environment that were previously reported to influence compound accumulation actually had the expected effect, providing a robust means to validate prior assumptions. We assessed the entry of a chemically diverse panel of antibiotic conjugates into mycobacteria, including an NTM species, revealing key differences in uptake. We were also able to examine the cytosolic entry of a preclinical antibiotic scaffold, GM, revealing the effect of subtle structural differences on its uptake. The ability to capture such effects could be crucial for SAR-based studies of molecules in this class. Additionally, expanding such assessments to a larger set of compounds could ultimately help define a “chemical grammar” for mycobacterial accumulation, guiding the rational design of future therapeutics against NTM species. We also used our method to examine the uptake of D-cys-tagged polyarginine peptides into mycobacteria and, to our knowledge, provide the first evidence that these peptides associate with mycobacterial cells residing within macrophages.

## ACKNOWLEDGMENTS

This study was supported by NIH grants 1R01AI178975-01 (M.M.P.), R35GM124893 (M.M.P.), and R01AI179080-01 (M.M.P.). We thank the director of the University of Massachusetts Amherst Biophysical Characterization facilities, Dr. Stephen Eyles, for his help and advice, and the Massachusetts Life Sciences Center for the funding for the equipment.

## SUPPORTING INFORMATION

Luciferin oxidation schemes and growth curves; cell-free luciferase and Nile Red membrane integrity controls; accumulation assays following spermidine, benzyl alcohol, or verapamil pretreatment; accumulation of d-cys-tagged polyarginine conjugates and the all-d diastereomer, with flow cytometry and confocal analysis of coumR7-treated macrophages; accumulation of d-cys-tagged antibiotic conjugates in *Msm* (luc) and *Mav* (luc); methyl ester controls; materials, biological methods, and synthesis and characterization of all peptide and antibiotic conjugates.

## REFERENCES

(1) Dahl, V. N.; Mølhave, M.; Fløe, A.; van Ingen, J.; Schön, T.; Lillebaek, T.; Andersen, A. B.; Wejse, C. Global Trends of Pulmonary Infections with Nontuberculous Mycobacteria: A Systematic Review. Int. J. Infect. Dis. 2022, 125, 120–131. 10.1016/j.ijid.2022.10.013

(2) Ratnatunga, C. N.; Lutzky, V. P.; Kupz, A.; Doolan, D. L.; Reid, D. W.; Field, M.; Bell, S. C.; Thomson, R. M.; Miles, J. J. The Rise of Non-Tuberculosis Mycobacterial Lung Disease. Front. Immunol. 2020, 11. 10.3389/fimmu.2020.00303

(3) Johansen, M. D.; Herrmann, J.-L.; Kremer, L. Non-Tuberculous Mycobacteria and the Rise of Mycobacterium Abscessus. Nat. Rev. Microbiol. 2020, 18 (7), 392–407. 10.1038/s41579-020-0331-1

(4) Dulberger, C. L.; Rubin, E. J.; Boutte, C. C. The Mycobacterial Cell Envelope — a Moving Target. Nat. Rev. Microbiol. 2020, 18 (1), 47–59. 10.1038/s41579-019-0273-7

(5) Bansal-Mutalik, R.; Nikaido, H. Mycobacterial Outer Membrane Is a Lipid Bilayer and the Inner Membrane Is Unusually Rich in Diacyl Phosphatidylinositol Dimannosides. Proc. Natl. Acad. Sci. 2014, 111 (13), 4958–4963. 10.1073/pnas.1403078111

(6) Chiaradia, L.; Lefebvre, C.; Parra, J.; Marcoux, J.; Burlet-Schiltz, O.; Etienne, G.; Tropis, M.; Daffé, M. Dissecting the Mycobacterial Cell Envelope and Defining the Composition of the Native Mycomembrane. Sci. Rep. 2017, 7 (1), 12807. 10.1038/s41598-017-12718-4

(7) Mateus, A.; Matsson, P.; Artursson, P. Rapid Measurement of Intracellular Unbound Drug Concentrations. Mol. Pharm. 2013, 10 (6), 2467–2478. 10.1021/mp4000822

(8) Overington, J. P.; Al-Lazikani, B.; Hopkins, A. L. How Many Drug Targets Are There? Nat. Rev. Drug Discov. 2006, 5 (12), 993–996. 10.1038/nrd2199

(9) Kelly, J. J.; Dalesandro, B. E.; Liu, Z.; Chordia, M. D.; Ongwae, G. M.; Pires, M. M. Measurement of Accumulation of Antibiotics to Staphylococcus Aureus in Phagosomes of Live Macrophages. Angew. Chem. Int. Ed. 2024, 63 (3), e202313870. 10.1002/anie.202313870

(10) Ongwae, G. M.; Lepori, I.; Chordia, M. D.; Dalesandro, B. E.; Apostolos, A. J.; Siegrist, M. S.; Pires, M. M. Measurement of Small Molecule Accumulation into Diderm Bacteria. ACS Infect. Dis. 2023, 9 (1), 97–110. 10.1021/acsinfecdis.2c00435

(11) Dash, R.; Holsinger, K. A.; Chordia, M. D.; Gh M. S.; Pires, M. M. Bioluminescence-Based Determination of Cytosolic Accumulation of Antibiotics in Escherichia Coli. ACS Infect. Dis. 2024, 10 (5), 1602–1611. 10.1021/acsinfecdis.3c00684

(12) Lepori, I.; Jackson, K.; Liu, Z.; Chordia, M. D.; Wong, M.; Rivera, S. L.; Roncetti, M.; Poliseno, L.; Freundlich, J. S.; Pires, M. M.; Siegrist, M. S. The Mycomembrane Differentially and Heterogeneously Restricts Antibiotic Permeation. ACS Infect. Dis. 2025, 11 (7), 1893–1906. 10.1021/acsinfecdis.4c01062

(13) Lepori, I.; Liu, Z.; Evbarunegbe, N.; Feng, S.; Brown, T. P.; Mane, K.; Dash, R.; Shivangi Wong, M.; Naick, A.; George, A.; Guo, T.; Shelke, A. M.; Sharpless, K. B.; Dong, J.; Freundlich, J. S.; Im, W.; Green, A. G.; Pires, M. M.; Siegrist, M. S. Identification of Chemical Features for Improved Outer Membrane Permeation in Mycobacteria Using Machine Learning. Nat. Microbiol. 2026, 1–19. 10.1038/s41564-026-02412-5

(14) Ongwae, G. M.; Liu, Z.; Feng, S.; Chordia, M. D.; Dash, R.; He, Y.; Gh, M. S.; Dalesandro, B. E.; Guo, T.; Sharpless, K. B.; Dong, J.; Sloan Siegrist, M.; Im, W.; Pires, M. M. Click-Based Determination of Accumulation of Molecules in Escherichia Coli. Nat. Commun. 2026, 17 (1), 2008. 10.1038/s41467-026-68717-5

(15) Dash, R.; Liu, Z.; Lepori, I.; Chordia, M. D.; Ocius, K.; Holsinger, K.; Zhang, H.; Kenyon, R.; Im, W.; Siegrist, M. S.; Pires, M. M. Systematic Determination of the Impact of Structural Edits on Peptide Accumulation into Mycobacteria. ACS Chem. Biol. 2025, 20 (8), 1962–1979. 10.1021/acschembio.5c00330

(16) Liu, Z.; Lepori, I.; Chordia, M. D.; Dalesandro, B. E.; Guo, T.; Dong, J.; Siegrist, M. S.; Pires, M. M. A Metabolic-Tag-Based Method for Assessing the Permeation of Small Molecules Across the Mycomembrane in Live Mycobacteria. Angew. Chem. Int. Ed. 2023, 62 (20), e202217777. 10.1002/anie.202217777

(17) Geddes, E. J.; Li, Z.; Hergenrother, P. J. An LC-MS/MS Assay and Complementary Web-Based Tool to Quantify and Predict Compound Accumulation in E. Coli. Nat. Protoc. 2021, 16 (10), 4833–4854. 10.1038/s41596-021-00598-y

(18) Iyer, R.; Ye, Z.; Ferrari, A.; Duncan, L.; Tanudra, M. A.; Tsao, H.; Wang, T.; Gao, H.; Brummel, C. L.; Erwin, A. L. Evaluating LC–MS/MS To Measure Accumulation of Compounds within Bacteria. ACS Infect. Dis. 2018, 4 (9), 1336–1345. 10.1021/acsinfecdis.8b00083

(19) Sullivan, M. R.; Rubin, E. J. Deep Learning-Based Prediction of Chemical Accumulation in a Pathogenic Mycobacterium. bioRxiv December 16, 2024, p 2024.12.15.628588. 10.1101/2024.12.15.628588

(20) Davis, T. D.; Gerry, C. J.; Tan, D. S. General Platform for Systematic Quantitative Evaluation of Small-Molecule Permeability in Bacteria. ACS Chem. Biol. 2014, 9 (11), 2535–2544. 10.1021/cb5003015

(21) Karatas, H.; Maric, T.; D’Alessandro, P. L.; Yevtodiyenko, A.; Vorherr, T.; Hollingworth, G. J.; Goun, E. A. Real-Time Imaging and Quantification of Peptide Uptake in Vitro and in Vivo. ACS Chem. Biol. 2019, 14 (10), 2197–2205. 10.1021/acschembio.9b00439

(22) Godinat, A.; Bazhin, A. A.; Goun, E. A. Bioorthogonal Chemistry in Bioluminescence Imaging. Drug Discov. Today 2018, 23 (9), 1584–1590. 10.1016/j.drudis.2018.05.022

(23) Sparks, I. L.; Derbyshire, K. M.; Jacobs, W. R.; Morita, Y. S. Mycobacterium Smegmatis: The Vanguard of Mycobacterial Research. J. Bacteriol. 2023, 205 (1), e00337–22. 10.1128/jb.00337-22

(24) Andreu, N.; Zelmer, A.; Fletcher, T.; Elkington, P. T.; Ward, T. H.; Ripoll, J.; Parish, T.; Bancroft, G. J.; Schaible, U.; Robertson, B. D.; Wiles, S. Optimisation of Bioluminescent Reporters for Use with Mycobacteria. PLOS ONE 2010, 5 (5), e10777. 10.1371/journal.pone.0010777

(25) Newton, G. L.; Buchmeier, N.; Fahey, R. C. Biosynthesis and Functions of Mycothiol, the Unique Protective Thiol of Actinobacteria. Microbiol. Mol. Biol. Rev. MMBR 2008, 72 (3), 471–494. 10.1128/MMBR.00008-08

(26) Reyes, A. M.; Pedre, B.; De Armas, M. I.; Tossounian, M.-A.; Radi, R.; Messens, J.; Trujillo, M. Chemistry and Redox Biology of Mycothiol. Antioxid. Redox Signal. 2018, 28 (6), 487–504. 10.1089/ars.2017.7074

(27) Van de Bittner, G. C.; Bertozzi, C. R.; Chang, C. J. Strategy for Dual-Analyte Luciferin Imaging: In Vivo Bioluminescence Detection of Hydrogen Peroxide and Caspase Activity in a Murine Model of Acute Inflammation. J. Am. Chem. Soc. 2013, 135 (5), 1783–1795. 10.1021/ja309078t

(28) Nakamura, M.; Niwa, K.; Maki, S.; Hirano, T.; Ohmiya, Y.; Niwa, H. Construction of a New Firefly Bioluminescence System Using L-Luciferin as Substrate. Tetrahedron Lett. 2006, 47 (7), 1197–1200. 10.1016/j.tetlet.2005.12.033

(29) Niwa, K.; Nakajima, Y.; Ohmiya, Y. Applications of Luciferin Biosynthesis: Bioluminescence Assays for l-Cysteine and Luciferase. Anal. Biochem. 2010, 396 (2), 316–318. 10.1016/j.ab.2009.09.014

(30) Zhang, C.; Yuan, M.; Han, G.; Gao, Y.; Tang, C.; Li, X.; Du, L.; Li, M. Novel Caged Luciferin Derivatives Can Prolong Bioluminescence Imaging in Vitro and in Vivo. RSC Adv. 2018, 8 (35), 19596–19599. 10.1039/C8RA02312C

(31) Reddy, G. R.; Thompson, W. C.; Miller, S. C. Robust Light Emission from Cyclic Alkylaminoluciferin Substrates for Firefly Luciferase. J. Am. Chem. Soc. 2010, 132 (39), 13586–13587. 10.1021/ja104525m

(32) Godinat, A.; Budin, G.; Morales, A. R.; Park, H. M.; Sanman, L. E.; Bogyo, M.; Yu, A.; Stahl, A.; Dubikovskaya, E. A. A Biocompatible “Split Luciferin” Reaction and Its Application for Non-Invasive Bioluminescent Imaging of Protease Activity in Living Animals. Curr. Protoc. Chem. Biol. 2014, 6 (3), 169–189. 10.1002/9780470559277.ch140047

(33) Campodónico, V. L.; Rifat, D.; Chuang, Y.-M.; Ioerger, T. R.; Karakousis, P. C. Altered Mycobacterium Tuberculosis Cell Wall Metabolism and Physiology Associated With RpoB Mutation H526D. Front. Microbiol. 2018, 9. 10.3389/fmicb.2018.00494

(34) Larsen, E. M.; Johnson, R. J. Microbial Esterases and Ester Prodrugs: An Unlikely Marriage for Combating Antibiotic Resistance. 10.1002/ddr.21468

(35) Rautio, J.; Kumpulainen, H.; Heimbach, T.; Oliyai, R.; Oh, D.; Järvinen, T.; Savolainen, J. Prodrugs: Design and Clinical Applications. Nat. Rev. Drug Discov. 2008, 7 (3), 255–270. 10.1038/nrd2468

(36) Larsen, E. M.; Stephens, D. C.; Clarke, N. H.; Johnson, R. J. Ester-Prodrugs of Ethambutol Control Its Antibacterial Activity and Provide Rapid Screening for Mycobacterial Hydrolase Activity. Bioorg. Med. Chem. Lett. 2017, 27 (19), 4544–4547. 10.1016/j.bmcl.2017.08.057

(37) Tallman, K. R.; Levine, S. R.; Beatty, K. E. Small-Molecule Probes Reveal Esterases with Persistent Activity in Dormant and Reactivating Mycobacterium Tuberculosis. ACS Infect. Dis. 2016, 2 (12), 936–944. 10.1021/acsinfecdis.6b00135

(38) Pieters, J. *Mycobacterium Tuberculosis* and the Macrophage: Maintaining a Balance. Cell Host Microbe 2008, 3 (6), 399–407. 10.1016/j.chom.2008.05.006

(39) Vandal, O. H.; Nathan, C. F.; Ehrt, S. Acid Resistance in Mycobacterium Tuberculosis. J. Bacteriol. 2009, 191 (15), 4714–4721. 10.1128/jb.00305-09

(40) Vandal, O. H.; Roberts, J. A.; Odaira, T.; Schnappinger, D.; Nathan, C. F.; Ehrt, S. Acid-Susceptible Mutants of Mycobacterium Tuberculosis Share Hypersusceptibility to Cell Wall and Oxidative Stress and to the Host Environment. J. Bacteriol. 2009. 10.1128/jb.00932-08

(41) Laudouze, J.; Canaan, S.; Gouzy, A.; Santucci, P. Unraveling Mycobacterium Tuberculosis Acid Resistance and pH Homeostasis Mechanisms. FEBS Lett. 2025, 599 (12), 1634–1648. 10.1002/1873-3468.70023

(42) Rao, M.; Streur, T. L.; Aldwell, F. E.; Cook, G. M. Intracellular pH Regulation by Mycobacterium Smegmatis and Mycobacterium Bovis BCG. Microbiology 2001, 147 (4), 1017–1024. 10.1099/00221287-147-4-1017

(43) Early, J.; Ollinger, J.; Darby, C.; Alling, T.; Mullen, S.; Casey, A.; Gold, B.; Ochoada, J.; Wiernicki, T.; Masquelin, T.; Nathan, C.; Hipskind, P. A.; Parish, T. Identification of Compounds with pH-Dependent Bactericidal Activity against Mycobacterium Tuberculosis. ACS Infect. Dis. 2019, 5 (2), 272–280. 10.1021/acsinfecdis.8b00256

(44) Ando, Y.; Niwa, K.; Yamada, N.; Enomoto, T.; Irie, T.; Kubota, H.; Ohmiya, Y.; Akiyama, H. Firefly Bioluminescence Quantum Yield and Colour Change by pH-Sensitive Green Emission. Nat. Photonics 2008, 2 (1), 44–47. 10.1038/nphoton.2007.251

(45) Vandal, O. H.; Pierini, L. M.; Schnappinger, D.; Nathan, C. F.; Ehrt, S. A Membrane Protein Preserves Intrabacterial pH in Intraphagosomal Mycobacterium Tuberculosis. Nat. Med. 2008, 14 (8), 849–854. 10.1038/nm.1795

(46) Dela Vega, A. L.; Delcour, A. H. Polyamines Decrease Escherichia Coli Outer Membrane Permeability. J. Bacteriol. 1996, 178 (13), 3715–3721. 10.1128/jb.178.13.3715-3721.1996

(47) Dai, T.; Xie, J.; Zhu, Q.; Kamariza, M.; Jiang, K.; Bertozzi, C. R.; Rao, J. A Fluorogenic Trehalose Probe for Tracking Phagocytosed Mycobacterium Tuberculosis. J. Am. Chem. Soc. 2020, 142 (36), 15259–15264. 10.1021/jacs.0c07700

(48) Sarathy, J. P.; Lee, E.; Dartois, V. Polyamines Inhibit Porin-Mediated Fluoroquinolone Uptake in Mycobacteria. PLOS ONE 2013, 8 (6), e65806. 10.1371/journal.pone.0065806

(49) Iyer, R.; Delcour, A. H. Complex Inhibition of OmpF and OmpC Bacterial Porins by Polyamines*. J. Biol. Chem. 1997, 272 (30), 18595–18601. 10.1074/jbc.272.30.18595

(50) Kumar, V.; Mishra, R. K.; Ghose, D.; Kalita, A.; Dhiman, P.; Prakash, A.; Thakur, N.; Mitra, G.; Chaudhari, V. D.; Arora, A.; Dutta, D. Free Spermidine Evokes Superoxide Radicals That Manifest Toxicity. eLife 2022, 11, e77704. 10.7554/eLife.77704

(51) Nguyen, P. P.; Kado, T.; Prithviraj, M.; Siegrist, M. S.; Morita, Y. S. Inositol Acylation of Phosphatidylinositol Mannosides: A Rapid Mass Response to Membrane Fluidization in Mycobacteria. J. Lipid Res. 2022, 63 (9), 100262. 10.1016/j.jlr.2022.100262

(52) Kado, T.; Akbary, Z.; Motooka, D.; Sparks, I. L.; Melzer, E. S.; Nakamura, S.; Rojas, E. R.; Morita, Y. S.; Siegrist, M. S. A Cell Wall Synthase Accelerates Plasma Membrane Partitioning in Mycobacteria. eLife 2023, 12, e81924. 10.7554/eLife.81924

(53) Webber, M. A.; Piddock, L. J. V. The Importance of Efflux Pumps in Bacterial Antibiotic Resistance. J. Antimicrob. Chemother. 2003, 51 (1), 9–11. 10.1093/jac/dkg050

(54) Du, D.; Wang-Kan, X.; Neuberger, A.; van Veen, H. W.; Pos, K. M.; Piddock, L. J. V.; Luisi, B. F. Multidrug Efflux Pumps: Structure, Function and Regulation. Nat. Rev. Microbiol. 2018, 16 (9), 523–539. 10.1038/s41579-018-0048-6

(55) Stavri, M.; Piddock, L. J. V.; Gibbons, S. Bacterial Efflux Pump Inhibitors from Natural Sources. J. Antimicrob. Chemother. 2007, 59 (6), 1247–1260. 10.1093/jac/dkl460

(56) Pasca, M. R.; Guglierame, P.; De Rossi, E.; Zara, F.; Riccardi, G. mmpL7 Gene of Mycobacterium Tuberculosis Is Responsible for Isoniazid Efflux in Mycobacterium Smegmatis. Antimicrob. Agents Chemother. 2005, 49 (11), 4775–4777. 10.1128/aac.49.11.4775-4777.2005

(57) Silva, P. E. A.; Bigi, F.; de la Paz Santangelo, M.; Romano, M. I.; Martín, C.; Cataldi, A.; Aínsa, J. A. Characterization of P55, a Multidrug Efflux Pump in Mycobacterium Bovis and Mycobacterium Tuberculosis. Antimicrob. Agents Chemother. 2001, 45 (3), 800–804. 10.1128/aac.45.3.800-804.2001

(58) Pasca, M. R.; Guglierame, P.; Arcesi, F.; Bellinzoni, M.; De Rossi, E.; Riccardi, G. Rv2686c-Rv2687c-Rv2688c, an ABC Fluoroquinolone Efflux Pump in Mycobacterium Tuberculosis. Antimicrob. Agents Chemother. 2004, 48 (8), 3175–3178. 10.1128/aac.48.8.3175-3178.2004

(59) Copolovici, D. M.; Langel, K.; Eriste, E.; Langel, Ü. Cell-Penetrating Peptides: Design, Synthesis, and Applications. ACS Nano 2014, 8 (3), 1972–1994. 10.1021/nn4057269

(60) Stanzl, E. G.; Trantow, B. M.; Vargas, J. R.; Wender, P. A. Fifteen Years of Cell-Penetrating, Guanidinium-Rich Molecular Transporters: Basic Science, Research Tools, and Clinical Applications. Acc. Chem. Res. 2013, 46 (12), 2944–2954. 10.1021/ar4000554

(61) Robison, A. D.; Sun, S.; Poyton, M. F.; Johnson, G. A.; Pellois, J.-P.; Jungwirth, P.; Vazdar, M.; Cremer, P. S. Polyarginine Interacts More Strongly and Cooperatively than Polylysine with Phospholipid Bilayers. J. Phys. Chem. B 2016, 120 (35), 9287–9296. 10.1021/acs.jpcb.6b05604

(62) Brčić, J.; Tong, A.; Wender, P. A.; Cegelski, L. Conjugation of Vancomycin with a Single Arginine Improves Efficacy against Mycobacteria by More Effective Peptidoglycan Targeting. J. Med. Chem. 2023, 66 (15), 10226–10237. 10.1021/acs.jmedchem.3c00565

(63) Silva, T.; Magalhães, B.; Maia, S.; Gomes, P.; Nazmi, K.; Bolscher, J. G. M.; Rodrigues, P. N.; Bastos, M.; Gomes, M. S. Killing of Mycobacterium Avium by Lactoferricin Peptides: Improved Activity of Arginine- and d-Amino-Acid-Containing Molecules. Antimicrob. Agents Chemother. 2014, 58 (6), 3461–3467. 10.1128/aac.02728-13

(64) Mitchell, D. j.; Steinman, L.; Kim, D. t.; Fathman, C. g.; Rothbard, J. b. Polyarginine Enters Cells More Efficiently than Other Polycationic Homopolymers. J. Pept. Res. 2000, 56 (5), 318–325. 10.1034/j.1399-3011.2000.00723.x

(65) Bhandari, S.; Ongwae, G. M.; Dash, R.; Liu, Z.; Chordia, M. D.; He, Y.; Pires, M. M. A Generalizable Assay for Intracellular Accumulation to Profile Cytosolic Drug Delivery in Mammalian Cells. Commun. Chem. 2026, 9 (1), 94. 10.1038/s42004-026-01898-8

(66) Verdurmen, W. P. R.; Bovee-Geurts, P. H.; Wadhwani, P.; Ulrich, A. S.; Hällbrink, M.; van Kuppevelt, T. H.; Brock, R. Preferential Uptake of L-versus D-Amino Acid Cell-Penetrating Peptides in a Cell Type-Dependent Manner. Chem. Biol. 2011, 18 (8), 1000–1010. 10.1016/j.chembiol.2011.06.006

(67) Hu, J.; Cochrane, W. G.; Jones, A. X.; Blackmond, D. G.; Paegel, B. M. Chiral Lipid Bilayers Are Enantioselectively Permeable. Nat. Chem. 2021, 13 (8), 786–791. 10.1038/s41557-021-00708-z

(68) Najjar, K.; Erazo-Oliveras, A.; Brock, D. J.; Wang, T.-Y.; Pellois, J.-P. An L-to d-Amino Acid Conversion in an Endosomolytic Analog of the Cell-Penetrating Peptide TAT Influences Proteolytic Stability, Endocytic Uptake, and Endosomal Escape*. J. Biol. Chem. 2017, 292 (3), 847–861. 10.1074/jbc.M116.759837

(69) Pidgeon, S. E.; Apostolos, A. J.; Nelson, J. M.; Shaku, M.; Rimal, B.; Islam, M. N.; Crick, D. C.; Kim, S. J.; Pavelka, M. S.; Kana, B. D.; Pires, M. M. L,D-Transpeptidase Specific Probe Reveals Spatial Activity of Peptidoglycan Cross-Linking. ACS Chem. Biol. 2019, 14 (10), 2185–2196. 10.1021/acschembio.9b00427

(70) Patiño, S.; Alamo, L.; Cimino, M.; Casart, Y.; Bartoli, F.; García, M. J.; Salazar, L. Autofluorescence of Mycobacteria as a Tool for Detection of Mycobacterium Tuberculosis. J. Clin. Microbiol. 2008, 46 (10), 3296–3302. 10.1128/JCM.02183-07

(71) Kling, A.; Lukat, P.; Almeida, D. V.; Bauer, A.; Fontaine, E.; Sordello, S.; Zaburannyi, N.; Herrmann, J.; Wenzel, S. C.; König, C.; Ammerman, N. C.; Barrio, M. B.; Borchers, K.; Bordon-Pallier, F.; Brönstrup, M.; Courtemanche, G.; Gerlitz, M.; Geslin, M.; Hammann, P.; Heinz, D. W.; Hoffmann, H.; Klieber, S.; Kohlmann, M.; Kurz, M.; Lair, C.; Matter, H.; Nuermberger, E.; Tyagi, S.; Fraisse, L.; Grosset, J. H.; Lagrange, S.; Müller, R. Antibiotics. Targeting DnaN for Tuberculosis Therapy Using Novel Griselimycins. Science 2015, 348 (6239), 1106–1112. 10.1126/science.aaa4690

(72) Spira, A.; Dash, R.; Lepori, I.; Luo, Y. C.; Newkirk, S. E.; Bhandari, S.; Siegrist, M. S.; Pires, M. M. Molecular Determinants Governing the Antitubercular Activity of Griselimycin. bioRxiv March 22, 2026, p 2026.03.19.712639. 10.64898/2026.03.19.712639

