## Supporting Information for "A Split-Luciferin Assay for Real-Time Measurement of Cytosolic Accumulation in Live Mycobacteria"

### TABLE OF CONTENTS


| | |
|---|---|
| **SUPPLEMENTARY FIGURES** | S4-S23 |
| **Figure S1.** Schematic of the luciferase-mediated oxidation of D-luciferin and D-luciferin methyl ester. | S4 |
| **Figure S2.** Optical density (O.D.) at 600 nm of *Msm (luc)* growth cultures from **Fig. 1d**. | S5 |
| **Figure S3.** Bioluminescence analysis of a cell-free assay with luciferase in the presence or absence of $NH_2$-CBT, D-cystine, and TCEP. | S6 |
| **Figure S4.** Nile Red-based mycobacterial membrane integrity assay of *Msm (luc)* cells incubated with $NH_2$-CBT, D-cystine, or D-luciferin. | S7 |
| **Figure S5.** Spermidine pretreatment accumulation assay. | S8 |
| **Figure S6.** Benzyl alcohol pretreatment accumulation assay. | S9 |
| **Figure S7.** Verapamil pretreatment accumulation assay. | S10 |
| **Figure S8.** Accumulation at the 20-minute time point of D-cys-tagged polyarginine peptide conjugates. | S11 |
| **Figure S9.** Bioluminescence analysis of *Msm (luc)* cells with varying concentrations of cysR7. | S12 |
| **Figure S10.** Nile Red-based mycobacterial membrane integrity assay of *Msm (luc)* cells incubated with D-cys tagged polyarginine peptide conjugates. | S13 |
| **Figure S11.** Accumulation of D-cys tagged polyarginine peptide diastereomers in *Msm (luc)*. | S14 |
| **Figure S12.** Infection assays of *Msm* in macrophages treated with coumR7. | S15 |
| **Figure S13.** Flow cytometry scatter plots of uninfected macrophages in the absence or presence of treatment with coumR7. | S16 |
| **Figure S14.** Bioluminescence analysis of a cell-free assay with luciferase in the presence of D-cys tagged antibiotic conjugates. | S17 |
| **Figure S15.** Endpoint accumulation of D-cys tagged antibiotic conjugates in *Msm (luc)*. | S18 |
| **Figure S16.** Nile Red-based mycobacterial membrane integrity assay of *Msm (luc)* cells incubated with D-cys tagged antibiotic conjugates. | S19 |
| **Figure S17.** Bioluminescence analysis of *Mav (luc)* cells in the presence of D-luciferin. | S20 |
| **Figure S18.** Endpoint accumulation of D-cys tagged antibiotic conjugates in *Mav (luc)*. | S21 |
| **Figure S19.** Bioluminescence analysis of *Mav (luc)* cells in the presence of $NH_2$-CBT and D-cystine or D-cystine-ME. | S22 |
| **Figure S20.** Bioluminescence analysis of *Msm (luc)* cells in the presence of $NH_2$-CBT and cys-GM or cys-ME-GM. | S23 |
| **EXPERIMENTAL PROCEDURES** | S24-S66 |
| **Materials** | S24 |
| **Biological Methods** | S27 |

| | |
|---|---|
| Preparation of electrocompetent mycobacteria cells and transformation | S27 |
| Mycobacteria cell culture | S27 |
| Mammalian cell culture | S28 |
| Luminescence imaging of multiwell plates | S28 |
| Bioluminescence-based accumulation assays | S28 |
| Cell-free luciferase assays | S29 |
| Nile red assay for assessment of mycobacterial membrane integrity | S29 |
| Flow cytometry analysis of PG-labeled *Msm* in J774A.1 macrophages treated with coumR7 | S30 |
| Confocal microscopy analysis of PG-labeled *Msm* in J774A.1 macrophages treated with coumR7 | S30 |
| MIC assays | S31 |
| **Synthesis and Characterization of Peptide Conjugates** | S32-S43 |
| Polyarginine peptide conjugates | S32 |
| Griselimycin peptide conjugates | S40 |
| **Synthesis and Characterization of Antibiotic Conjugates** | S44-S64 |
| General Methods | S44 |
| D-cystine-ciprofloxacin-methyl-ester (cys-Cipro) | S45 |
| Linezolid-cystamine- D-cysteine disulfide (cys-Line) | S52 |
| Puromycin-cystamine- D-cysteine disulfide (cys-Puro) | S58 |
| Rifamycin-cystamine- D-cysteine disulfide (cys-Rifa) | S63 |
| **REFERENCES** | S67 |

### SUPPLEMENTARY FIGURES

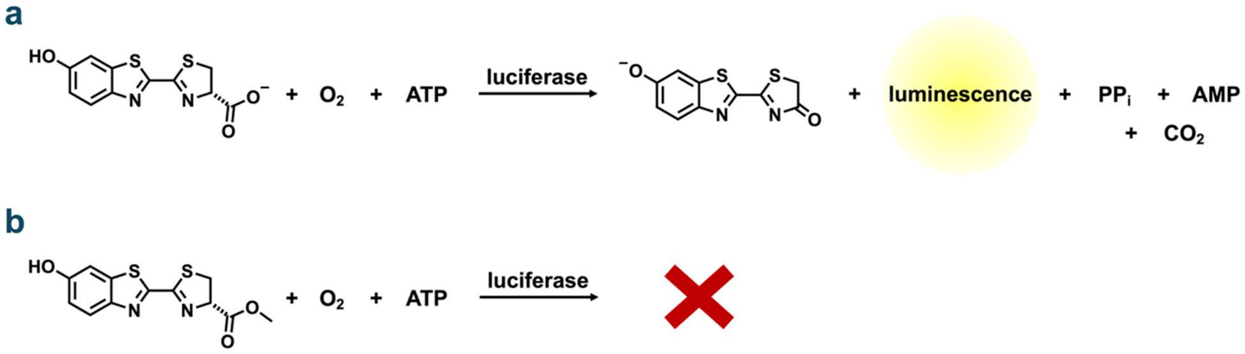


**Figure S1.** Schematic of the luciferase-mediated oxidation of **(a)** D-luciferin, resulting in the emission of light. **(b)** D-luciferin methyl ester, resulting in no emission of light.

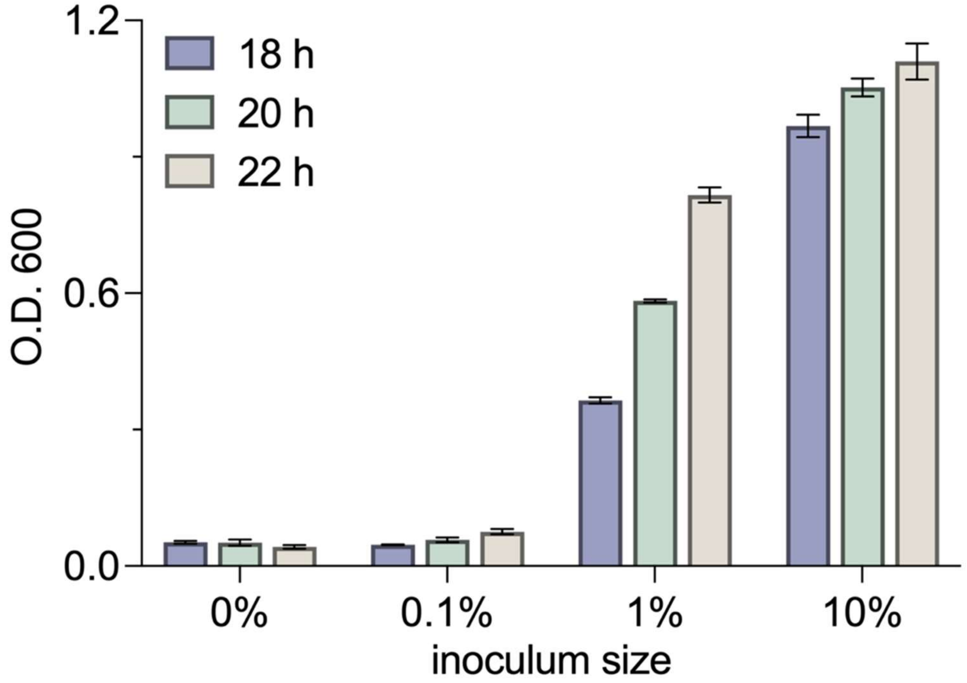


**Figure S2.** Optical density (O.D.) at 600 nm of *Msm (luc)* growth cultures from **Fig. 2d** with varying inoculum sizes and growth times. Data are represented as mean ± SD (n = 3).

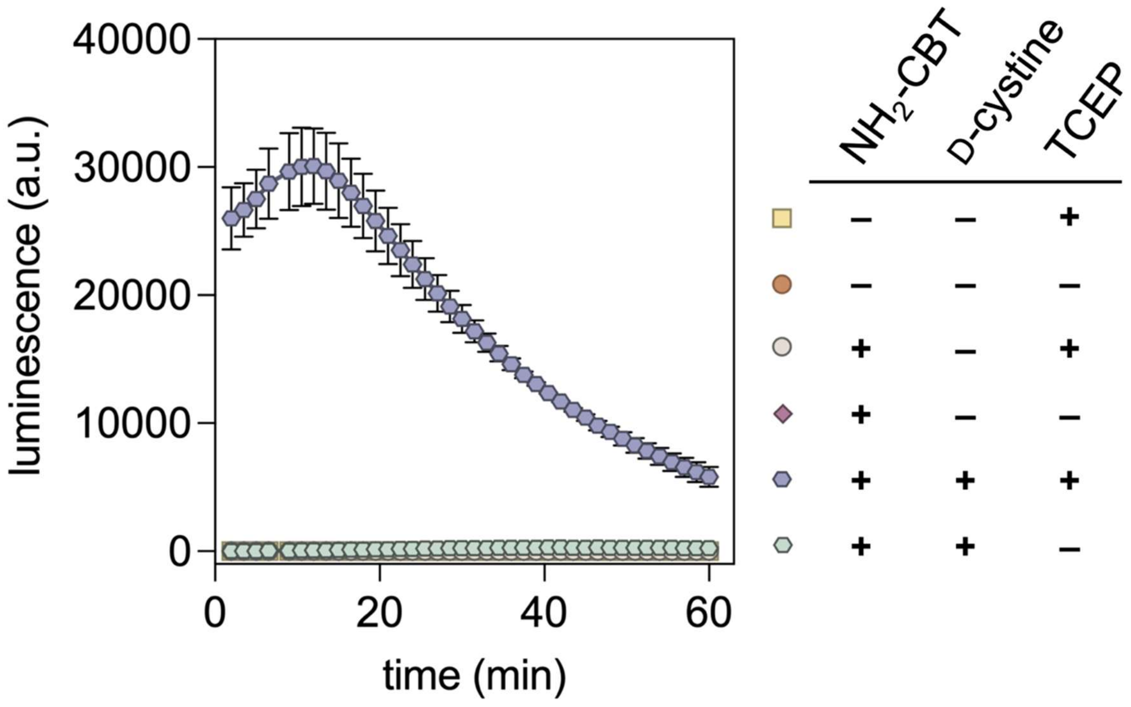


**Figure S3.** Bioluminescence analysis of a cell-free assay with luciferase in the presence or absence of $NH_2$-CBT (50 μM), D-cystine (50 μM), and TCEP (1 mM) over 60 minutes. Data are represented as mean ± SD (n = 3).

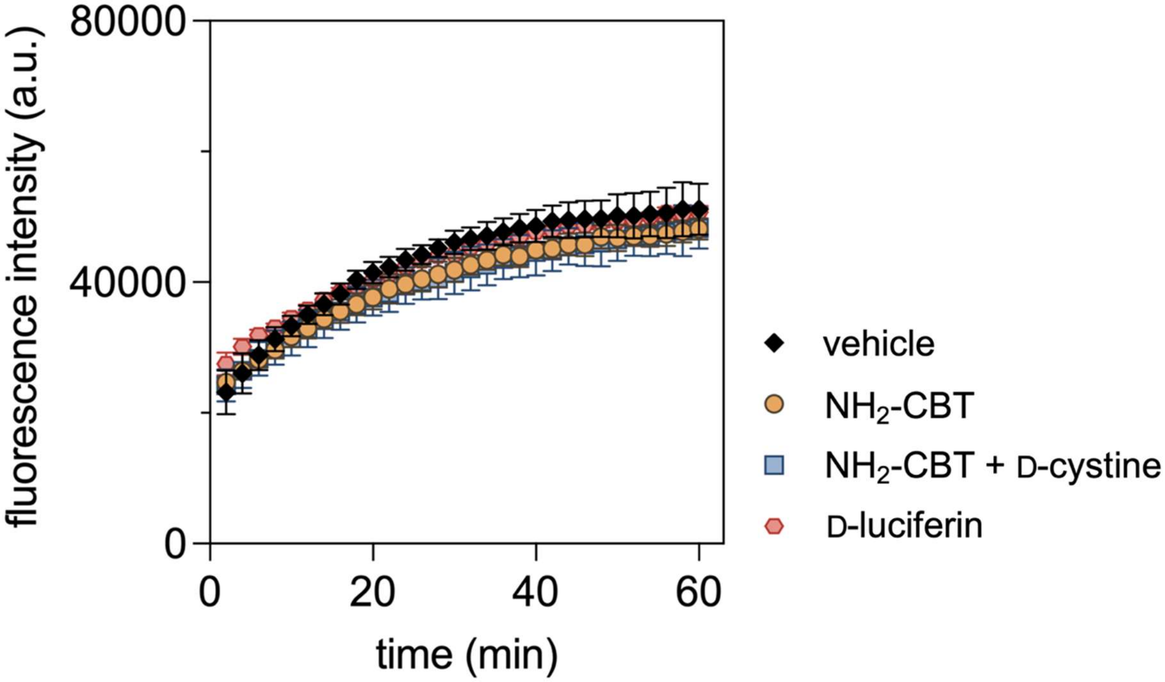
fluorescence intensity (a.u.)
80000
40000
0
0
20
40
60
time (min)
vehicle
NH2-CBT
NH2-CBT + D-cystine
D-luciferin

**Figure S4.** Nile Red-based mycobacterial membrane integrity assay. *Msm (luc)* cells were incubated with Nile Red (10 μM), $NH_2$-CBT (50 μM), and either D-cystine (50 μM), or D-luciferin (25 μM). Fluorescence was monitored over 60 minutes. Data are represented as mean ± SD (n = 3).

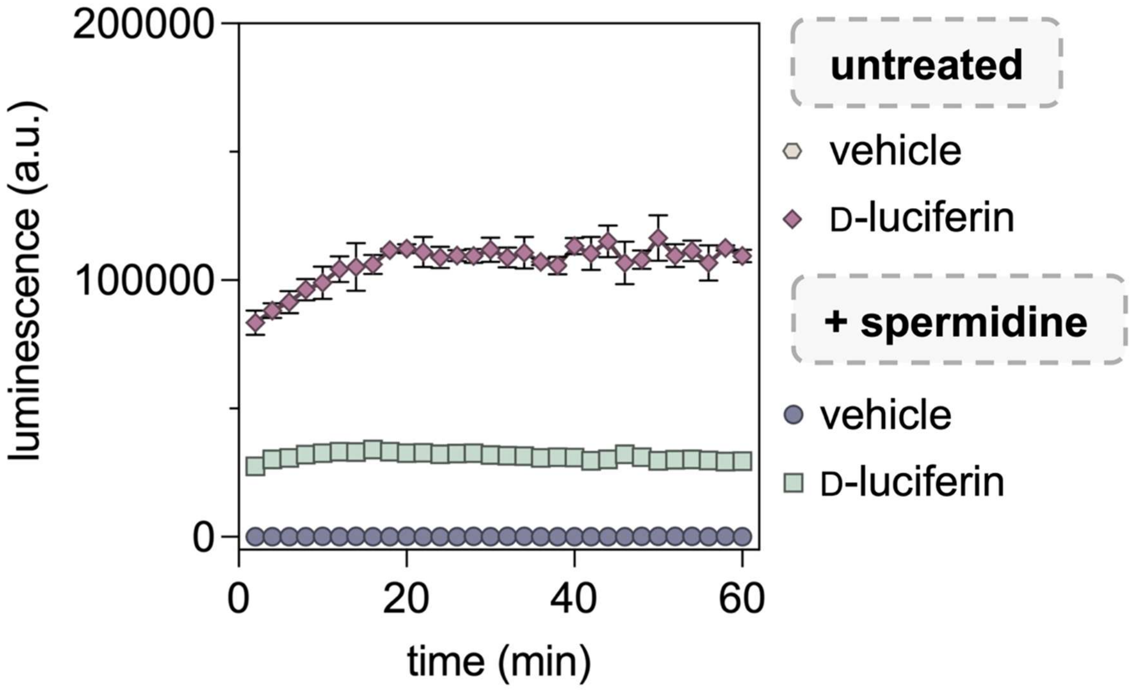


**Figure S5.** Spermidine pretreatment accumulation assay. *Msm (luc)* cells were pretreated with spermidine (10 mM) for 10 minutes, then washed as described. Luminescence was monitored over 60 minutes with or without the addition of D-luciferin (25 μM). O.D. 600 was normalized between treatments before adding spermidine. Data are represented as mean ± SD (n = 3).

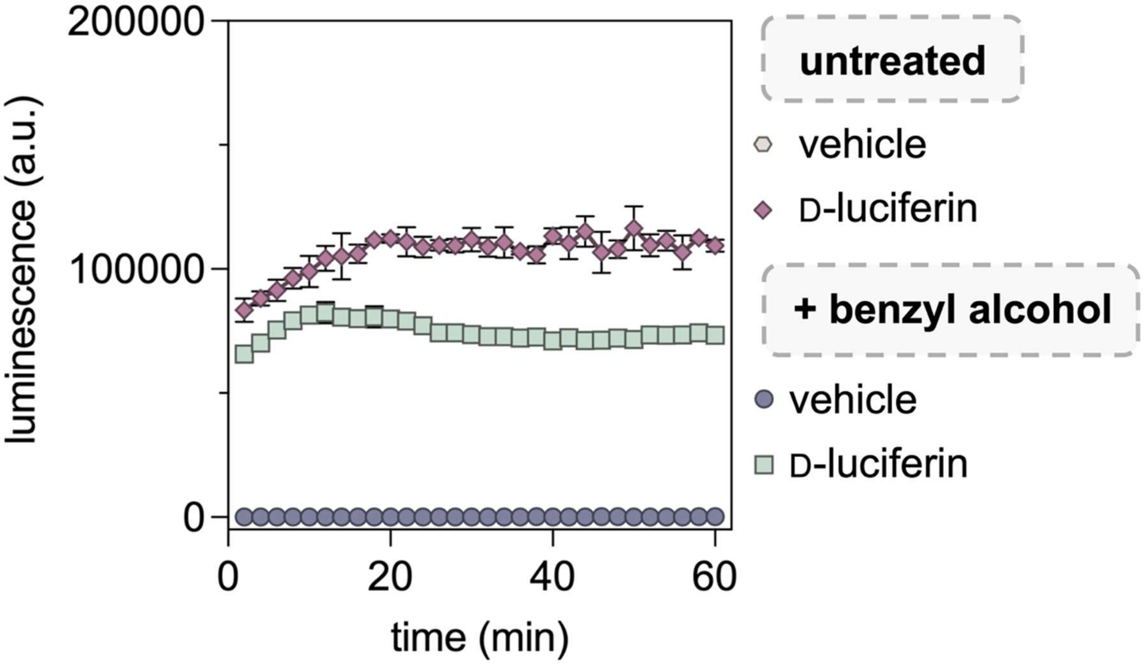


**Figure S6.** Benzyl alcohol pretreatment accumulation assay. *Msm (luc)* cells were pretreated with benzyl alcohol (25 mM) for 60 minutes, then washed as described. Luminescence was monitored over 60 minutes with or without the addition of D-luciferin (25 μM). O.D. 600 was normalized between treatments before adding benzyl alcohol. Data are represented as mean ± SD (n = 3).

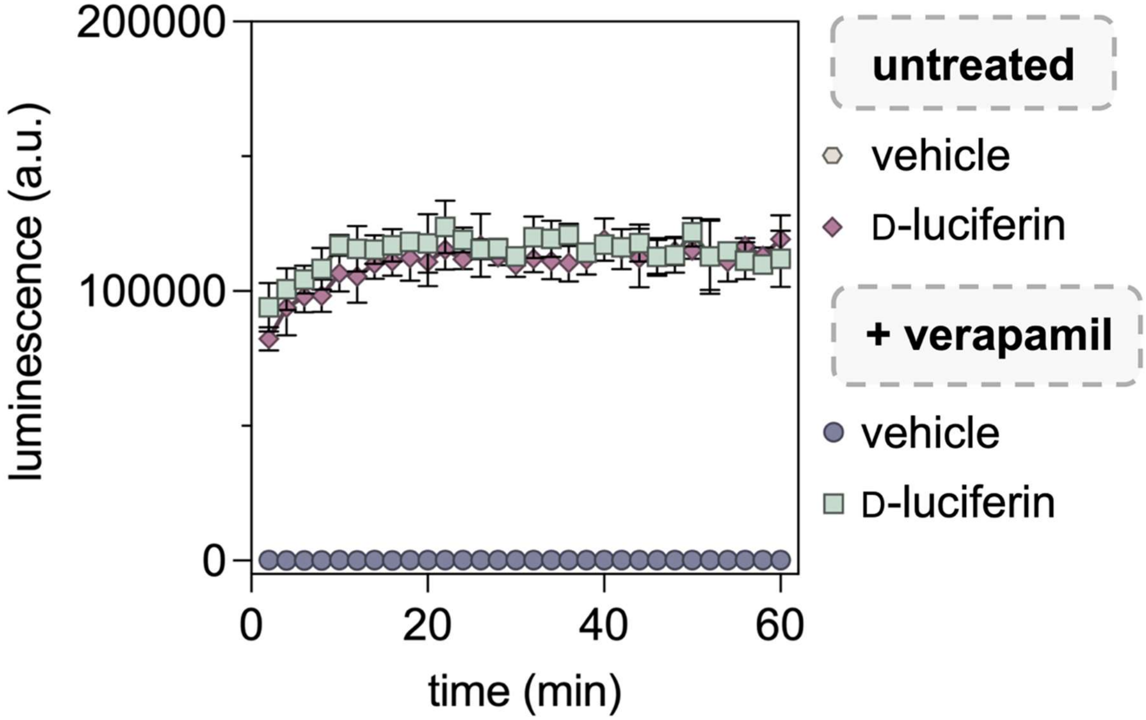


**Figure S7.** Verapamil pretreatment accumulation assay. *Msm (luc)* cells were pretreated with verapamil (75 µM) for 60 minutes, then washed as described. Luminescence was monitored over 60 minutes with or without the addition of D-luciferin (25 µM). O.D. 600 was normalized between treatments before adding verapamil. Data are represented as mean ± SD (n = 3).

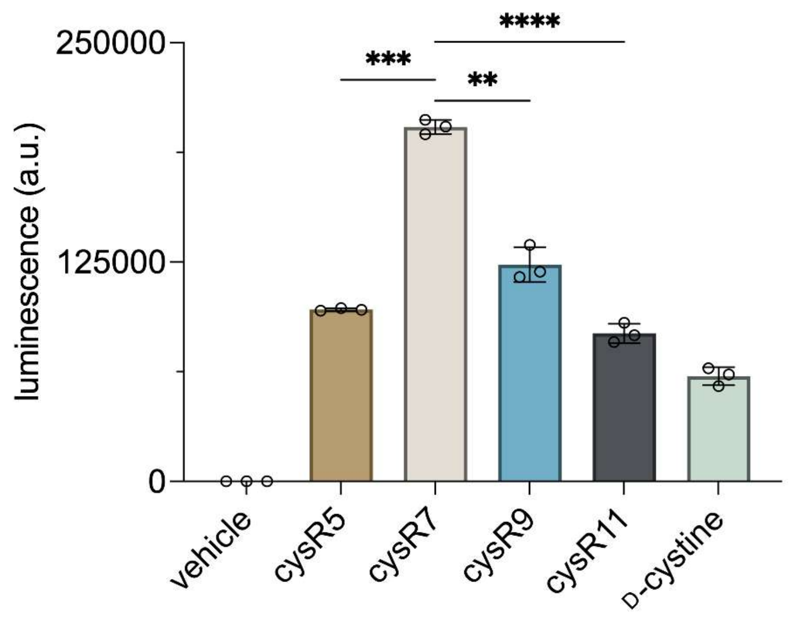


**Figure S8.** Accumulation at the 20-minute time point of D-cys-tagged polyarginine peptide conjugates. *Msm (luc)* cells were incubated with cysR5, cysR7, cysR9, or cysR11 (all 50 μM) for 20 minutes, after which luminescence was recorded. Data are represented as mean ± SD (n = 3). P-values were determined by a two-tailed t-test (ns = not significant, *p ≤ 0.05, **p ≤ 0.01, ***p ≤ 0.001, ***p ≤ 0.0001 ).

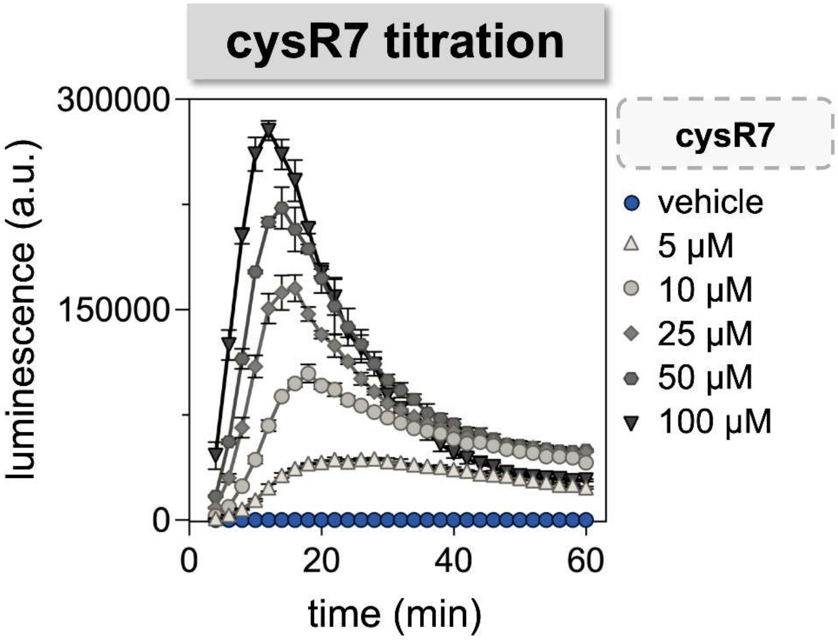


**Figure S9.** Luminescence of *Msm (luc)* cells after the addition of 50 µM $NH_2$-CBT and cysR7 (varying concentrations). Data are represented as mean ± SD (n = 3).

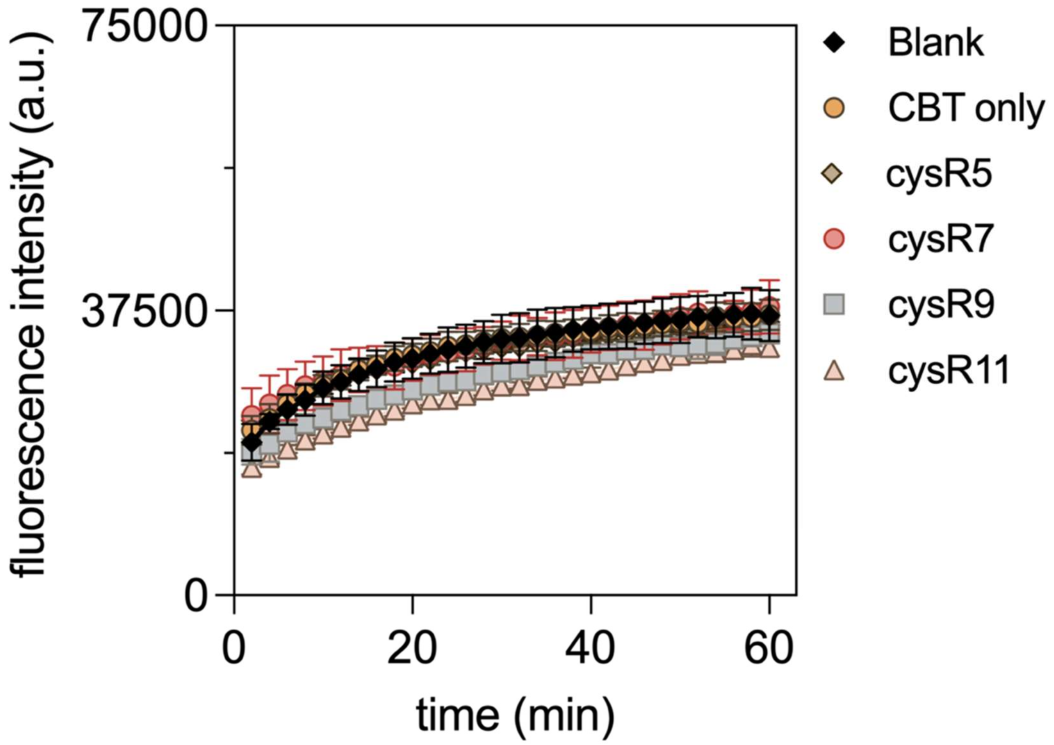


**Figure S10.** Nile Red-based mycobacterial membrane integrity assay. *Msm (luc)* cells were incubated with Nile Red (10 µM), CBT (50 µM) and either cysR5, cysR7, cysR9, or cysR11 (all 50 µM). Fluorescence was monitored over 60 minutes. Data are represented as mean ± SD (n = 3).

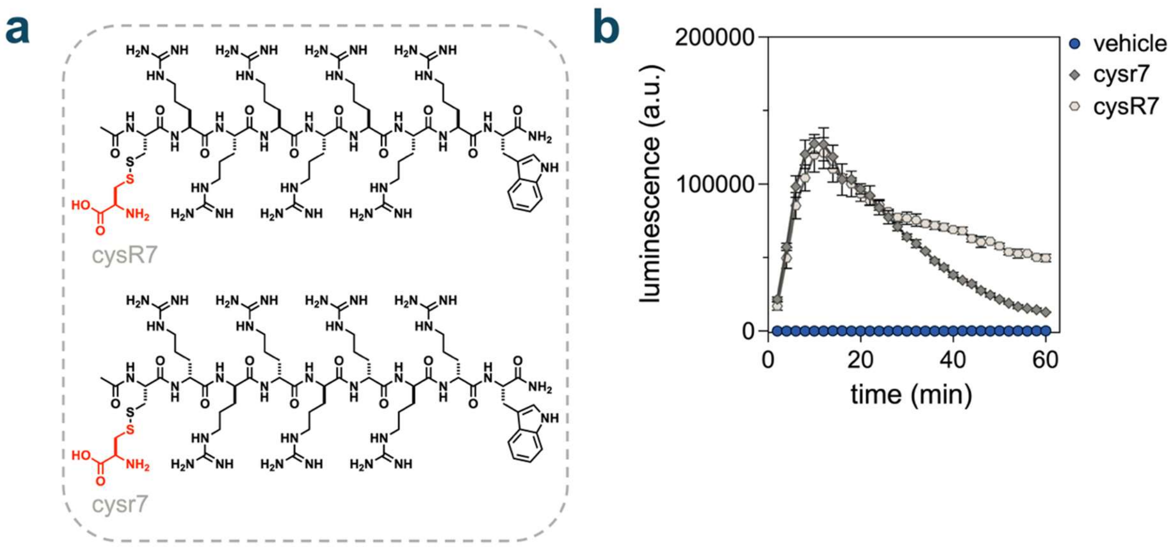


**Figure S11.** Accumulation of D-cys tagged polyarginine diastereomers in *Msm (luc)* cells. **(a)** Structure of cysr7, the D-cys tagged diastereomer of cysR7, where the stereochemistry at every arginine residue is inverted **(b)** Luminescence was monitored for 60 minutes after the addition of $NH_2$-CBT (50 μM) and either cysr7 or cys-R7 (both 50 μM). Data are represented as mean ± SD (n = 3).

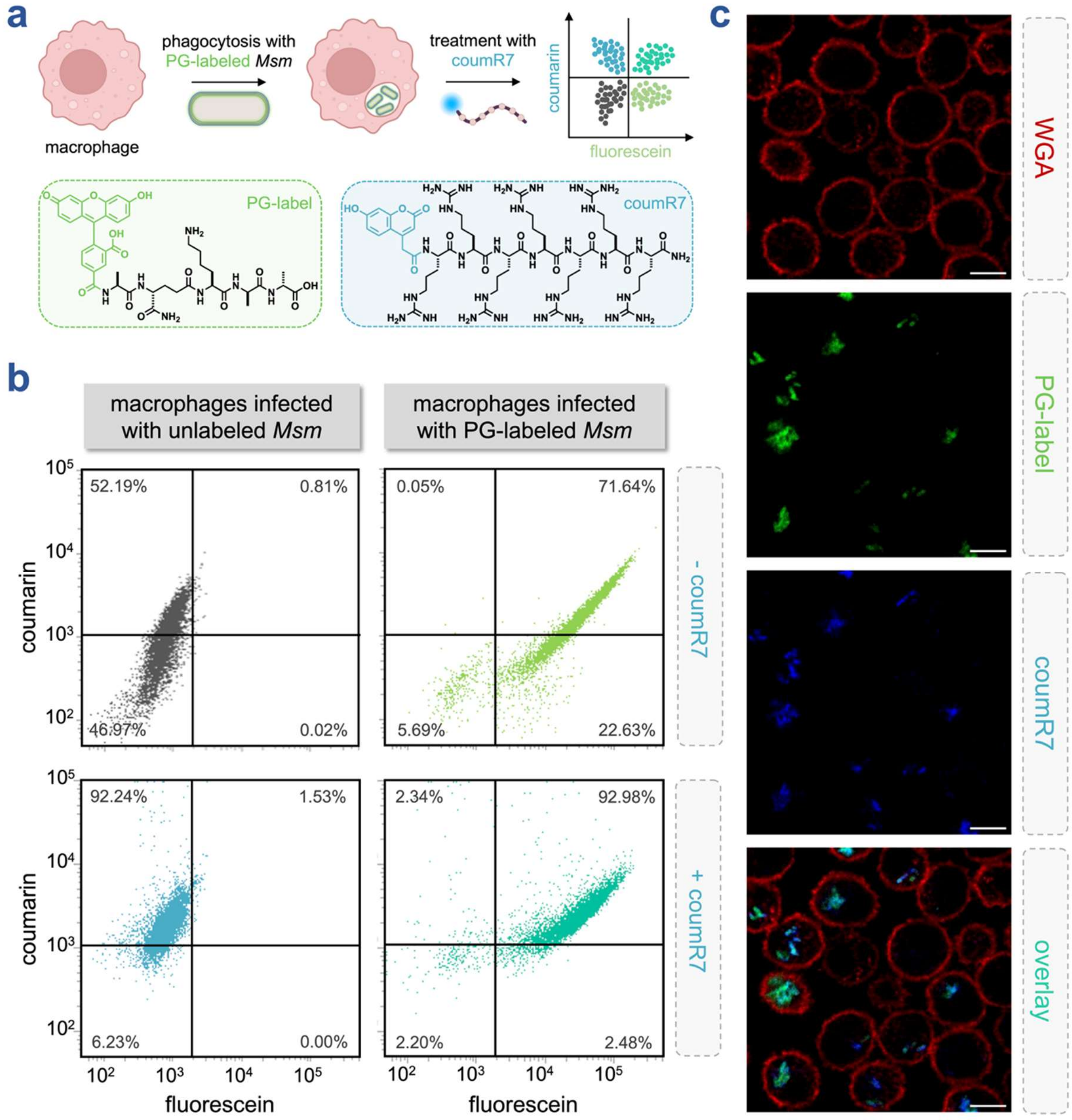


**Figure S12. (a)** Schematic depicting workflow for the infection assays of *Msm* in macrophages treated with coumR7. **(b)** Flow cytometry scatter plots of macrophages infected with either unlabeled or PG-labeled (25 μM) *Msm (wt)*, in the absence or presence of coumR7 (10 μM). **(c)** Confocal analysis of PG-labeled (25 μM) *Msm (wt)* in macrophages treated with coumR7 (10 μM). Macrophage membranes were labeled with 5 μg/mL wheat germ agglutin (WGA) tetramethylrhodamine. Scale bar = 10 μm.

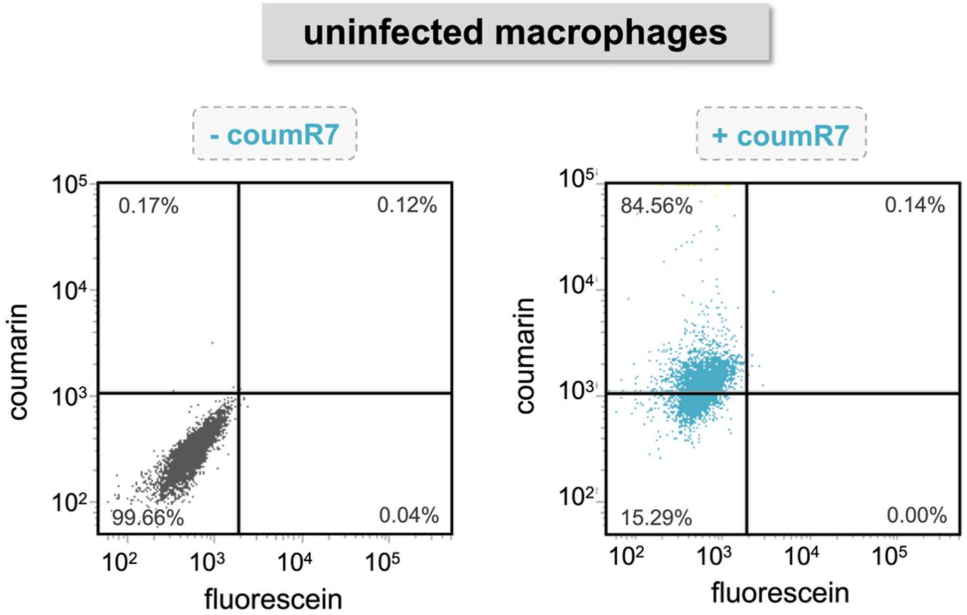


**Figure S13.** Flow cytometry scatter plots of uninfected macrophages in the absence or presence of treatment with coumR7. J774A.1 macrophages were incubated with 10 µM of coumR7 for 30 min, fixed with 4% formaldehyde in PBS, and 10,000 events per sample were analyzed *via* flow cytometry.

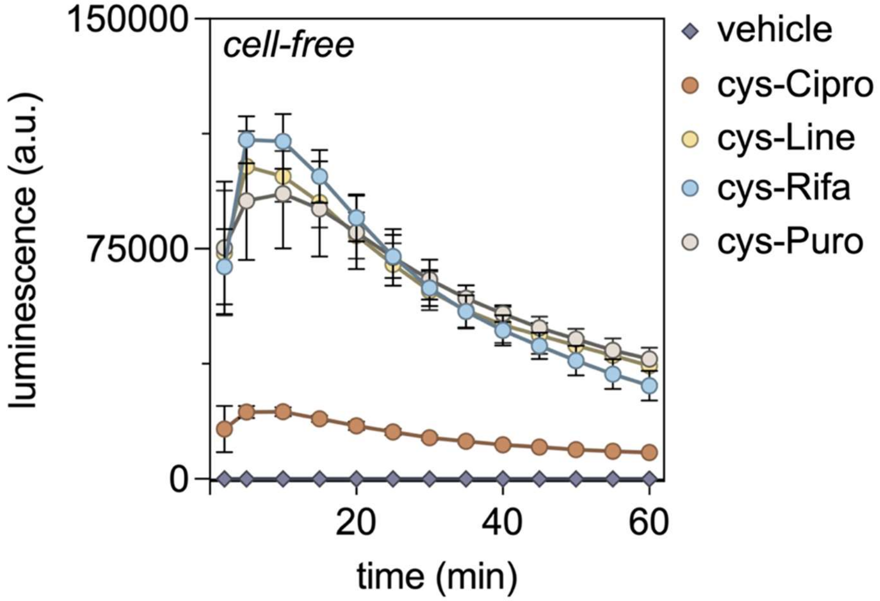


**Figure S14.** Bioluminescence analysis of a cell-free assay with D-cys tagged antibiotic conjugates (all 50 μM) in the presence of $NH_2$-CBT (50 μM) and TCEP (1 mM) over 60 minutes. Data are represented as mean ± SD (n = 3).

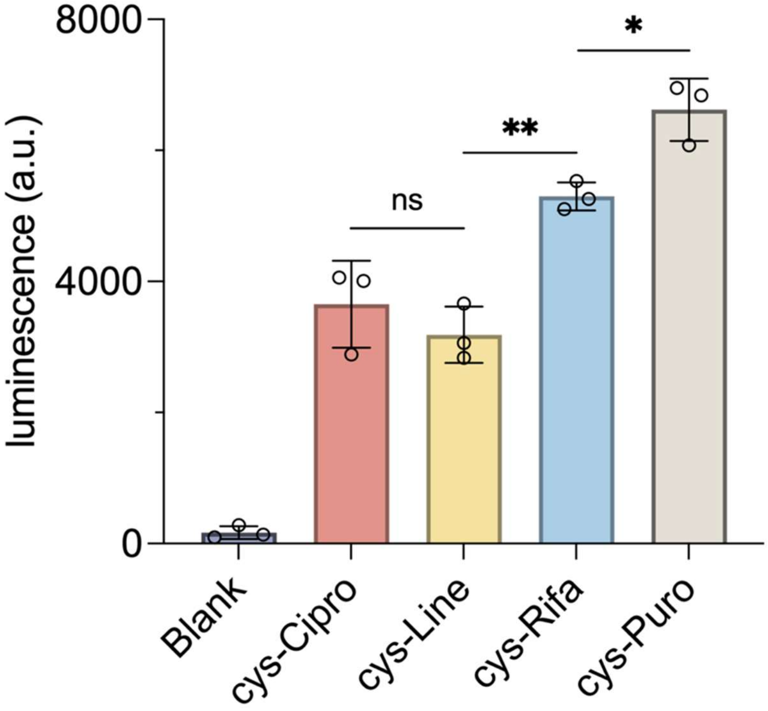


**Figure S15.** Endpoint accumulation of D-cys tagged antibiotic conjugates in *Msm (luc)*. *Msm (luc)* cells were incubated with cys-Cipro, cys-Line, cys-Rifa, or cys-Puro (all 50 μM) for 60 minutes, after which luminescence was recorded. Data are represented as mean ± SD (n = 3). P-values were determined by a two-tailed t-test (ns = not significant, *p ≤ 0.05, **p ≤ 0.01, ***p ≤ 0.001, ***p ≤ 0.0001 ).

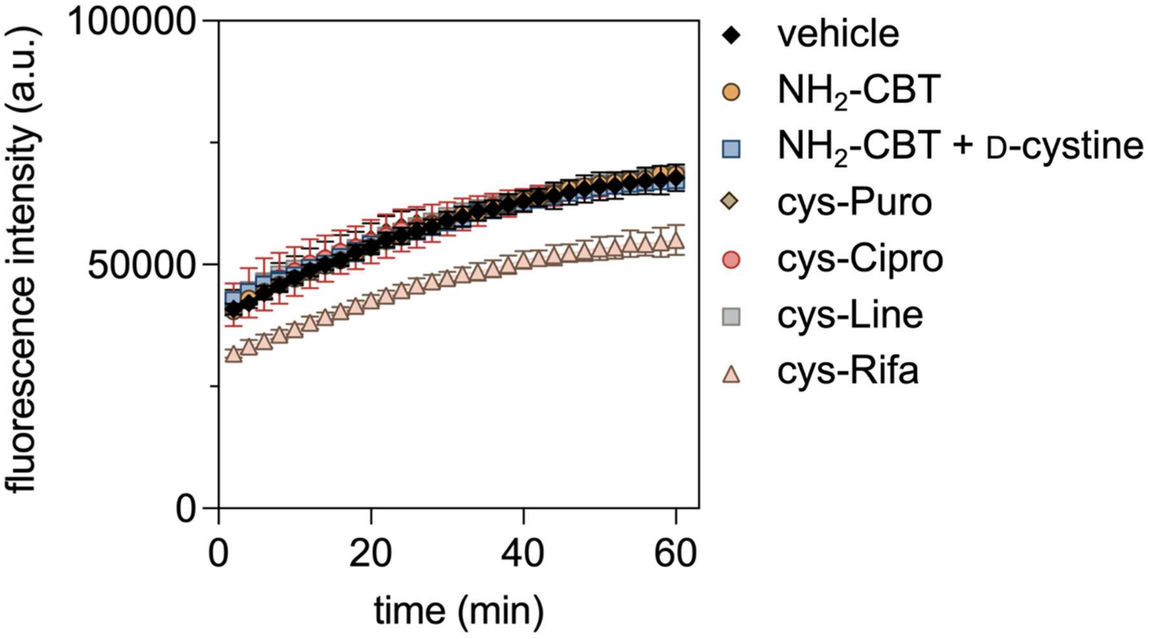


**Figure S16.** Nile Red-based mycobacterial membrane integrity assay. *Msm (luc)* cells were incubated with Nile Red (10 μM), $NH_2$-CBT (50 μM), and either D-cystine, cys-Cipro, cys-Line, cys-Rifa, or cys-Puro (all 50 μM). Fluorescence was monitored over 60 minutes. Data are represented as mean ± SD (n = 3).

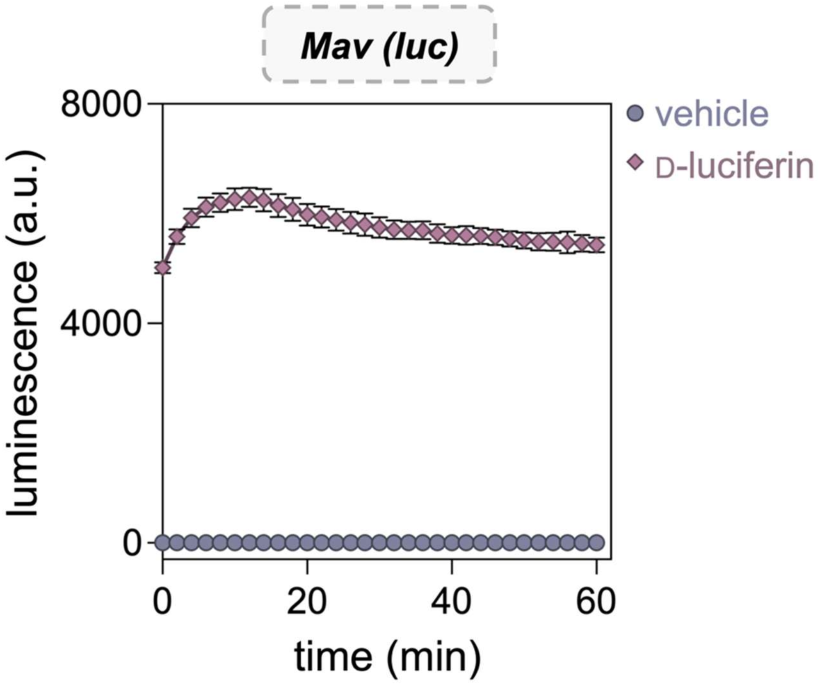


**Figure S17.** Luminescence of *Mav (luc)* cells in the presence or absence of D-luciferin (25 μM). Data are represented as mean ± SD ($n$ = 3)

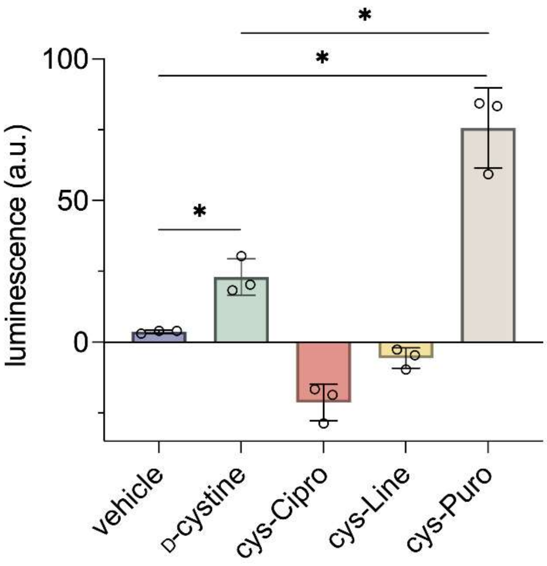


**Figure S18.** Endpoint accumulation of D-cys tagged antibiotic conjugates in *Mav (luc)*. *Mav (luc)* cells were incubated with cys-Cipro, cys-Line, cys-Rifa, or cys-Puro (all 50 μM) for 60 minutes, after which luminescence was recorded. Data are represented as mean ± SD (n = 3). P-values were determined by a two-tailed t-test (ns = not significant, *p ≤ 0.05, **p ≤ 0.01, ***p ≤ 0.001, ***p ≤ 0.0001 ).

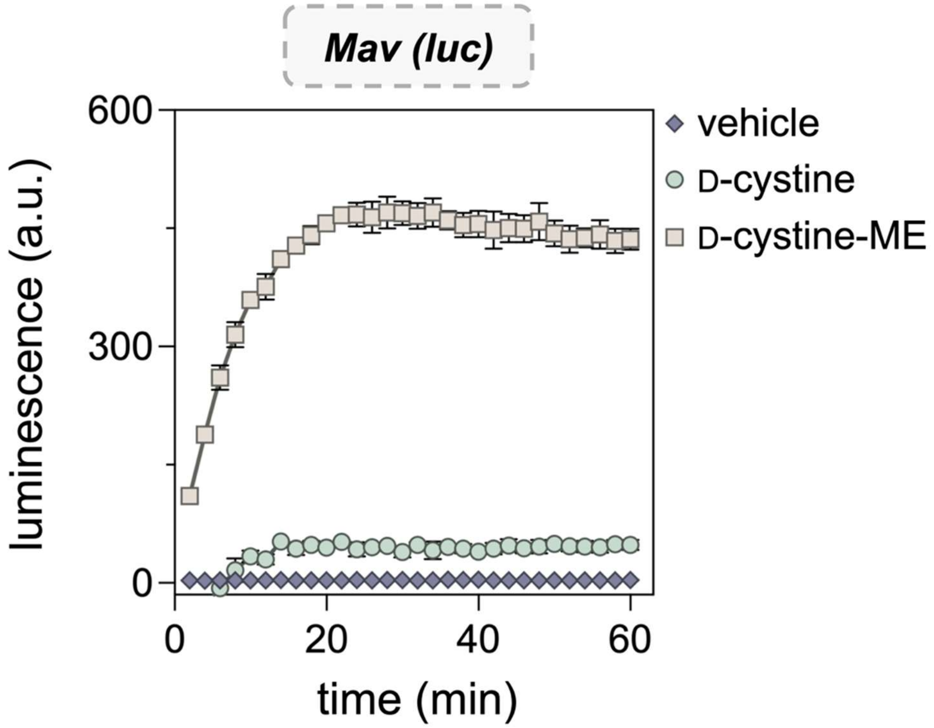


**Figure S19.** Luminescence of *Mav (luc)* cells in the presence of $NH_2$-CBT and D-cystine or D-cystine-ME. All compounds were tested at 50 μM. Data are represented as mean ± SD (*n* = 3).

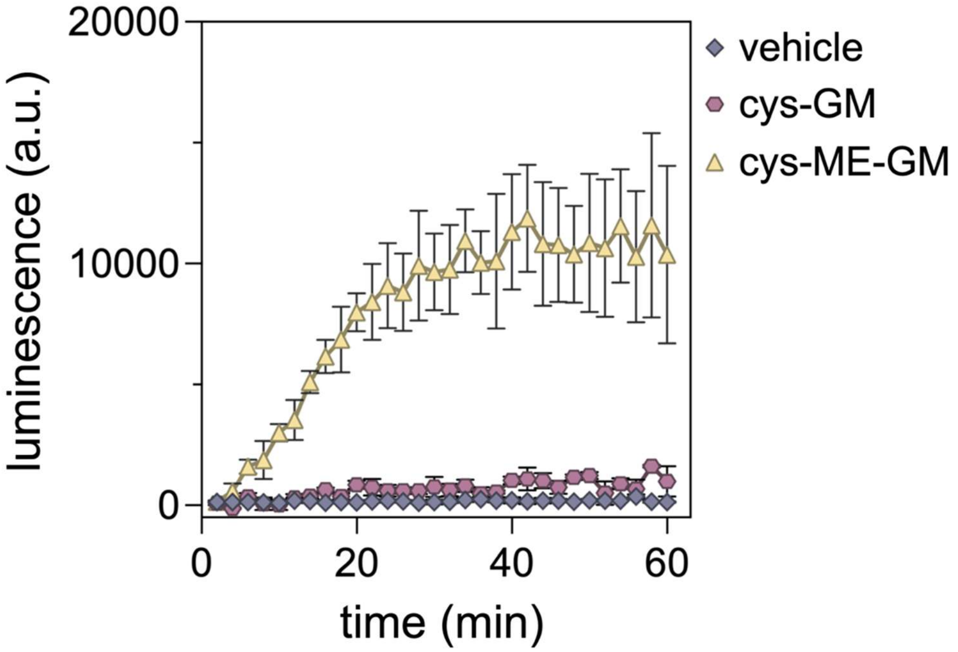

**Figure S20.** Luminescence of *Msm (luc)* cells in the presence of $NH_2$-CBT and cys-GM or cys-ME-GM measured over 60 minutes. All compounds were at 50 µM. Data are represented as mean ± SD (*n* = 3).

### EXPERIMENTAL PROCEDURES

#### Materials

| REAGENTS | VENDOR SOURCE | CATALOG # |
|---|---|---|
| **Reagents for biological methods** | | |
| Middlebrook 7H9 media | VWR | 90003-876 |
| Catalase from bovine liver | Millipore Sigma | C1345 |
| D(+)-Glucose, anhydrous (Dextrose) | Chem Impex | 00805 |
| Bovine serum albumin fraction V | Millipore Sigma | 10735078001 |
| Glycerol | Millipore Sigma | G7893 |
| Tween 80 | VWR | 97061-674 |
| Oleic Albumin Dextrose Catalase (OADC) | BD | BD 212351 |
| L- leucine | Millipore Sigma | L8000 |
| D-Pantothenic acid hemicalcium salt | Millipore Sigma | 21210 |
| Citric acid | Millipore Sigma | 791725 |
| Sodium Phosphate Dibasic Anhydrous | Fisher Scientific | S374-500 |
| Tris Base | Fisher Scientific | BP152-1 |
| Nile Red | Chem Impex | 22855 |
| Ethidium bromide solution (10 mg/mL), Biotechnology Grade | VWR | 97064-970 |
| Spermidine | Thermo Scientific | A19096-06 |
| Benzyl Alcohol | Ambeed | A194236 |
| Verapamil hydrochloride | Chem Impex | 38355 |
| pMV306G13+FFluc | Addgene | 26157 |
| Electroporation Cuvettes, 0.2 cm gap | Bio-Rad | 652086 |
| QuantiLum® Recombinant Luciferase | Promega | E170A |
| Dulbecco’s Modified Eagle Medium (DMEM) | Fisher Scientific | 11885-084 |
| Fetal Bovine Serum | Fisher Scientific | A52568-01 |
| Penicillin-Streptomycin | Fisher Scientific | P4333 |
| Wheat Germ Agglutinin (WGA) Rhodamine | Vector Laboratories | RL-1022-5 |
| **Reagents for the synthesis and characterization of test molecules** | | |
| α-*N*-Fmoc-amino acids | Chem Impex | Various |
| D-Cystine | Chem Impex | 02872 |
| Rink amide resin, (0.3-0.6 meq/g, 100-200 mesh) | Chem Impex | 12662 |

| | | |
|---|---|---|
| *N*, *N′*-Diisopropylethylamine (DIEA) | Chem Impex | 00141 |
| *N*, *N′*-Diisopropylcarbodiimide (DIC) | Chem Impex | 00110 |
| Ethyl Cyano(hydroxyimino)acetate (Oxyma) | TCI Chemicals | E0847 |
| *N, N*-Dimethylformamide (DMF) ACS-grade, anhydrous | Millipore Sigma | 319937, 589565 |
| Dichloromethane (DCM) ACS-grade, anhydrous | Millipore Sigma | D65100, 270997 |
| Methanol ($CH_3OH$) ACS-grade, HPLC-grade | Millipore Sigma | 179337, 34860 |
| Acetonitrile ($CH_3CN$) ACS-grade, HPLC-grade | Millipore Sigma | 360457, 34851 |
| Anhydrous Tetrahydrofuran (THF) | Millipore Sigma | 186562 |
| Piperidine | Millipore Sigma | 104094 |
| Trifluoroacetic acid (TFA) HPLC grade | Millipore Sigma | 302031 |
| Trifluoroacetic acid (TFA) reagent-grade | Chem Impex | 00289 |
| Ammonium Bicarbonate | Fisher Scientific | A643-500 |
| Triisopropylsilane | Chem Impex | 01966 |
| 1,3-Dimethoxybenzene | Millipore Sigma | 126306 |
| Ellman’s Reagent (5,5-dithio-bis-(2-nitrobenzoic acid)) | Thermo Scientific | 22582 |
| *N*-Boc-D-Cys | Chem Impex | 16133 |
| Ciprofloxacin | Millipore Sigma | 17850 |
| 4-(Dimethylamino)pyridine | Millipore Sigma | 107700 |
| *N*-(3-Dimethylaminopropyl)-*N*'-ethylcarbodiimide hydrochloride | Millipore Sigma | 03450 |
| (S)-5-(Aminomethyl)-3-(3-fluoro-4-morpholinophenyl)oxazolidin-2-one | AK Scientific | Z3498 |
| 1,1'-Carbonyldiimidazole | Millipore Sigma | 21860 |
| D-Cysteine | Chem Impex | 21754 |
| Puromycin dihydrochloride | Alfa Aesar | J61278 |
| 2-(Pyridin-2-yldisulfanyl)ethanamine | Ambeed | A446494 |
| Diethyl ether | Millipore Sigma | 673811 |
| Rifamycin B | Toronto Research Chemicals | R508170 |
| *N*,*N′*-Dicyclohexylcarbodiimide | Millipore Sigma | D80002 |
| Triethylamine | Millipore Sigma | 471283 |
| **General materials and equipment** | | |
| 96-well black flat bottom plates | Greiner | 655076 |

| | | |
|---|---|---|
| 96-well clear untreated round bottom plates | VWR | 82050-622 |
| T75 treated culture flasks | Fisher Scientific | 07-000-229 |
| 48 well tissue culture plate | VWR | 10062-898 |
| Glass bottom 35 mm petri dishes | MatTek | P35GC-1.5-14-C |

### Biological Methods

#### Preparation of electrocompetent mycobacteria cells and transformation.

*Mycobacterium smegmatis* mc²155 (*Msm*), and *Mycobacterium avium* clinical isolate 11 (MAH 11)[1] were grown in 100 mL liquid medium to mid-log phase (OD600 = 0.5–0.7. Cells were harvested by centrifugation at 5000 × g for 10 min at 4°C in a Multifuge X Pro series centrifuge (Thermo Fisher Scientific) and resuspended in half the original culture volume of ice-cold 10% glycerol. The cells were washed several times with ice-cold 10% glycerol, concentrating two-fold with each wash, until a final volume of 2 mL was obtained. Competent cells were aliquoted into 200 µL portions and stored at −80°C.

For transformation, electroporation cuvettes were used with electroporation settings of 2.5 kV, 25 µF, and 200 Ω on a Bio-Rad gene pulser II. Cuvettes were pre-chilled on ice, and 100–150 µL of thawed competent cells were added per cuvette. One microgram of the firefly luciferase plasmid pMV306G13+FFluc (a gift from Brian Robertson & Siouxsie Wiles) was pipetted onto the cuvette wall before adding the cells to facilitate mixing. The cuvette exterior was dried, placed in the electroporator, and pulsed under the preset conditions.

Immediately after electroporation, cells were recovered by transferring them as quickly as possible into a vial containing pre-warmed, antibiotic-free 7H9 medium. Cultures were incubated for 3 h (for *Msm*) or overnight (for *Mav*) at 37°C with shaking, then pelleted by centrifugation. The cell pellet was resuspended in 7H9 and 100 µL of the suspension was plated as is, while the rest of the culture was plated after centrifugation to concentrate cells. Culturing of the resulting colonies on 50 µg/mL kanamycin-containing agar plates yielded *Msm (luc)* and *Mav (luc).*

#### Mycobacteria cell culture.

*Msm (luc)* or *Msm (wt)* cells were grown in 7H9 media with 0.5% glycerol, 0.05% tween 80, and 1x ADC enrichment. 10x ADC was made with 5 g bovine serum albumin, 2 g dextrose, 3 mg catalase in 100 mL autoclaved milliQ $H_2O$. Glycerol stocks were prepared from stationary-phase cells in 30% glycerol, and aliquots were stored at −80°C. Unless otherwise mentioned, *Msm (luc)* cells were inoculated from glycerol stocks into 7H9 medium at a 1:100 dilution in standard culture tubes and grown for 22 h at 37°C until reaching an $OD_{600}$ of approximately 1.0. Whenever appropriate, *Msm (wt)* cells were also grown to $OD_{600}$ of approximately 1.0 as a control. For assays evaluating dilution conditions, cultures were initiated at 1:10, 1:100, and 1:1000 dilutions. Cells were then harvested by centrifugation at 3000 × g for 2 min in a in a HERAEUS multicentrifuge ×1 centrifuge (Thermo Fisher Scientific), washed twice with phosphate-buffered saline containing 0.05% Tween 80 (PBST, pH 7.4) and resuspended in PBST for subsequent assays.

*Mav (luc)* was grown in 7H9 medium prepared as before, except ADC contained NaCl added to a final concentration of 0.9%, instead of catalase. For preparation of stocks, *Mav* *(luc)* cells were grown until OD600 ~0.6, aliquoted, and frozen at -80°C. Cells were inoculated from stocks into 7H9 Kanamycin 50 ug/mL medium at a 1:50 dilution in 50 mL conical tubes (double-contained) and grown for 48h at 37°C until reaching an OD600 of approximately 0.6.

For MIC assays, double auxotroph *M. tuberculosis* mc$^2$6206 strain (H37Rv Δ*panCD* Δ*leuCD*1) was provided by Dr. William Jacobs and was cultured in Middlebrook 7H9 supplemented with 0.5% glycerol, 10% Middlebrook Oleic Albumin Dextrose Catalase (OADC; BD), and either 0.05% Tyloxapol, for strain maintenance, or 0.05% Tween-80, for experimental manipulation.[2] Growth media were additionally supplemented with 50 µg/mL l-leucine and 50 µg/mL pantothenic acid. Bacterial stocks were prepared by directly freezing at -80°C *M. tuberculosis* culture at OD600 ~0.5.

**Mammalian cell culture.**

J774A.1 cells were cultured in Dulbecco's modified Eagle's medium (DMEM) supplemented with 10% (v/v) FBS, 50 IU/mL penicillin, and 50 µg/mL streptomycin in a T75 flask and maintained in a humidified atmosphere of 5% $CO_2$ at 37°C.

**Luminescence imaging of multiwell plates.**

In a 96-well, black, flat-bottomed plate, D-luciferin was added to a final concentration of 100 µM and imaged using a Bio-Rad ChemiDoc XRS+. Plates were placed directly on the imaging platform, and luminescence was captured using the "no filter, no illumination" setting, with exposure time automatically optimized for detection of low-intensity signals. Luminescence images were overlaid with plate images taken under white epi-illumination without a filter. Images were processed and analyzed using the instrument software to visualize relative luminescence across wells.

**Bioluminescence-based accumulation assays.**

In a 96-well, black, flat-bottomed plate, molecules of interest were added to achieve the desired final concentrations in a total volume of 100 µL. Unless otherwise mentioned, D-luciferin was assessed at a concentration of 25 µM whereas D-cystine, *N*-acetyl-D-cysteine (NAC), CBT, antibiotic conjugates and polyarginine peptide conjugates were assessed at 50 µM. Stock solutions of these molecules were prepared in PBS. PBS was also added to blank and negative control wells to maintain equal total volumes and cell numbers across all wells. A stock solution of the appropriate CBT (2-cyanobenzothiazole) was prepared in DMF, and calculated volumes were added to culture tubes containing cells to achieve the target concentrations. Equal volumes of washed and resuspended cells were then transferred to each well, mixed, and the plate was placed in an Agilent BioTek Synergy H1 Microplate Reader set to kinetic luminescence mode, typically recording every 2 min over 60 min. During all luminescence measurements, continuous slow double-orbital shaking at 237 cpm was applied. Luminescence was measured using fiber optics, with the gain set to 200, integration time of 0:01:00, and a 100 ms delay after plate movement. The read height was 4.5 mm, and the temperature was maintained at 37 °C.

For assays assessing luminescence in the pellet and supernatant of centrifuged luciferase-expressing mycobacterial cells, the plate was initially read for 30 min. Selected wells treated with D-luciferin were then transferred to a separate plate and pelleted. The supernatant was carefully separated from the pellet (which was resuspended in PBST), following which both were returned to the plate. Non-pelleted cells treated with D-luciferin served as controls. Luminescence was subsequently recorded after a 60 min incubation period.

For assays designed to mimic phagosomal pH, cells were resuspended in a phosphate-citrate buffer consisting of 0.2 M disodium phosphate and 0.1 M citric acid, with the pH adjusted to 4.5. In spermidine/benzyl alcohol/verapamil pretreatment experiments, an incubation with the appropriate compound was performed at this time at the desired concentrations (10 mM spermidine for 10 min; 25 mM benzyl alcohol for 60 mins, and 75 µM verapamil for 60 mins). Cells were then washed and resuspended in 1× PBST in the same manner as described above. For experiments involving pretreatments or multiple strains, the optical density (O.D.) was normalized between treatment arms to ensure consistent cell numbers across all conditions.

**Cell-free luciferase assays.**

The 14.9 mg/mL luciferase enzyme stock (QuantiLum® Recombinant Luciferase received from Promega) was diluted in 25 mM Tris buffer (pH 8) to prepare 0.4 mg/mL aliquots (40 µL each), which were stored at −80 °C. All assays were conducted in 25 mM Tris buffer (pH 8) in 96-well, black, flat-bottomed plates. Molecules of interest were added to achieve the desired final concentrations in a total volume of 100 µL. Appropriate volumes of $MgSO_4$ (100 mM), freshly prepared ATP (20 mM), and luciferase stock (0.4 mg/mL) were added to yield final concentrations of 5 mM, 1 mM, and 20 µg/mL, respectively. Where applicable, TCEP (20 mM stock in 25 mM Tris, pH 8) was added to a final concentration of 1 mM. Plates were placed in an Agilent BioTek Synergy H1 Microplate Reader set to kinetic luminescence mode, typically recording every 2 min for 60 min. Continuous slow double-orbital shaking at 237 cpm was applied during all measurements. Luminescence was recorded using fiber optics, with gain set to 100, integration time of 0:01:00, a 100 ms delay after plate movement, read height of 4.5 mm, and temperature maintained at 37 °C.

**Nile Red-based mycobacterial membrane integrity assay**

In a 96-well, black, flat-bottomed plate, *Msm (luc)* cells were co-incubated with test compounds and Nile Red to assess membrane integrity following treatment. Appropriate volumes of cell suspension were added in triplicate, and 5 µL of a 200 µM Nile Red stock solution was added to achieve a final concentration of 10 µM. The plate was placed in an Agilent BioTek Synergy H1 Microplate Reader set to kinetic fluorescence mode, recording every 2 min over 60 min. Fluorescence was measured with excitation and emission wavelengths of 540 nm and 630 nm, respectively. Continuous slow double-orbital shaking at 237 cpm was applied, and the temperature was maintained at 37 °C.

#### Flow cytometry analysis of PG-labeled *Msm* in J774A.1 macrophages treated with coumR7.

*Msm (wt)* cells were grown in liquid medium as described above for approximately 24 hours to mid-log phase ($OD_{600}$ = 0.5-0.7) while shaking (250 rpm) at 37°C. At this time, 25 µM of tetrapeptide probe PG-label was added to the media and the cells were further grown for 18 h at 37°C while shaking (250 rpm) to a stationary phase. The bacteria were harvested and centrifuged for 2 min at 4000 x g then washed three times with PBST. J774A.1 cells were cultured as described above. On the day prior to the experiment, J774A.1 cells were seeded into a 48-well plate and allowed to adhere. On the day of the experiment, J774A.1 cells were washed with twice with 1X PBS. The PG-labeled *Msm* cells were resuspended in DMEM + 10% FBS containing no antibiotics and added to the washed J774A.1 cells at an MOI of 100. The cell mixture was then incubated at 37°C for 1 hour to induce phagocytosis. The cell mixture was washed five times with 1X PBS to remove extracellular bacteria. The cells were then incubated with a solution containing 10 µM coumR7 in DMEM containing no FBS and no antibiotics for 30 min at 37°C. The cells were washed three times with 1X PBS and fixed for 30 min with 4% formaldehyde in 1X PBS. Samples were then removed from the well plate by scraping and analyzed using an Attune NxT Flow Cytometer (Thermo Fisher) equipped with both 488 nm and 405 nm lasers with 530/30 nm and 440/50 nm bandpass filters.

#### Confocal microscopy analysis of PG-labeled *Msm* in J774A.1 macrophages treated with coumR7.

*Msm (wt)* cells were grown in liquid medium as described above for approximately 24 hours to mid-log phase ($OD_{600}$ = 0.5-0.7) while shaking (250 rpm) at 37°C. At this time, 25 µM of tetrapeptide probe PG-label was added to the media and the cells were further grown for 18 h at 37°C while shaking (250 rpm) to a stationary phase. The bacteria were harvested and centrifuged for 2 min at 4000 x g then washed three times with PBST. J774A.1 cells were cultured as described above. On the day prior to the experiment, J774A.1 cells were seeded into 35 mm glass bottom microwell dishes and allowed to adhere. On the day of the experiment, J774A.1 cells were washed with twice with 1X PBS. The PG-labeled *Msm* cells were resuspended in DMEM + 10% FBS containing no antibiotics and added to the washed J774A.1 cells at an MOI of 100. The cell mixture was then incubated at 37°C for 1 hour to induce phagocytosis. The cell mixture was washed five times with 1X PBS to remove extracellular bacteria. The cells were then incubated with a solution containing 10 μM coumR7 in DMEM containing no FBS and no antibiotics for 30 min at 37°C. The cells were washed three times with 1X PBS and fixed for 30 min with 4% formaldehyde in 1X PBS. J774A.1 macrophages were then treated with 5 μg/mL of TMR-tagged Wheat Germ Agglutinin (Vector Laboratories, RL-1022) in 1X PBS for 15 min at 4°C and then washed three times with 1X PBS. Cells were imaged using a Leica STELLARIS 8 Super resolution microscopy system (40x/1.3 oil-immersion lens) equipped with 405 nm, 488 nm, and 561 nm lasers. We acknowledge the Keck Center for Cellular Imaging for the usage of the Leica STELLARIS 8 microscopy system. [PI: AP; NIH-OD030409].

**MIC assays.**

Briefly, mycobacterial cultures were prepared as described above and grown to an $OD_{600}$ of ~0.6. Peptide stocks (25.6 mM in DMSO) were diluted in 7H9 media to 2X the highest concentration; all dilutions in this assay were done using enriched 7H9 media. 200 μL of each stock was added in triplicate to a 1 mL deep well 96-well plate, after which two-fold serial dilutions were performed. The cells were diluted 50-fold, then 200 μL of this solution was added to each well. One row of the plate was reserved for wells without peptides (growth control) and wells without peptides or cells (sterility control). The plates were covered with a porous coverslip and incubated at 37 °C while shaking at 250 RPM for 40 h. 200 μL from each well was transferred to a clear flat-bottom 96 well plate. MIC values were determined as the lowest concentration of peptide that gave $OD_{600}$ < 0.1 read using a Biotek Synergy H1 plate reader.

### SYNTHESIS AND CHARACTERIZATION OF PEPTIDE CONJUGATES

#### Polyarginine peptide conjugates.

#### Synthesis of cysR5, cysR7, cysR9, cysR11, coumR7.

The polyarginine peptide conjugates were prepared using standard Fmoc-based solid-phase chemistry using Rink-amide resin (0.48 mmol/g of loading capacity) such that the *C*-terminus would be amidated upon cleavage. Synthesis was performed in accordance with established procedures for the resin. Briefly, to a 25 mL peptide synthesis vessel, an appropriate amount of resin was added, followed by 20% piperidine in DMF (15 mL) to perform Fmoc deprotection of the resin. This was followed by shaking at room temperature for 30 min. The resin was then washed with $CH_3OH$ and DCM three times. After the last wash, 4 eqiv. of the appropriate protected amino acid was added along with 4 eqiv. of ethyl cyanohydroxyiminoacetate (Oxyma) and 4 eqiv. of *N, N'*-Diisopropylcarbodiimide (DIC). The resin was shaken at room temperature for 2 h, then washed with $CH_3OH$/ DCM. The remainder of the protected amino acids were coupled in the same manner. For coumR7, after coupling the last protected amino acid, Fmoc deprotection was performed and 4 equiv. of 7-hydroxycoumarin-4-acetic acid, 4 equiv. of Oxyma, and 4 equiv. of DIC were used to couple the fluorophore to the peptide. For all other peptides, after the addition of protected L-cysteine to the *N*-terminus, Fmoc deprotection was performed followed by acetylation, which was carried out by combining the peptide with a mixture of acetic anhydride, DIEA, and DMF (5:8.5:86.5, v/v) for 1 hour at room temperature. Peptides were then cleaved from the resin using a Trifluoracetic acid/1,3-Dimethoxybenzene/Triisopropylsilane (92.5:5:2.5, v/v/v) solution with shaking at room temperature for 2 h. The solution was filtered and concentrated prior to precipitation by the addition of cold diethyl ether to yield crude peptide. To a 50 mL conical tube, the crude peptide was then added (30 µmol, 1 equiv.) along with D-cystine (150 µmol, 5 equiv.). A mixture of $CH_3CN$ and $H_2O$ (20 mL, 1:1 v/v) was then added, followed by 0.5 M ammonium bicarbonate buffer (≈ 5 mL) to adjust the pH to 8.5. If necessary, the pH was adjusted to 8.5 with 1 M NaOH, and the reaction was incubated for 2 h at room temperature. Because the D-cystine does not fully dissolve under these conditions, the reaction was centrifuged in a in a in a HERAEUS multicentrifuge ×1 centrifuge (Thermo Fisher Scientific) for 5 min at 4000 × g, to pellet any undissolved material, and the supernatant was collected. The solvent was then removed under reduced pressure by rotary evaporation. The residue was purified by reverse-phased preparative high-performance liquid chromatography (RP-HPLC) equipped with Waters 1525 with 2489 UV/Visible Detector on a Phenomenex Luna 10 µm C8(2) 100 Å (250 x 21.2 mm) column using gradient elution with $H_2O/CH_3OH$ with 0.1% TFA at 10 mL/min. The HPLC fractions of the desired purified compounds were first concentrated under reduced pressure using a rotary evaporator. The final concentrated aqueous solutions were lyophilized to dryness using Labconco Freezone 4.5 L (-84 $^{o}$C) lyophilizer.

**Characterization of cysR5, cysR7, cysR9, cysR11, coumR7.**

The peptide conjugates were analyzed for purity using Phenomenex Luna 5 µm C8(2) on the same RP-HPLC; gradient elution in $H_2O$/$CH_3OH$ with 0.1% TFA at 1 mL/min. Peptides were analyzed *via* matrix-assisted laser desorption ionization time-of-flight (MALDI-TOF) mass spectrometry (Shimadzu 8020). All peptides in this library were characterized using UV-Vis absorbance of the tryptophan residue in their sequence at 280 nm ($\varepsilon$ = 5690 $cm^{-1}M^{-1}$). See data starting next page.

**cysR5**

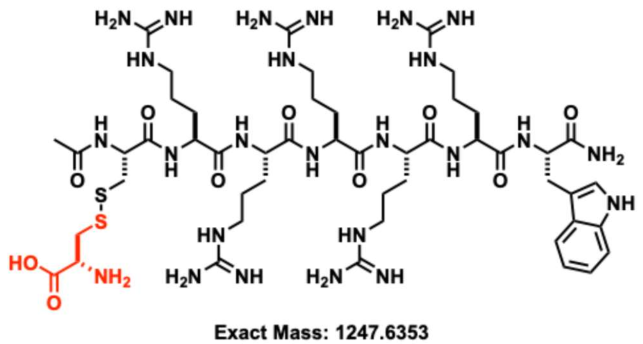


Chemical structure of **cysR5**.

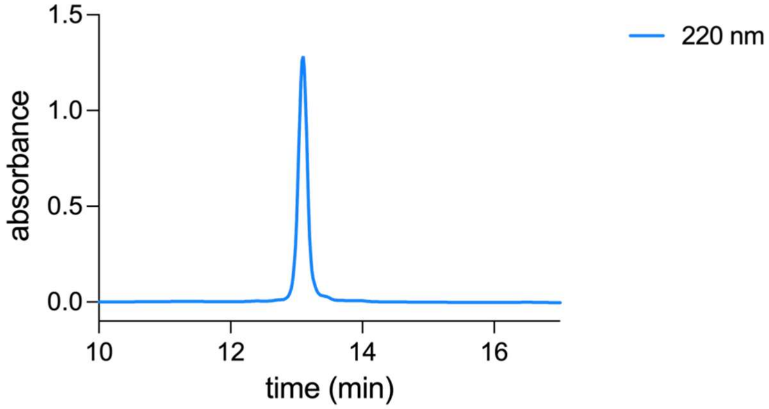


Analytical HPLC Chromatogram of **cysR5**.

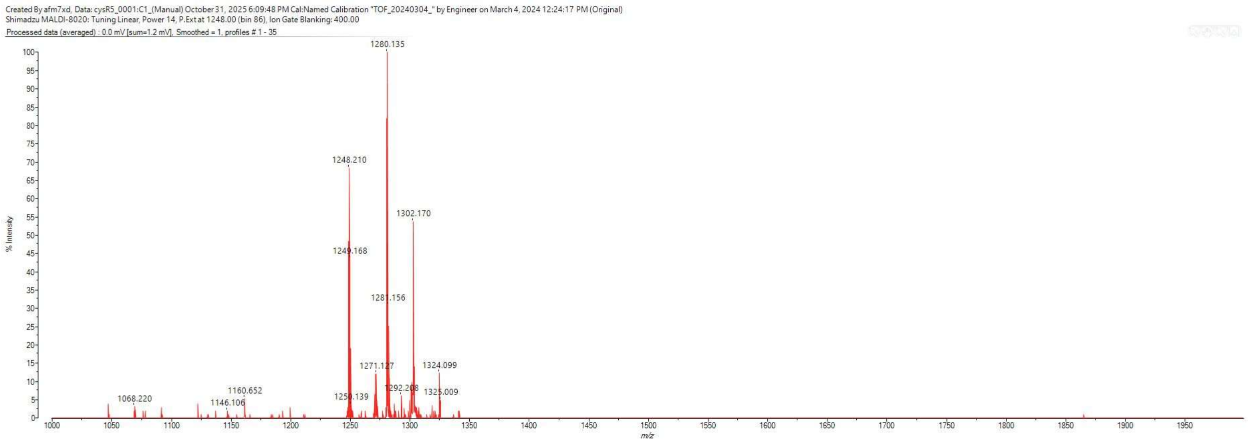


MALDI-TOF mass spectrum for **cysR5** (m/z 1248.643 for $[M+H]^+$).

**cysR7**

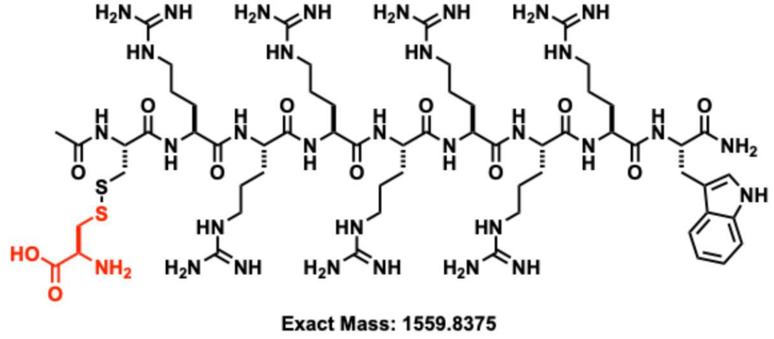


Chemical structure of **cysR7**.

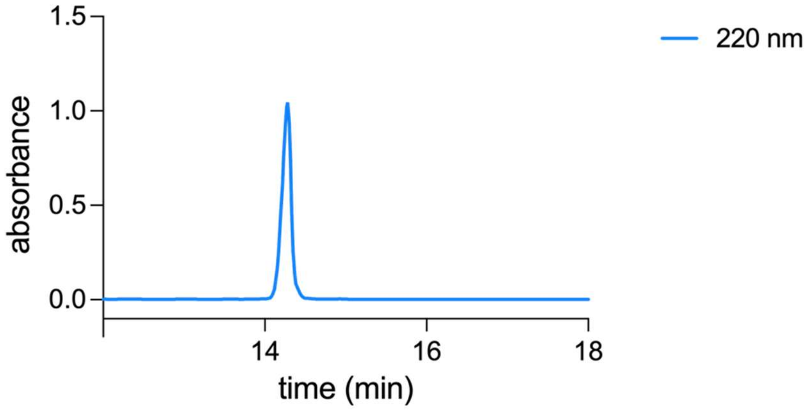


Analytical HPLC Chromatogram of **cysR7**.

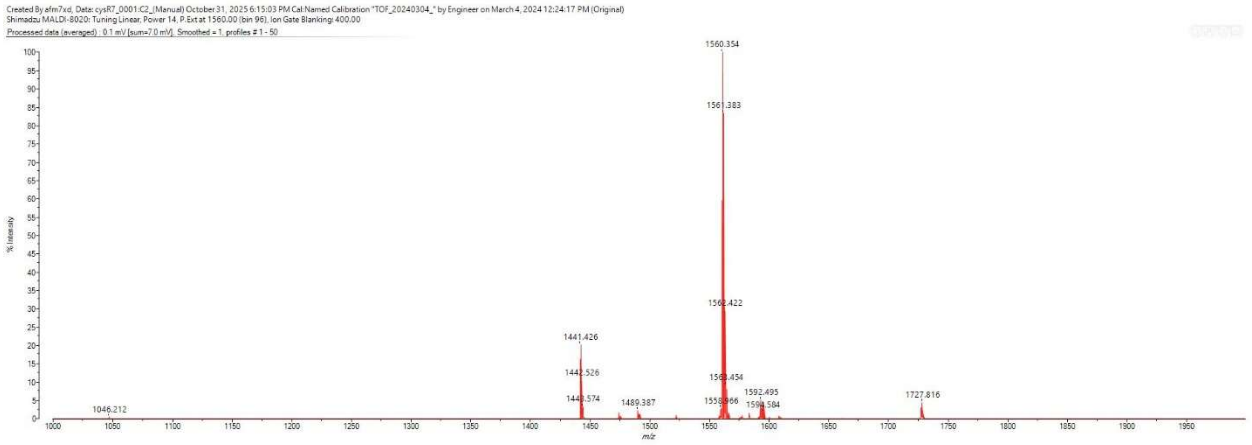


MALDI-TOF mass spectrum for **cysR7** (m/z 1560.845 for $[M+H]^+$).

**cysR9**

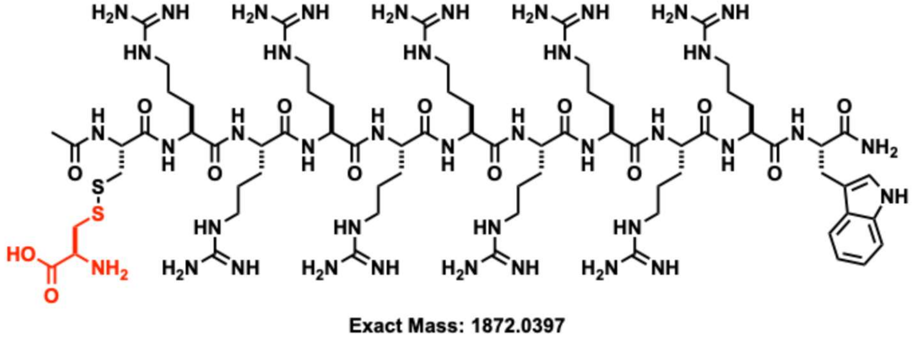


Chemical structure of **cysR9**.

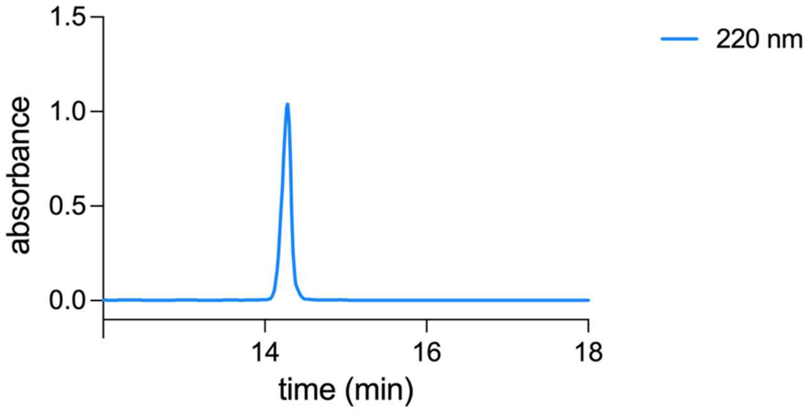


Analytical HPLC Chromatogram of **cysR9**.

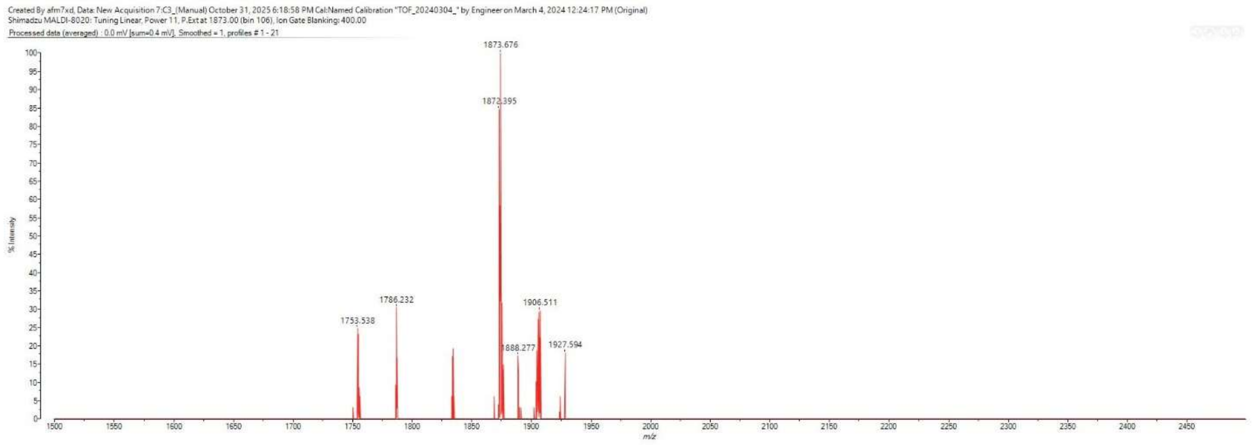


MALDI-TOF mass spectrum for **cysR9** (m/z 1873.047 for $[M+H]^+$).

**cysR11**

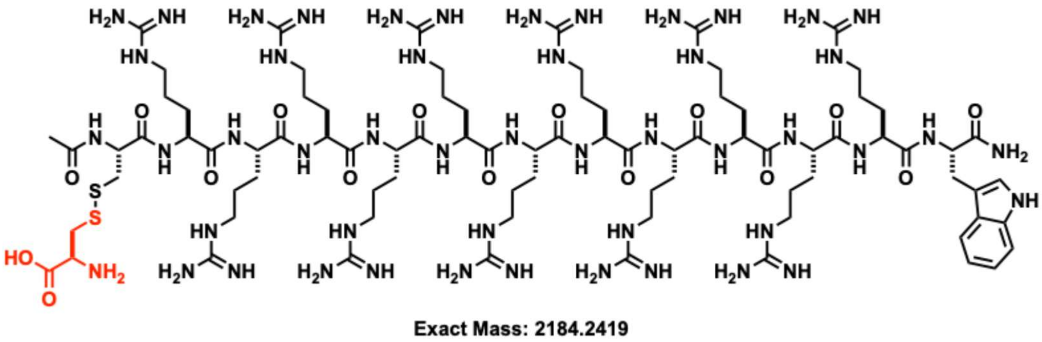


Chemical structure of **cysR11**.

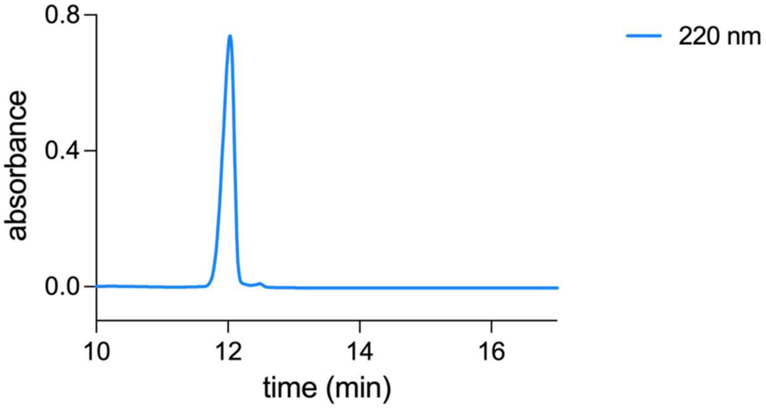


Analytical HPLC Chromatogram of **cysR11**.

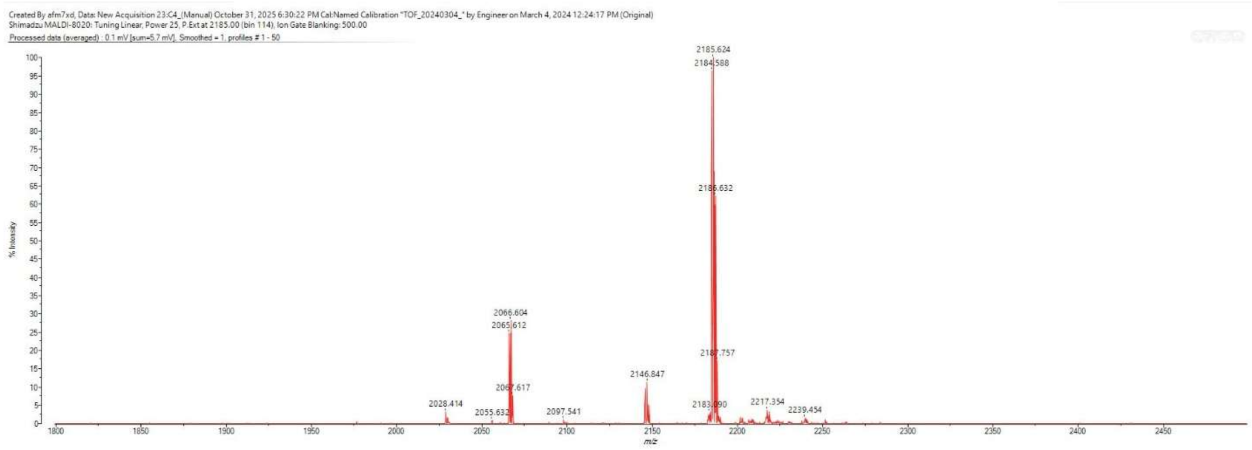


MALDI-TOF mass spectrum for **cysR11** (m/z 2185.249 for $[M+H]^+$).

**cysr7**

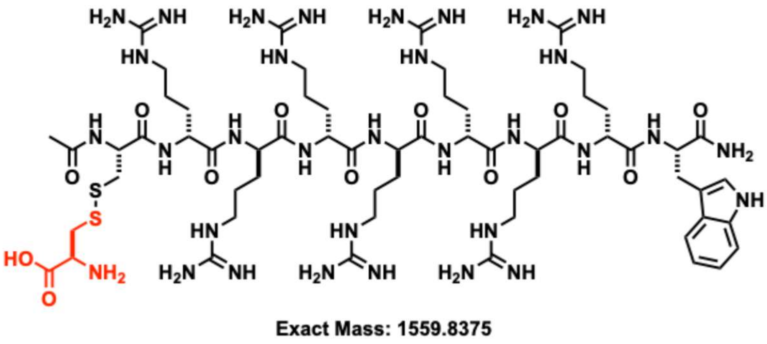


Chemical structure of **cysr7**.

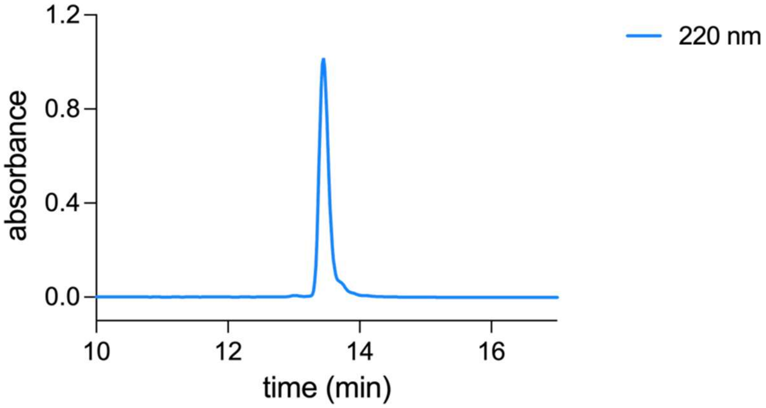


Analytical HPLC Chromatogram of **cysr7**.

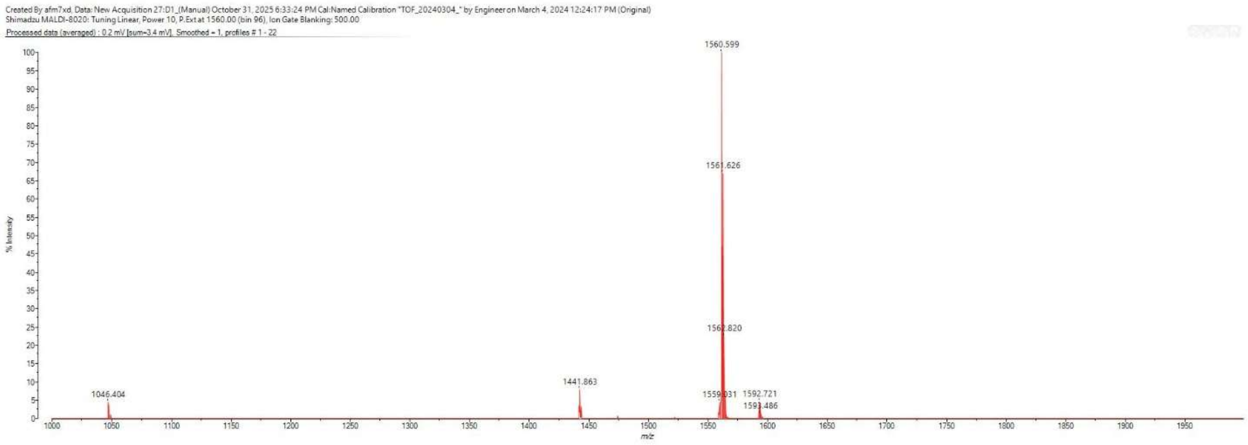


MALDI-TOF mass spectrum for **cysr7** (m/z 1560.845 for $[M+H]^+$).

**coumR7**

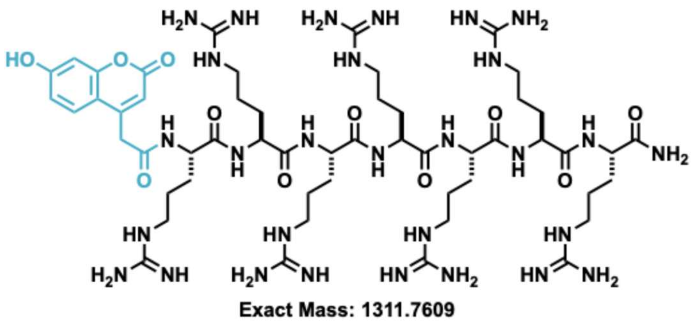


Chemical structure of **coumR7**.

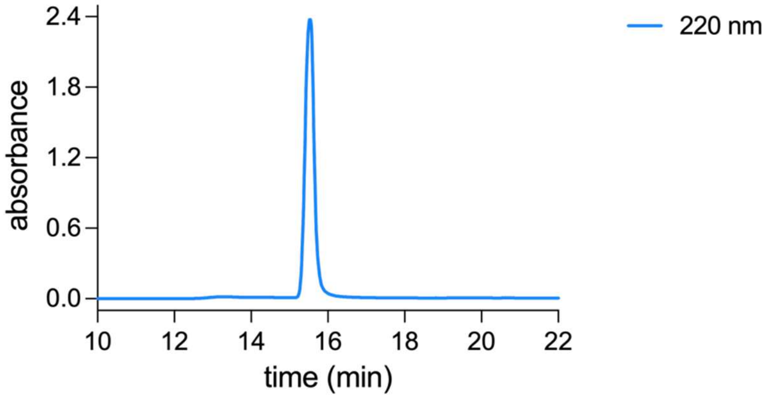


Analytical HPLC Chromatogram of **coumR7**.

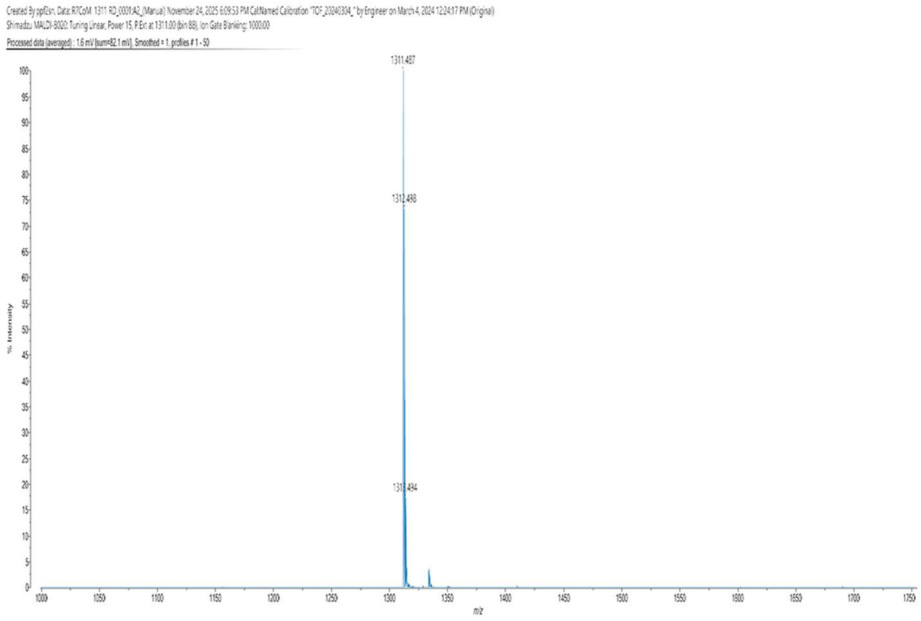


MALDI-TOF mass spectrum for **coumR7** (m/z 1312.768 for $[M+H]^+$).

**Griselimycin peptide conjugates.**

**Synthesis of cys-GM and cys-ME-GM.**

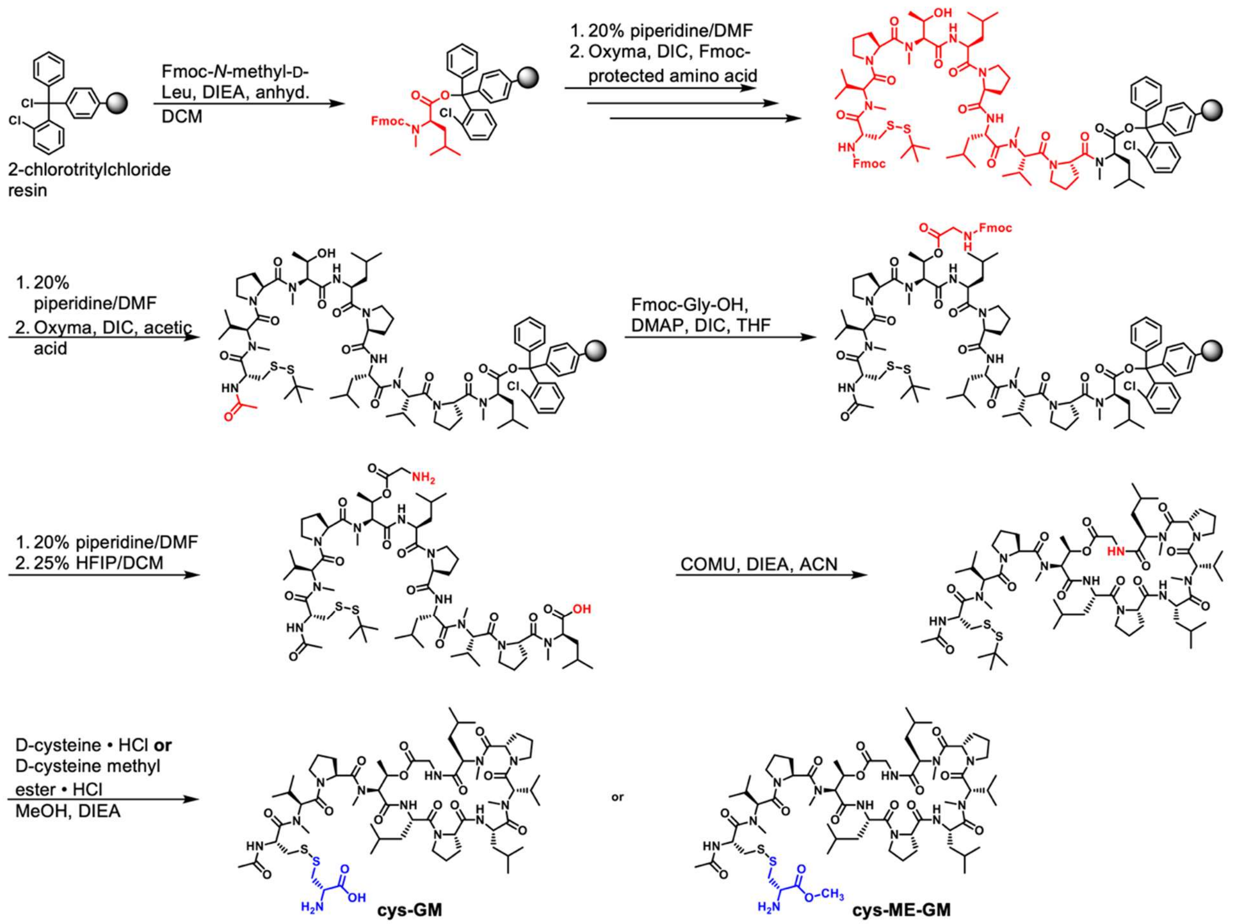


All peptides were synthesized via Fmoc-based solid-phase peptide synthesis using a previously reported route.[3] Briefly, 2-chlorotritylchloride resin (1 eq., 0.25 mmol, 1.55 mmol/g loading capacity) was added to an oven-dried peptide synthesis vessel. Fmoc-*N*-methyl-D-leucine (1.1 eq., 0.275 mmol) was dissolved in anhydrous DCM (5 mL), to which anhydrous DIEA (4.4 eq, 1 mmol) was added. This solution was added to the resin and shaken at room temperature for 1 h. The resin was washed (~5 mL each of DCM, MeOH, DCM, MeOH, DCM, DCM) and the Fmoc protecting group was removed by shaking with 10 mL of piperidine in DMF (20% v/v) for 30 minutes, then washing as described. Each sequential Fmoc-protected amino acid (5 eq., 1.25 mmol) was coupled by adding a solution of Oxyma (5 eq., 1.25 mmol) and DIC (5 eq., 1.25 mmol) in DMF (7 mL) to the resin, shaking for 2 h at room temperature, then washing and deprotecting as described. Fmoc- amino acids were added in the following order: proline, *N*-methyl-valine, leucine, proline, leucine, *N*-methyl-threonine, proline, *N*-methyl-valine. Fmoc-Cys(Stbu)-OH (5 eq., 1.25 mmol) was coupled last. The *N*-terminus was acetylated by coupling acetic acid (5 eq., 1.25 mmol) in the same manner as above. The peptide was esterified via Steglich esterification by adding Fmoc-glycine (10 eq., 2.5 mmol), DIC (10 eq., 2.5 mmol), and DMAP (0.25 eq., 0.063 mmol) dissolved in anhydrous THF (10 mL) to the resin and shaking at room temperature overnight. The Fmoc protecting group was then removed as described. Peptides were cleaved from the resin by adding a solution of HFIP in DCM (25% v/v, 10 mL) and shaking at room temperature overnight. This solution was filtered and concentrated, then dissolved in ACN (20 mL) containing DIEA (3 eq., 0.75 mmol). The peptide was cyclized by adding this solution dropwise to a solution of COMU (3 eq., 0.75 mmol) and DIEA (3 eq., 0.75 mmol) dissolved in ACN (150 mL) and stirring at room temperature overnight. This solution was concentrated and washed with hexanes (3 x 5 mL) to afford crude peptide. The crude cyclic peptide was dissolved in a minimum volume of methanol (~15 mL). DIEA (20 eq., 5 mmol) was added along with either D-cysteine HCl salt (10 eq., 2.5 mmol) for cys-GM or D-cysteine methyl ester HCl salt (10 eq., 2.5 mmol) for cys-ME-GM. This solution was shaken at room temperature for 16 hours, then concentrated to afford crude peptide. The crude peptide was purified by reverse-phased preparative high-performance liquid chromatography (RP-HPLC) equipped with Waters 1525 with 2489 UV/Visible Detector on a Phenomenex Luna 10 µm C8(2) 100 Å (250 x 21.2 mm) column using gradient elution with $H_2O/CH_3OH$ with 0.1% TFA at 10 mL/min. The HPLC fractions of the desired purified compounds were first concentrated under reduced pressure using a rotary evaporator. The final concentrated aqueous solutions were lyophilized to dryness using Labconco Freezone 4.5 L (-84 $^oC$) lyophilizer.

**cys-GM**

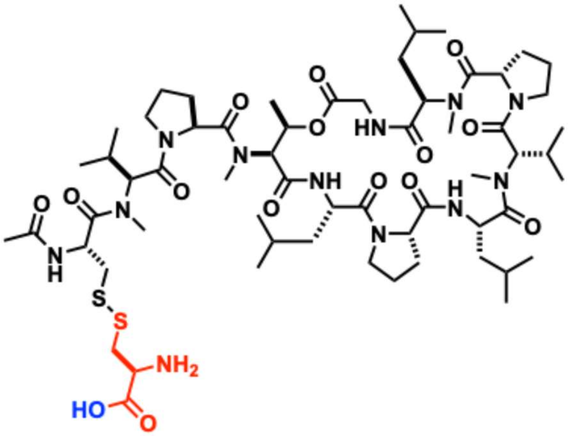

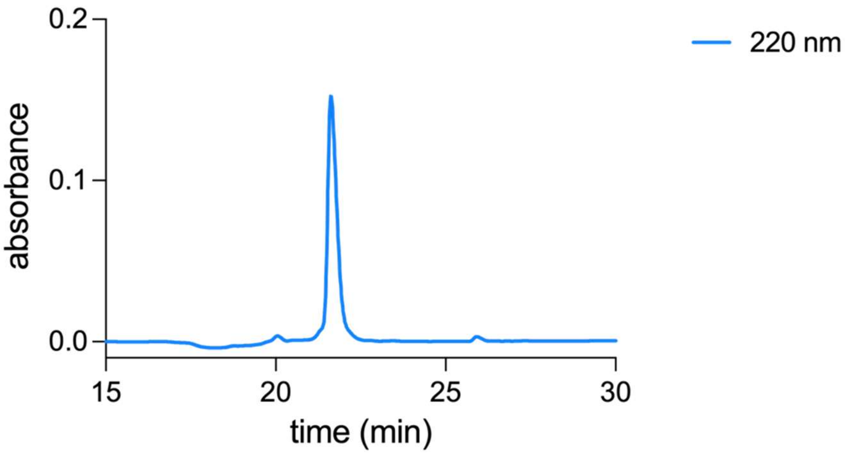


Analytical HPLC Chromatogram of **cys-GM**.

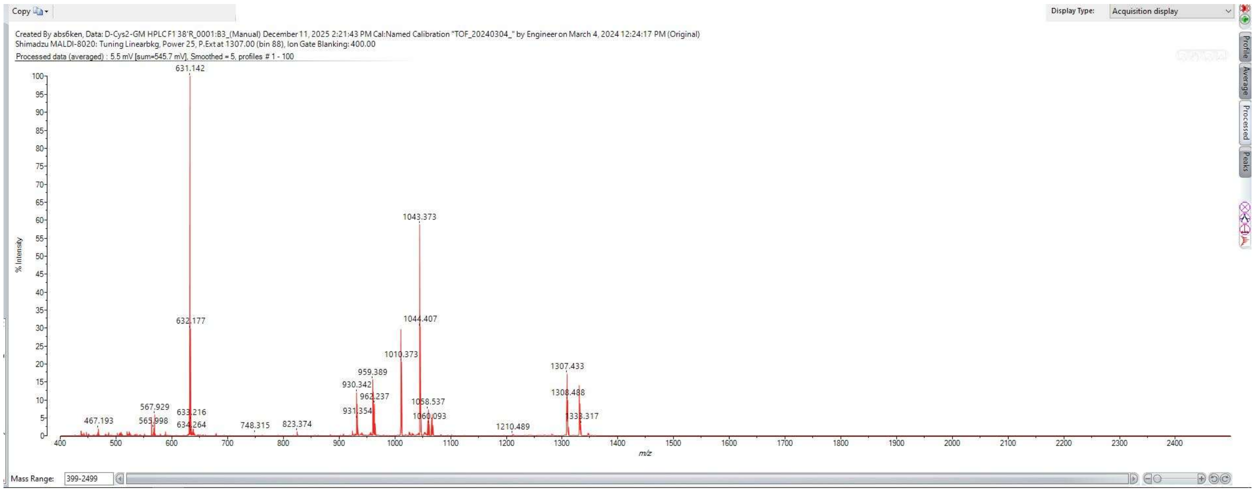


MALDI-TOF mass spectrum for **cys-GM** (m/z 1307.703 for $[M+H]^+$).

**cys-ME-GM**

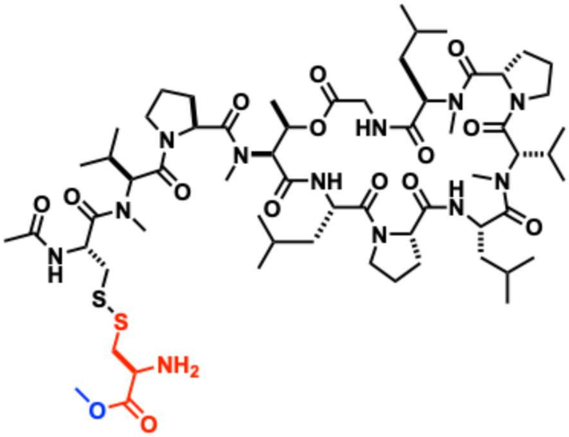

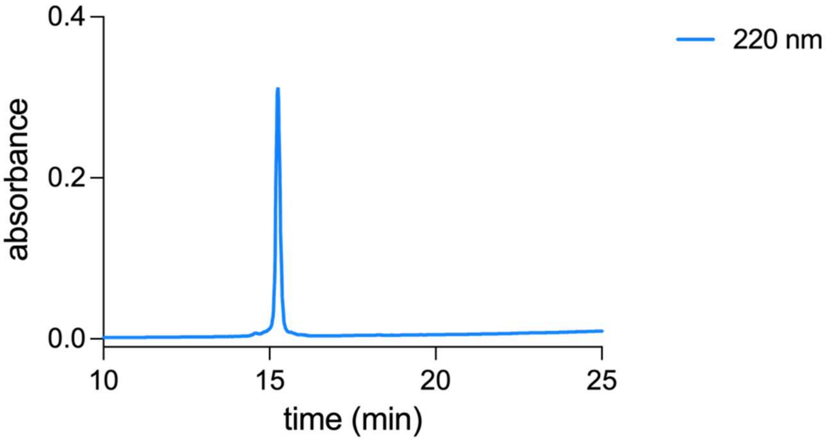


Analytical HPLC Chromatogram of **cys-ME-GM**.

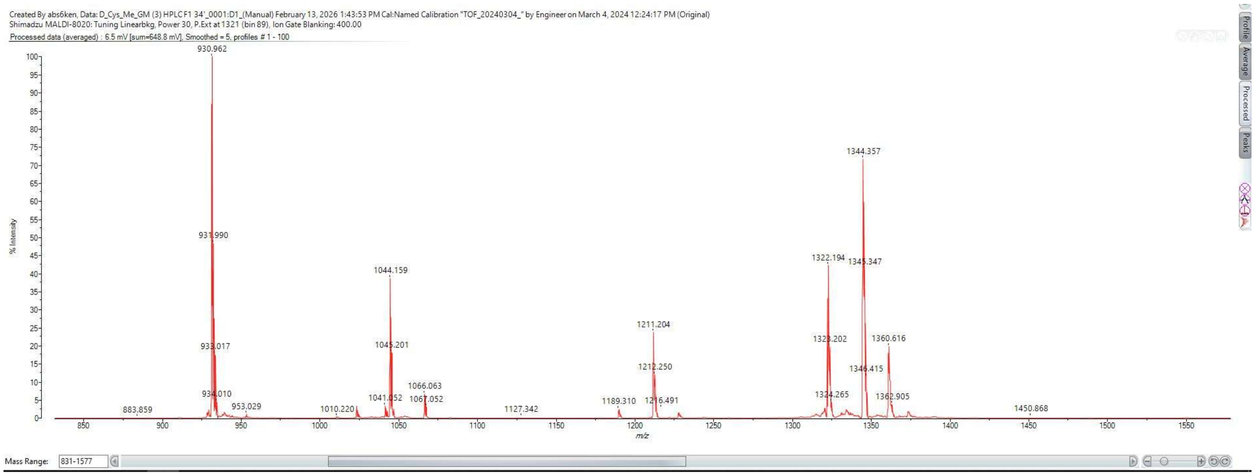


MALDI-TOF mass spectrum for **cys-ME-GM** (m/z 1321.719 for $[M+H]^+$).

### SYNTHESIS AND CHARACTERIZATION OF ANTIBIOTIC CONJUGATES

#### General Methods.

RP-HPLC was performed on instruments equipped with Waters 1525 with 2489 UV/Visible Detector on a Phenomenex Luna 10 µm C8(2) 100 Å (250 x 21.2 mm) column using gradient elution with $H_2O/CH_3OH$ with 0.1% TFA at 10 mL/min. The HPLC fractions of the desired purified compounds were first concentrated under reduced pressure using a rotary evaporator. The final concentrated aqueous solutions were lyophilized to dryness using Labconco Freezone 4.5 L (-84 $^{o}$C) lyophilizer. The peptides were analyzed for purity using Phenomenex Luna 5 µm C8(2) on the same RP-HPLC; gradient elution in $H_2O/CH_3OH$ with 0.1% TFA at 1 mL/min. Electrospray ionization (ESI) mass spectrometry was performed on an Advion Expression® CMS using standard parameters for intermediate compounds. A low-fragmentation, low-energy setup was employed for fragmentation-sensitive samples. High-resolution mass spectrometry (HRMS) analyses were conducted using a Luna C18(2) column (5 µm, 100 Å, 250 × 4.6 mm; Phenomenex) coupled to an Agilent 1260 Infinity II Prime LC system and an Agilent 6545B QTOF mass spectrometer. Data were acquired and processed using MassHunter software (Agilent Technologies). UV–visible spectra were recorded on a Genesys 50 UV–Vis spectrophotometer (Thermo Scientific). $^{1}$H and $^{13}$C NMR spectra of final compounds and intermediates were obtained on a Varian 600 MHz spectrometer. All spectra were processed and analyzed using MestreNova software. Residual solvent signals from deuterated solvents were used as an internal standard with reference to tetramethylsilane (TMS) for defining chemical shifts. Chemical shifts are reported in parts per million or ppm ($\delta$) and coupling constants (J) are reported in hertz (Hz).

**D-cystine-ciprofloxacin-methyl-ester (cys-Cipro).**

**Synthesis of D-cystine-ciprofloxacin-methyl-ester (cys-Cipro).**

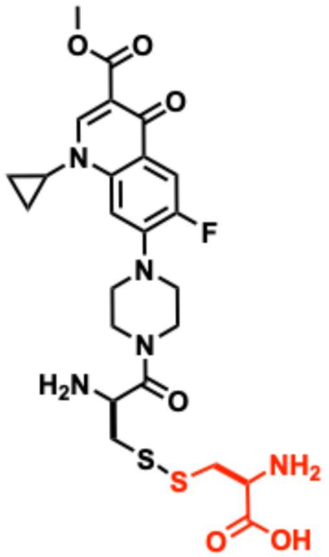


Synthesis of the ciprofloxacin-methyl-ester was carried out following a previously reported procedure in the literature.[4] Column chromatography on silica gel (Supelco, 60 Å, 230–400 mesh, 40–63 µm particle size) was performed using manually packed columns and solvent systems selected based on compound polarity. The purified compound was characterized by mass spectrometry and $^1$H NMR spectroscopy.

MF: $C_{19}H_{22}FN_3O_3$, calcd. m/z 346.15 obs. 346.1 for $[M+H]^+$. $^1$H-NMR: ($CDCl_3$) d 1.14 (m, 2H, cyclopropyl-$CH_2$), 1.32 (m, 2H, cyclopropyl-$CH_2$), 3.12 (m, 4H, piperidyl 2 x $CH_2$), 3.27 (m, 4H, piperidyl 2 x $CH_2$), 3.43 (m, 1H, cyclopropyl CH-N), 3.91 (s, 3H, $OCH_3$), 7.26 (d, *J* = 6 Hz, 1H, ArH), 8.02 (d, *J* = 18 Hz, 1H, ArH), 8.54 (s, 1H, ArH).

Commercially available *N*-$^t$Boc-D-Cys (200 mg, 0.905 mmol) was then oxidized to *N,N'*-di-$^t$Boc-D-cystine by dissolving it in a solution of $CH_3CN$ (22.5 mL) and 0.5 M ammonium bicarbonate (7.5 mL). The solution was stirred at room temperature (~18 h) while being continuously bubbled with air. Completion of oxidation was confirmed by Ellman's assay, which indicated the absence of free thiols. The reaction mixture was then concentrated under reduced pressure and lyophilized to dryness, yielding *N*-di-$^t$Boc-D-cystine as a white solid (185 mg, 97%), which was used after removal of ammonium bicarbonate. Conjugation of *N*-di-$^t$Boc-D-cystine to ciprofloxacin-methyl-ester was performed as follows. A solution of *N*-di-$^t$Boc-D-cystine (135 mg, 0.307 mmol) in 10–15 mL dimethylformamide (DMF) was treated with ciprofloxacin-methyl-ester (50 mg, 0.145 mmol), 4-dimethylaminopyridine (DMAP, 4 mg, 0.03 mmol), and 1-ethyl-3-(3-dimethylaminopropyl)carbodiimide (EDC, 25 mg, 0.130 mmol). The mixture was stirred at room temperature, and reaction progress was monitored by thin-layer chromatography (TLC) and electrospray ionization (ESI) mass spectrometry. After 24 h, the reaction was subjected to liquid–liquid extraction using $H_2O$ and DCM to remove DMF and water-soluble byproducts. The organic layer was separated, washed with 0.1 M HCl, dried over anhydrous $Na_2SO_4$, and concentrated under reduced pressure to afford a crude residue. Purification was carried out by column chromatography on silica gel (Supelco, 60 Å, 230–400 mesh, 40–63 µm) using chloroform followed by an increasing gradient of $CHCl_3$: $CH_3OH$. Fractions corresponding to *N*-di-$^t$Boc-D-cystine conjugated to ciprofloxacin-methyl-ester were collected and combined. These fractions were concentrated and subjected to tBoc deprotection with TFA (0.5 mL) and DCM (0.5 mL) for 2 h at room temperature. The reaction mixture was concentrated under reduced pressure and triturated with diethyl ether to remove TFA, yielding a crude solid. The product was further purified by reverse-phase HPLC. Fractions containing the desired product (based on the expected m/z) were collected and concentrated under reduced pressure to yield a transparent residue, which was lyophilized to give a white solid (21 mg). Minor amounts of the expected bis-ciprofloxacin side product were observed in later fractions. The desired conjugate was characterized by $^1$H NMR and mass spectrometry before use in accumulation assays. A stock solution was prepared by dissolving the compound in Milli-Q $H_2O$ and analyzed for purity by analytical RP-HPLC. Broad peaks observed in the NMR spectrum were attributed to H/D exchange.

$^1$H-NMR: ($D_2O$) $\delta$ 1.14 (m, 2H, cyclopropyl-$CH_2$), 1.32 (m, 2H, cyclopropyl-$CH_2$), 1.39-1.49 (m, 6H, 3 x $CH_2$), 1.49 (s, 9H, 3 x$CH_3$), 1.58 (dt, *J*=6Hz, 128Hz, 2H, $CH_2$), 1.74 (dt, *J*=6, 12Hz, 2H, $CH_2$), 3.22 (t, , *J*=6Hz, 4H, 2 x $CH_2$), 3.43 (m, 1H, CH), 3.45-3.51 (m, 4H, piperidyl 2 x CH2), 3.62 (m, 2H), 3.63-3.67 (m, 4H, 2 x $CH_2$), 7.31 (d, *J* = 6 Hz, 1H, ArH), 8.03 (d, *J* = 12 Hz, 1H, ArH), 8.81 (s, 1H, ArH), 10.1 (brs, 1H, NH). $^{13}$C-NMR: ($CDCl_3$) d 8.2, 25.4, 26.8, 28.4, 29.5, 32.6, 34.7, 39.2, 45.1, 50.0, 70.0, 70.2, 70.6, 71.3, 105.0, 111.4, 112.7, 112.8, 122.2, 138.4, 144.8, 146.8, 152.6, 154.2, 154.6, 165.1, 175.4. HRMS-QTOF: for MF; $C_{24}H_{31}FN_5{}_4O_6S_2$ calcd. m/z 568.1694 obs. 568.1699 for $[M+H]^+$

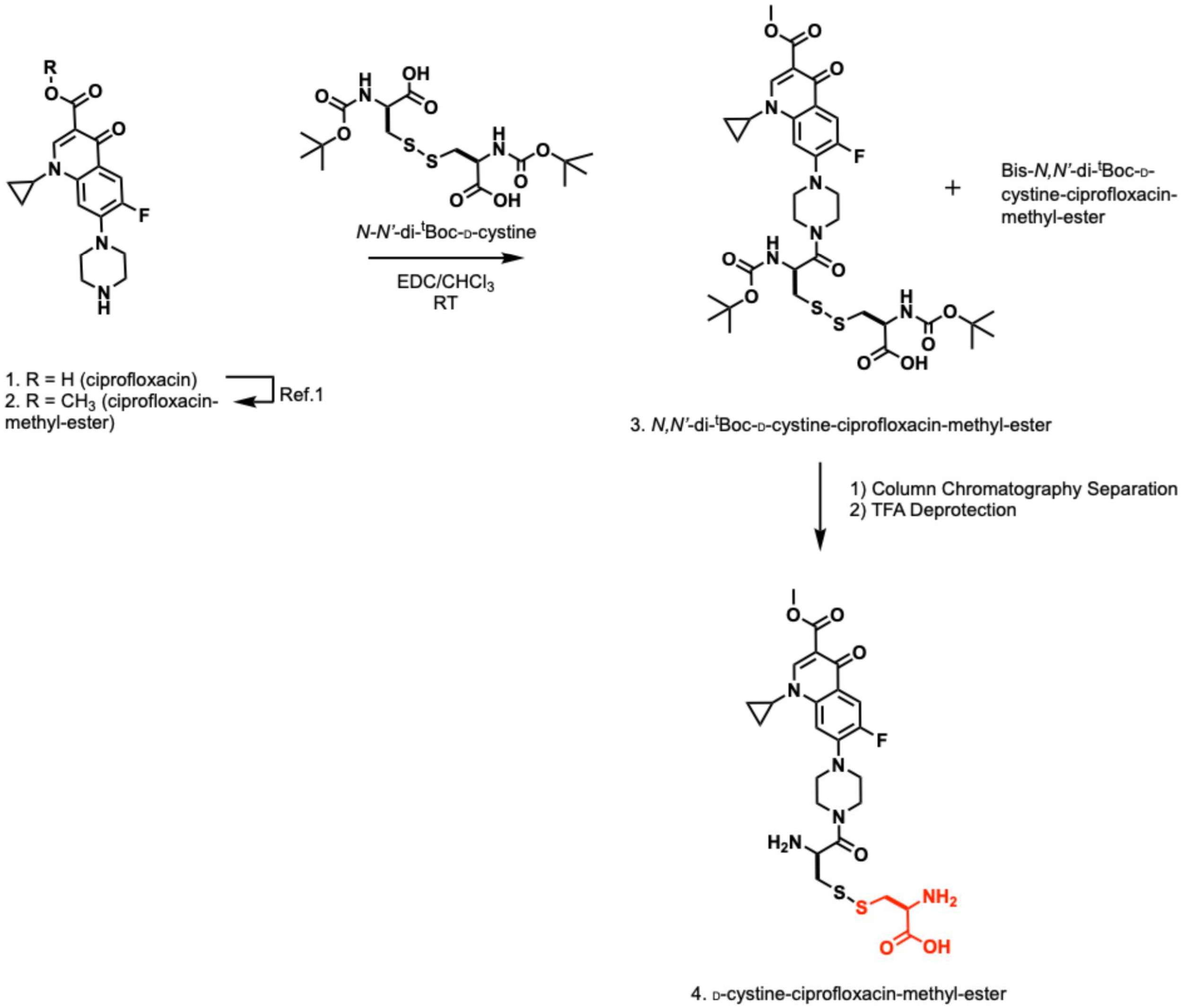


Synthesis scheme of D-cystine-ciprofloxacin-methyl-ester (**cys-Cipro**).

**Characterization of D-cystine-ciprofloxacin-methyl-ester (cys-Cipro).**

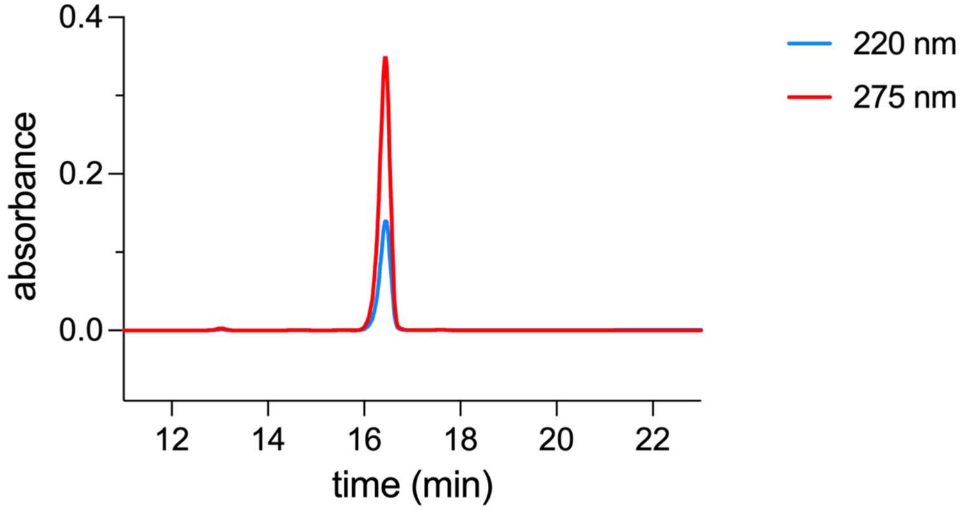


Analytical HPLC Chromatogram of **D-cystine-ciprofloxacin-methyl-ester** (**cys-Cipro**).

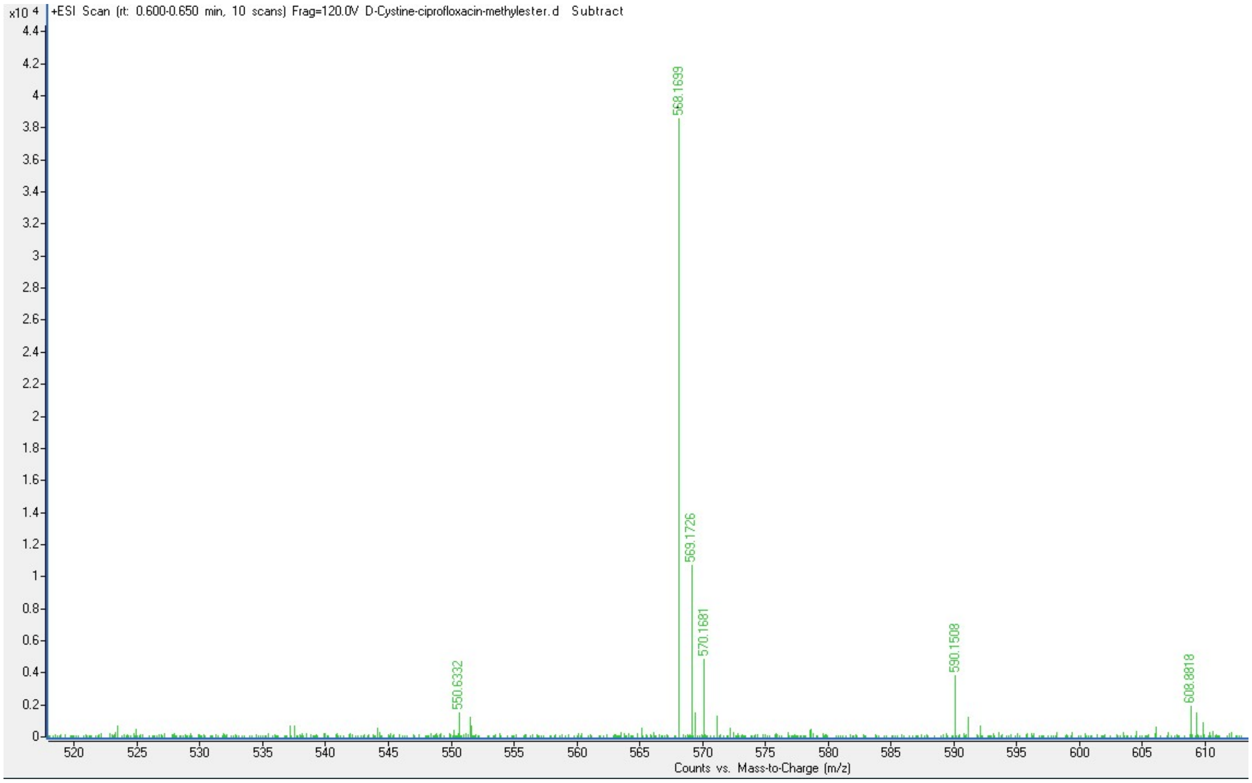


QTOF High Resolution Mass Spectrum for **D-cystine-ciprofloxacin-methyl-ester** (**cys-Cipro**) (m/z 568 for $[M+H]^+$).

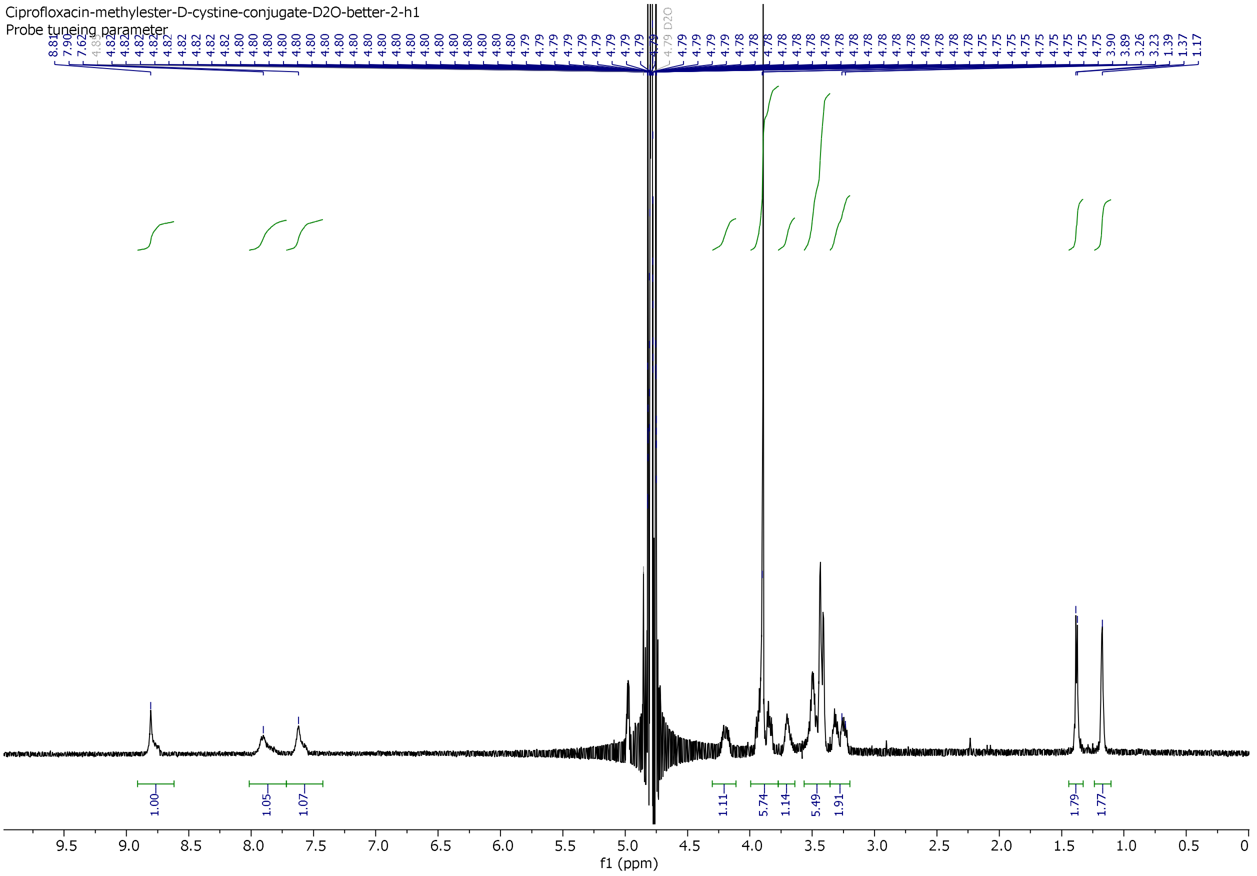


[1]H NMR spectrum of **D-cystine-ciprofloxacin-methyl-ester** (**cys-Cipro**).

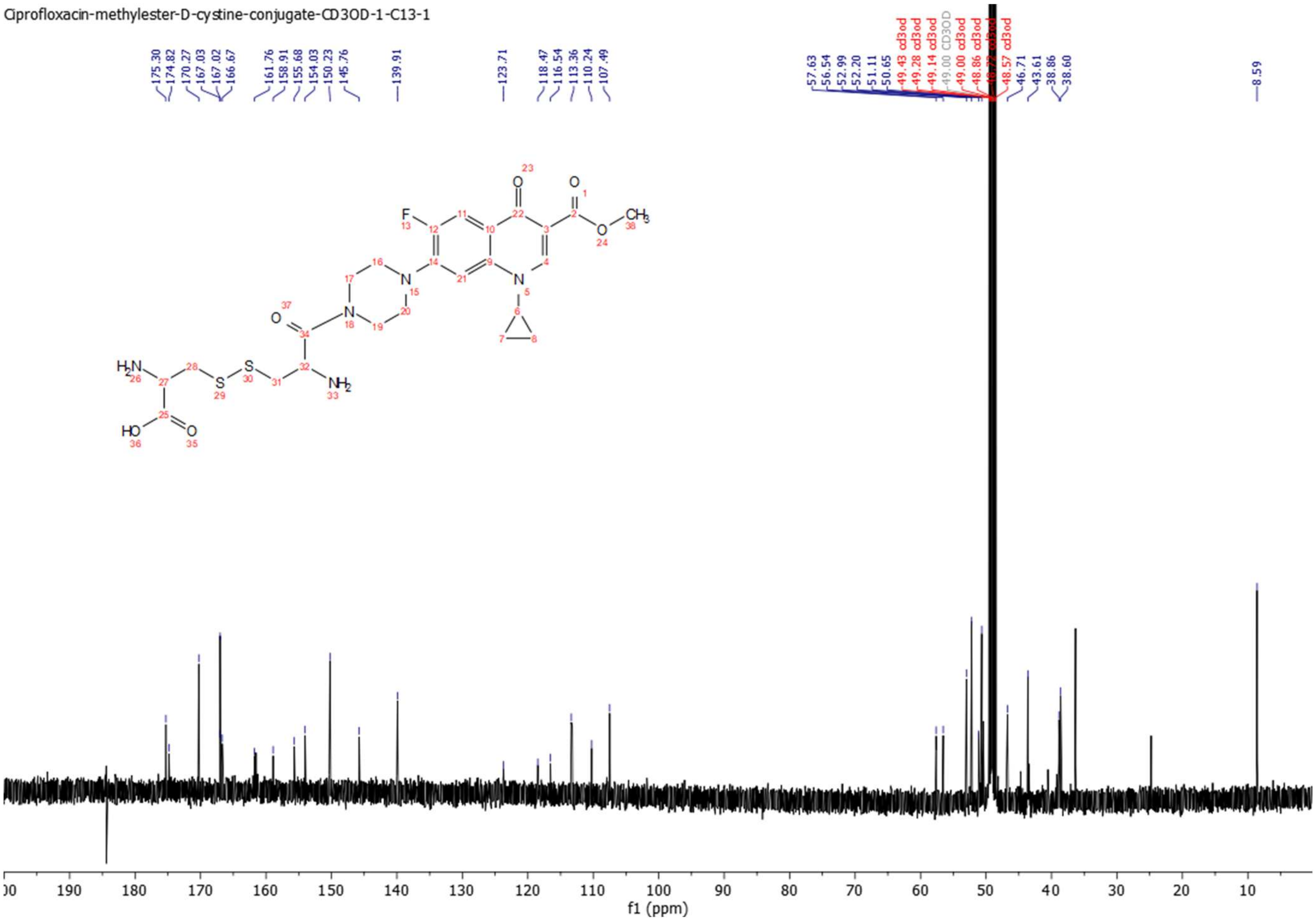


$^{13}$C NMR Spectrum of **D-cystine-ciprofloxacin-methyl-ester** (**cys-Cipro**).

**Linezolid-cystamine-D-cysteine disulfide (cys-Line).**

**Synthesis of linezolid-cystamine-D-cysteine disulfide (cys-Line).**

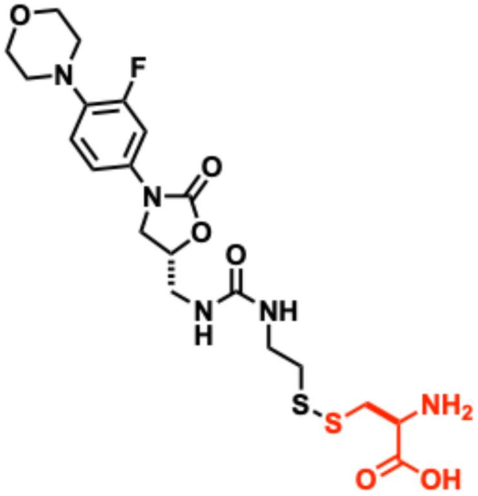

Compound linezolid-cystamine-D-cysteine disulfide (**cys-Line**) was synthesized as outlined in the synthetic scheme below. Linezolid amine (30.0 mg, 0.10 mmol) was dissolved in anhydrous dichloromethane (3 mL), followed by addition of carbonyldiimidazole (18 mg, 0.11 mmol) and DIEA (10 µL). The homogeneous solution was stirred at room temperature overnight. The reaction mixture was then concentrated under reduced pressure, and a solution of cystamine–pyridine disulfide hydrochloride (23.5 mg, 0.10 mmol) in anhydrous MeCN (3 mL) containing DIEA (20 µL) was added. The mixture was stirred at room temperature for 4 h, during which reaction progress was monitored by ESI–MS, confirming consumption of the starting material and formation of the linezolid–pyridyl–cystamine disulfide intermediate. The reaction mixture was concentrated under reduced pressure and the residue redissolved in $CH_3OH$. An aqueous solution of D-cysteine (25.0 mg, 0.20 mmol in 2.0 mL deionized $H_2O$) was then added, and the solution was stirred for 2 h at room temperature. ESI–MS analysis confirmed formation of the desired linezolid–cystamine–D-cysteine disulfide product. The reaction mixture was concentrated under reduced pressure and dissolved in $CH_3CN/H_2O$ (1:3, 10 mL) for purification by RP-HPLC. Fractions corresponding to the desired product (based on m/z) were collected and concentrated under reduced pressure to yield a colorless solid (27.7 mg, 53%). The product was characterized by $^1H$ and $^{13}C$ NMR spectroscopy and mass spectrometry prior to use in accumulation assays. A stock solution was prepared by dissolving the purified compound in Milli-Q $H_2O$, and its purity was confirmed by analytical RP-HPLC.

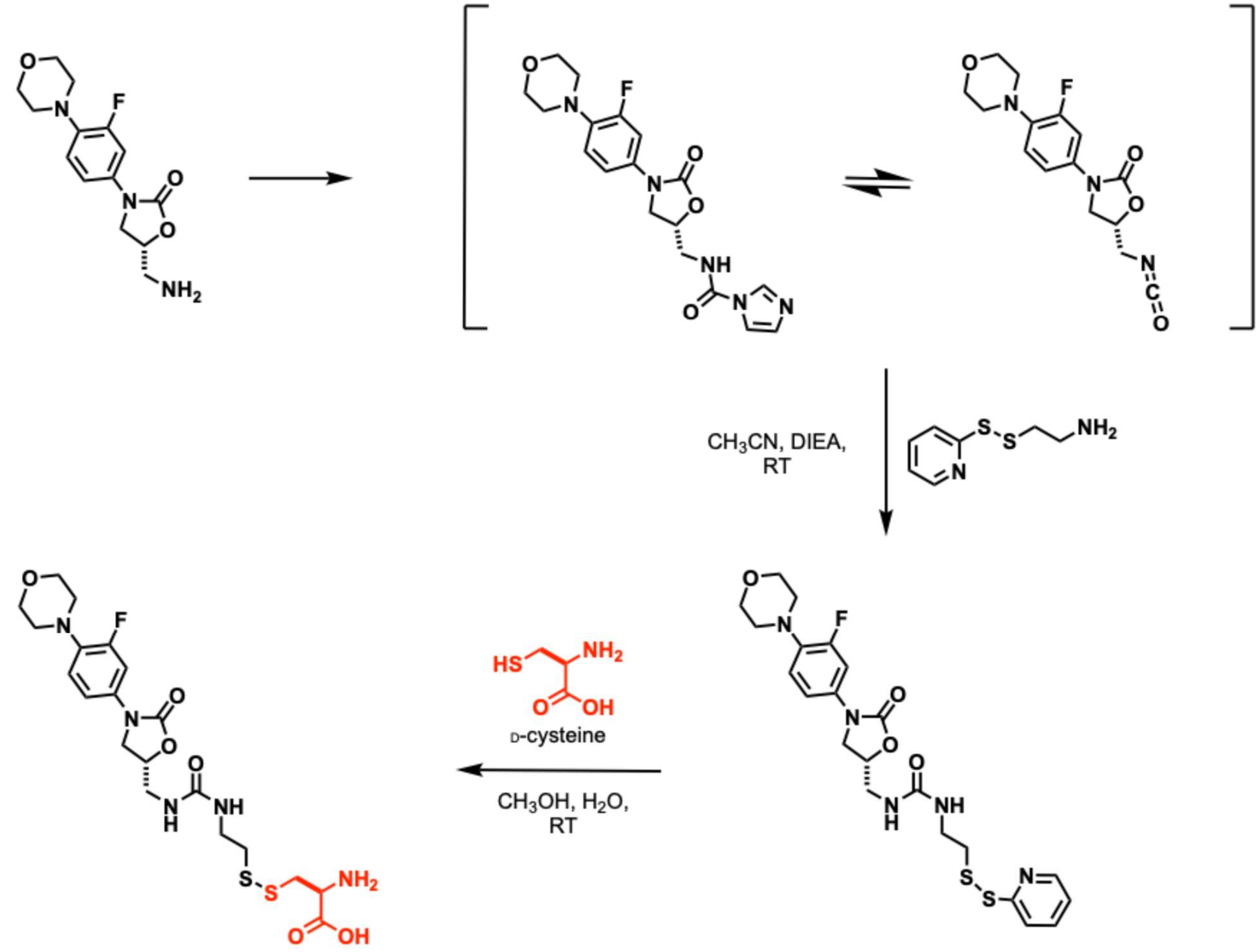


Synthesis scheme of linezolid-cystamine-D-cysteine disulfide (**cys-Line**).

**Characterization of linezolid-cystamine-D-cysteine disulfide (cys-Line).**

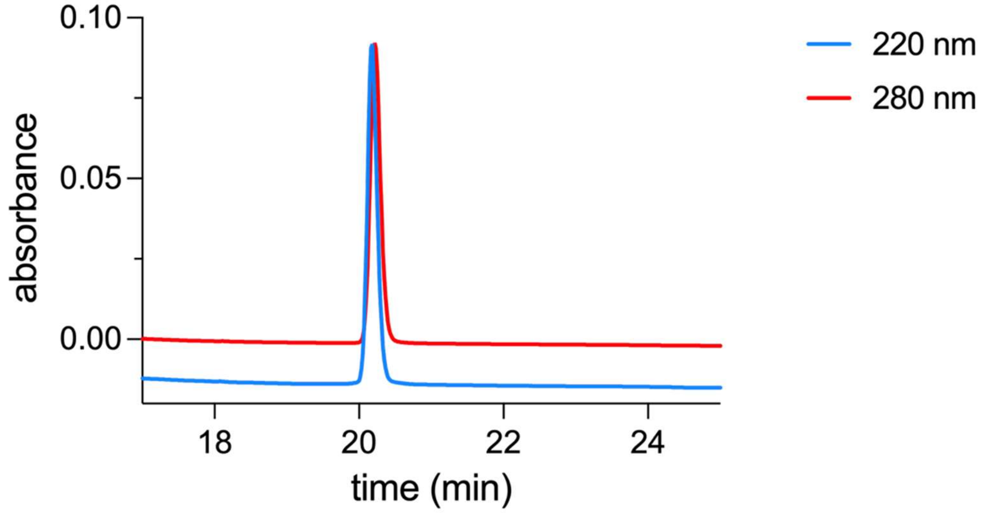


Analytical HPLC chromatogram of linezolid-cystamine-D-cysteine disulfide (**cys-Line**).

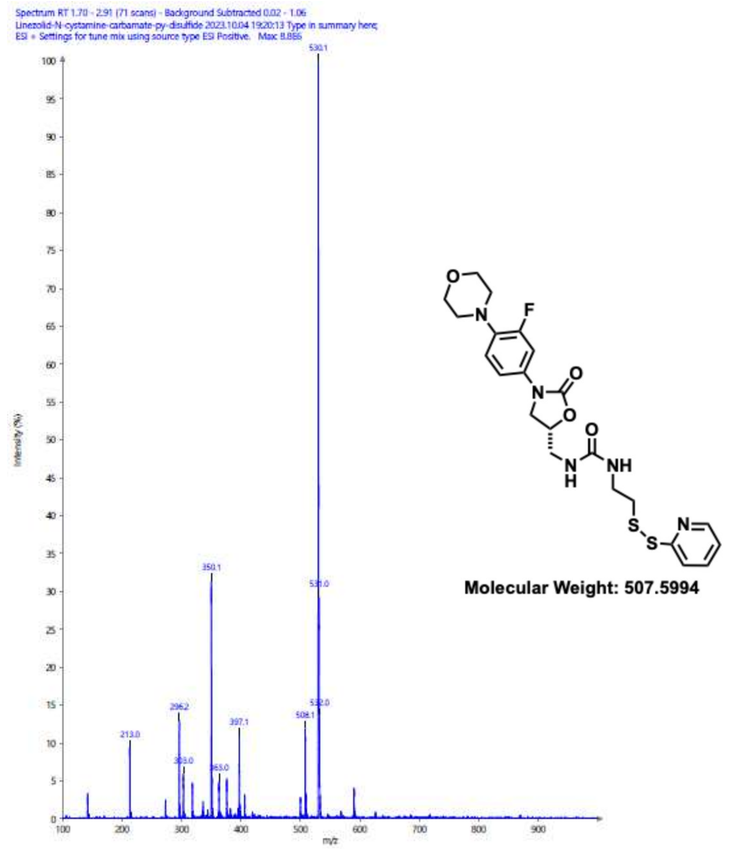


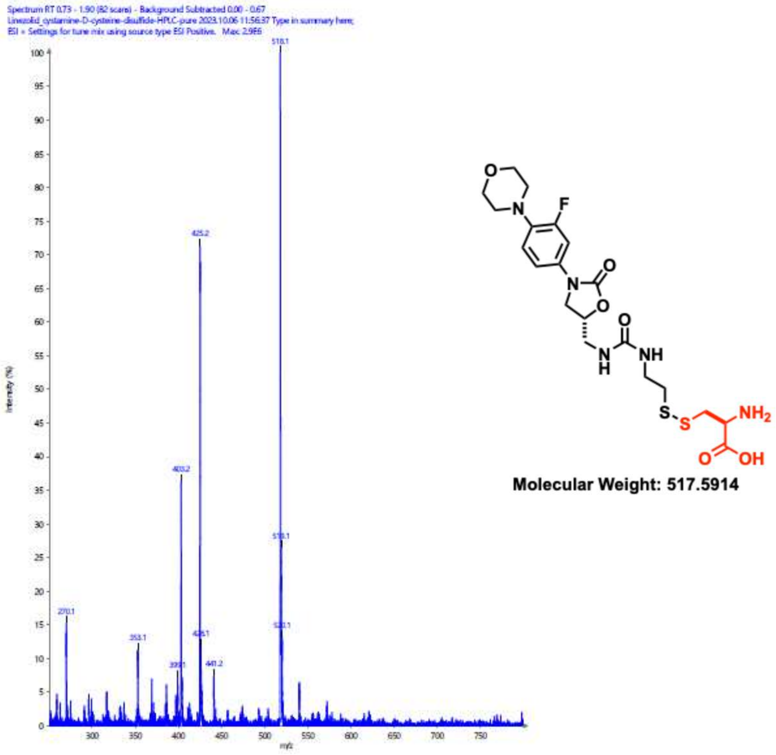


ESI-Mass spectra for intermediate (m/z 508 for [M+H]$^+$) and linezolid-cystamine-D-cysteine disulfide (**cys-Line**) (m/z 518 for [M+H]$^+$).

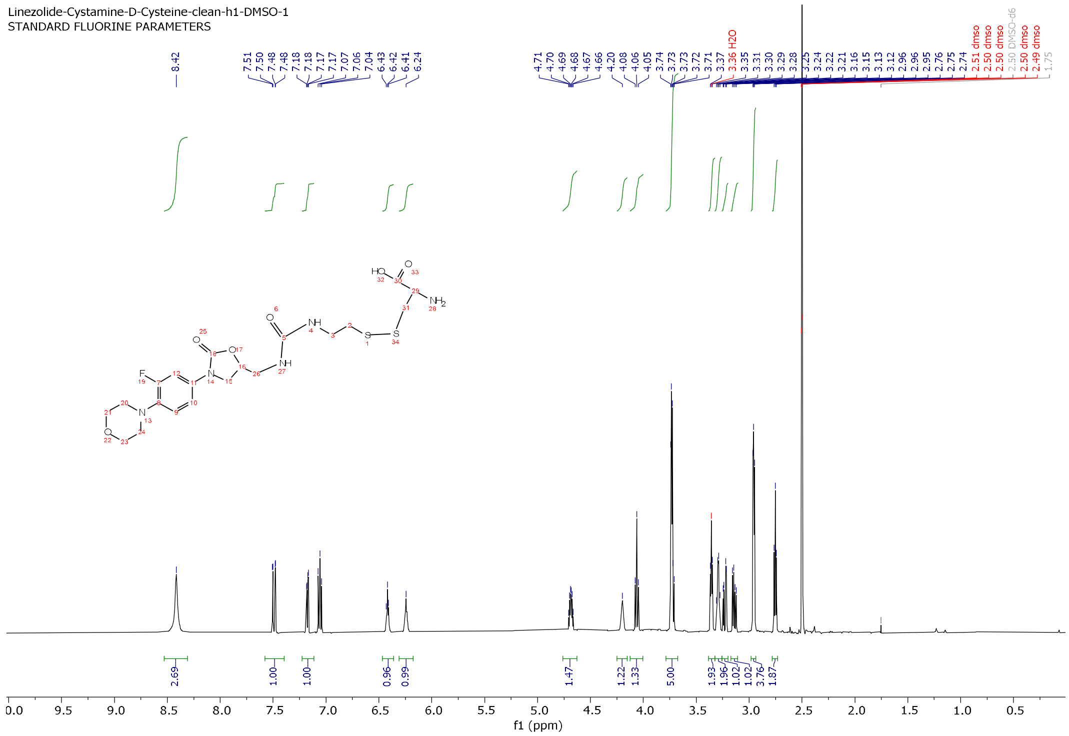


$^{1}$H NMR spectrum of linezolid-cystamine-D-cysteine disulfide (**cys-Line**).

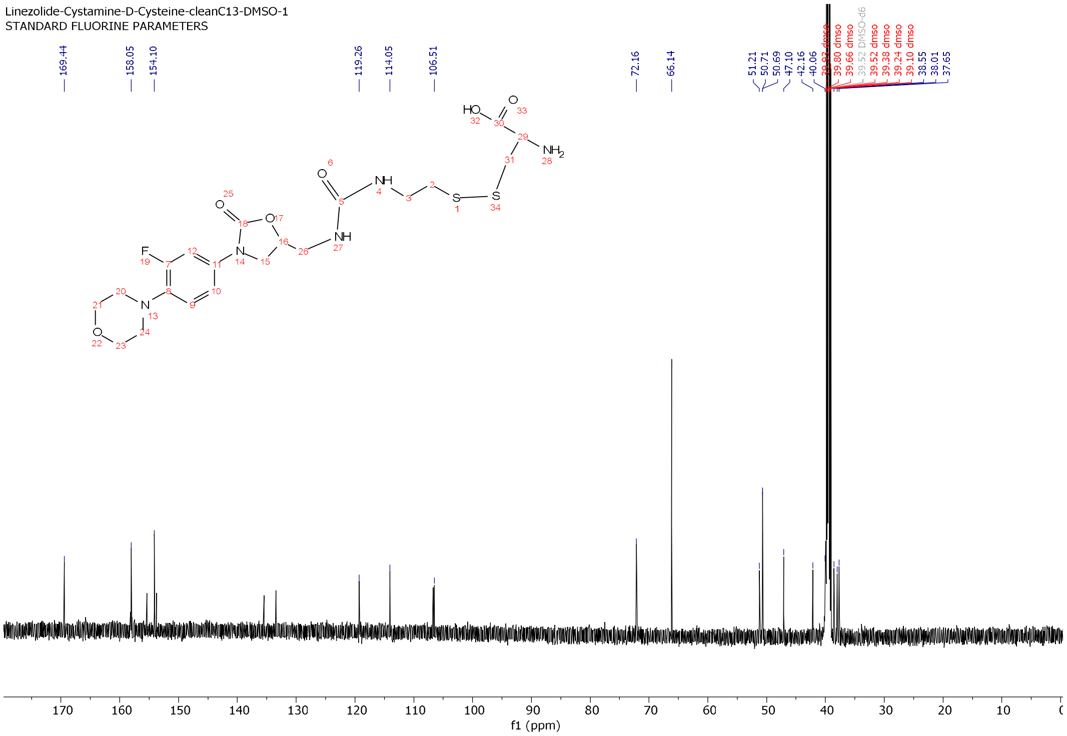


$^{13}$C NMR Spectrum of linezolid-cystamine-D-cysteine disulfide (**cys-Line**).

**Puromycin-cystamine-D-cysteine disulfide (cys-Puro).**

**Synthesis of puromycin-cystamine-D-cysteine disulfide (cys-Puro).**

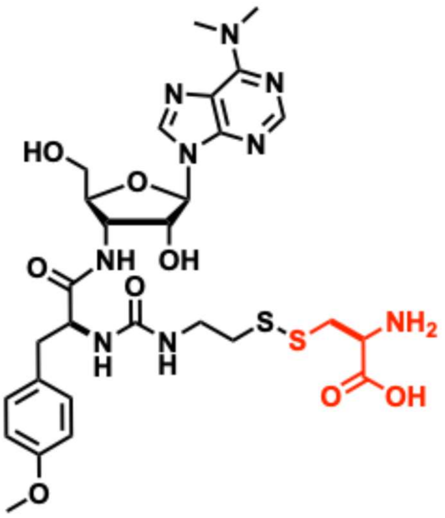

Synthesis of puromycin-cystamine-D-cysteine disulfide (**cys-Puro**) was carried out as outlined in the synthetic scheme below. Puromycin dihydrochloride (27.4 mg, 0.05 mmol) was dissolved in anhydrous DMF (2 mL), followed by addition of preformed 2-((2-isocyanatoethyl)disulfaneyl)pyridine (20.0 mg); prepared from carbonyldiimidazole and 2-(pyridine-2-yldisulfaneyl)ethan-1-amine), followed by DIEA (10 µL). The resulting homogeneous mixture was stirred at room temperature overnight. The following day, ESI–MS confirmed formation of the intermediate pyridyl disulfide conjugate. The reaction mixture was then triturated with diethyl ether (3 × 5 mL) to remove residual DMF. The resulting residue was redissolved in $CH_3OH$, and an aqueous solution of D-cysteine (75.0 mg, 0.4 mmol in 2.0 mL deionized $H_2O$) was added. The mixture was stirred at room temperature for 2 h, during which ESI–MS analysis confirmed formation of the desired puromycin-cystamine-D-cysteine disulfide (m/z 694 $[M+H]^+$). The reaction mixture was concentrated under reduced pressure to remove $CH_3OH$, and the residue was dissolved in $CH_3CN/H_2O$ (1:4, 10 mL). The crude product was purified by preparative RP-HPLC on a C18 silica column. Fractions containing the desired compound, as identified by ESI–MS, were pooled and concentrated to remove $CH_3CN$. The remaining aqueous solution was frozen at –80 °C for 2 h and lyophilized to yield a colorless solid (23.8 mg). The compound was isolated as a trifluoroacetate salt and was characterized by mass spectrometry, $^1$H-NMR spectroscopy. The stock solution was then analyzed for purity by analytical RP-HPLC. The spectroscopic data was in agreement with the assigned structure.

$^1$H-NMR: (DMSO-$d_6$) d 2.69-2.74 (m, 3H, -$CH_2$), 2.86 (dd, 1H, *J* = 6 and 12 Hz, CH2), 3.13 (dd, 1H, *J* = 6 and 12 Hz, -CH2), 3.20-3.25 (m, 2H, -CH2), 3.46 (dd, 1H, *J* = 2 and 6 Hz, -CH), 3.69 (dd, 1H, *J* = 6 and 18 Hz, -$CH_2$), 3.92 (m, 1H), 4.19 (brs, 1H), 4.42-4.85 (m, 2H), 4.98 (d, *J* = 2 Hz, 1H), 6.21 (d, *J* = 12 Hz, 1H), 6.33 (brs, 1H), 6.82 (d, *J* = 12 Hz, 2H, ArH), 7.11 (d, *J* = 12 Hz, 2H, ArH), 8.11 (d, 1H, *J* = 6 Hz, ArH), 8.25 (s, ArH),

8.41 (brs, 1H) 8.45 (s, 1H). HRMS-QTOF: for MF; $C_{24}H_{31}FN5_4O_6S_2$ calcd. m/z 568.1694 obs. 568.1699 for $[M+H]^+$

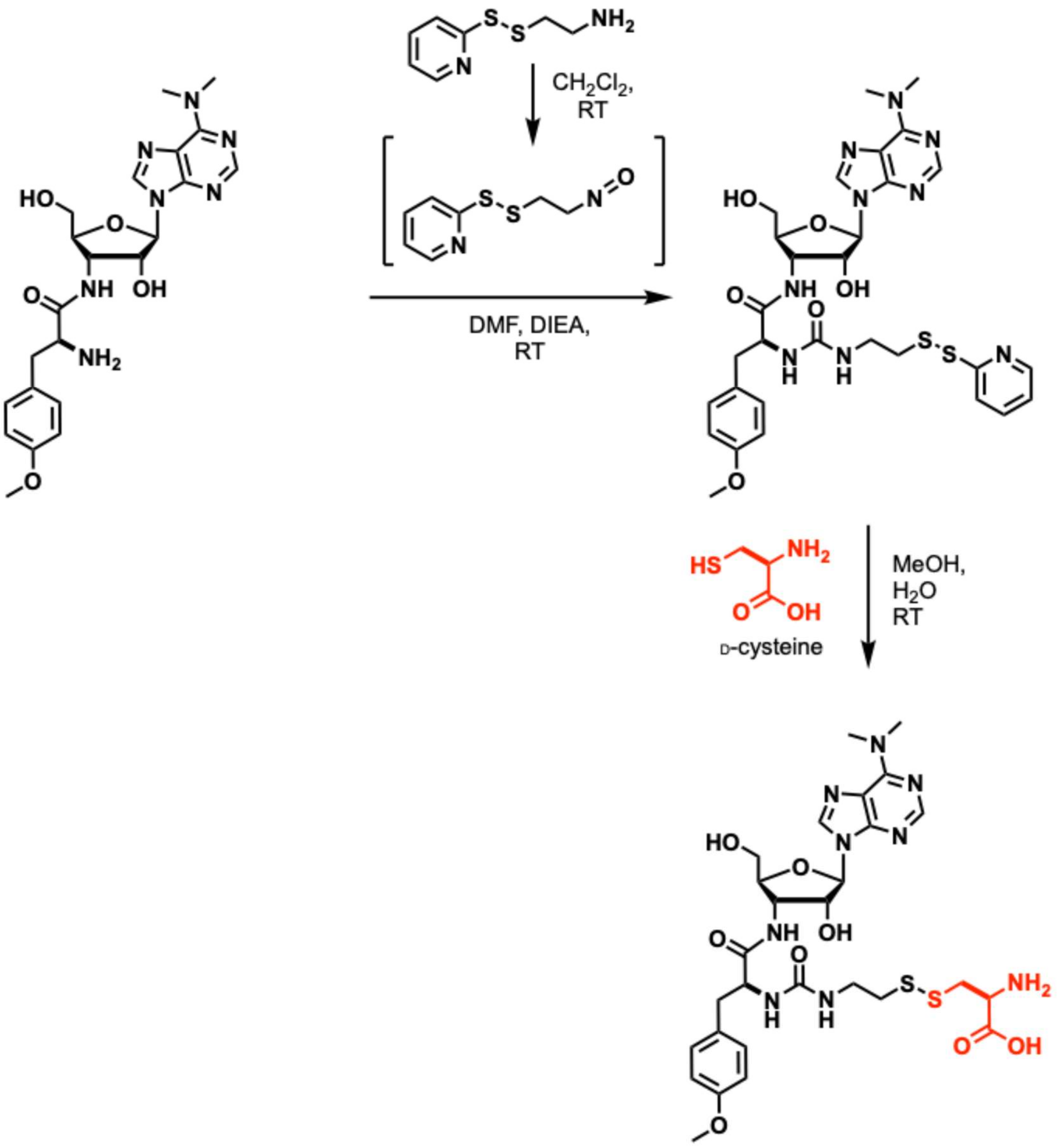


Synthesis scheme for puromycin-cystamine-D-cysteine disulfide (**cys-Puro**).

**Characterization of puromycin-cystamine-D-cysteine disulfide (cys-Puro).**

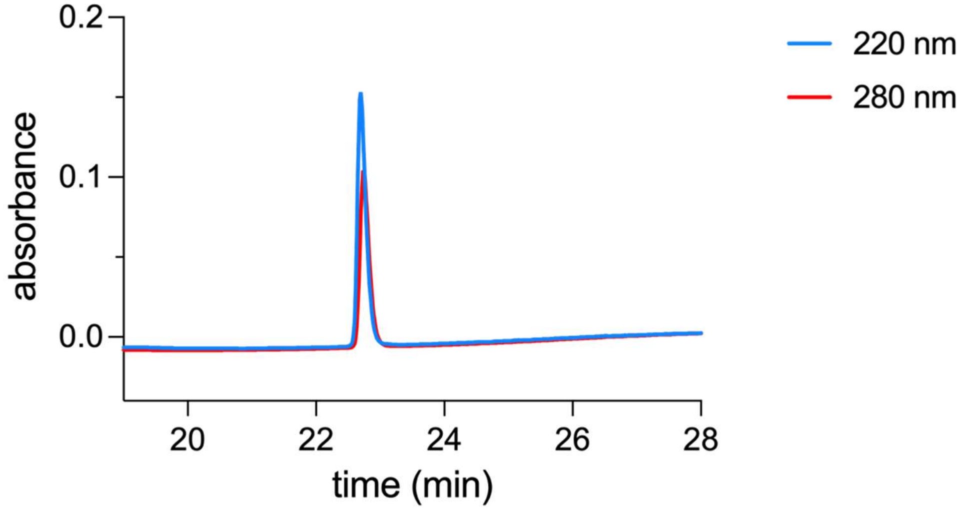


Analytical HPLC chromatogram of puromycin-cystamine-D-cysteine disulfide (**cys-Puro**).

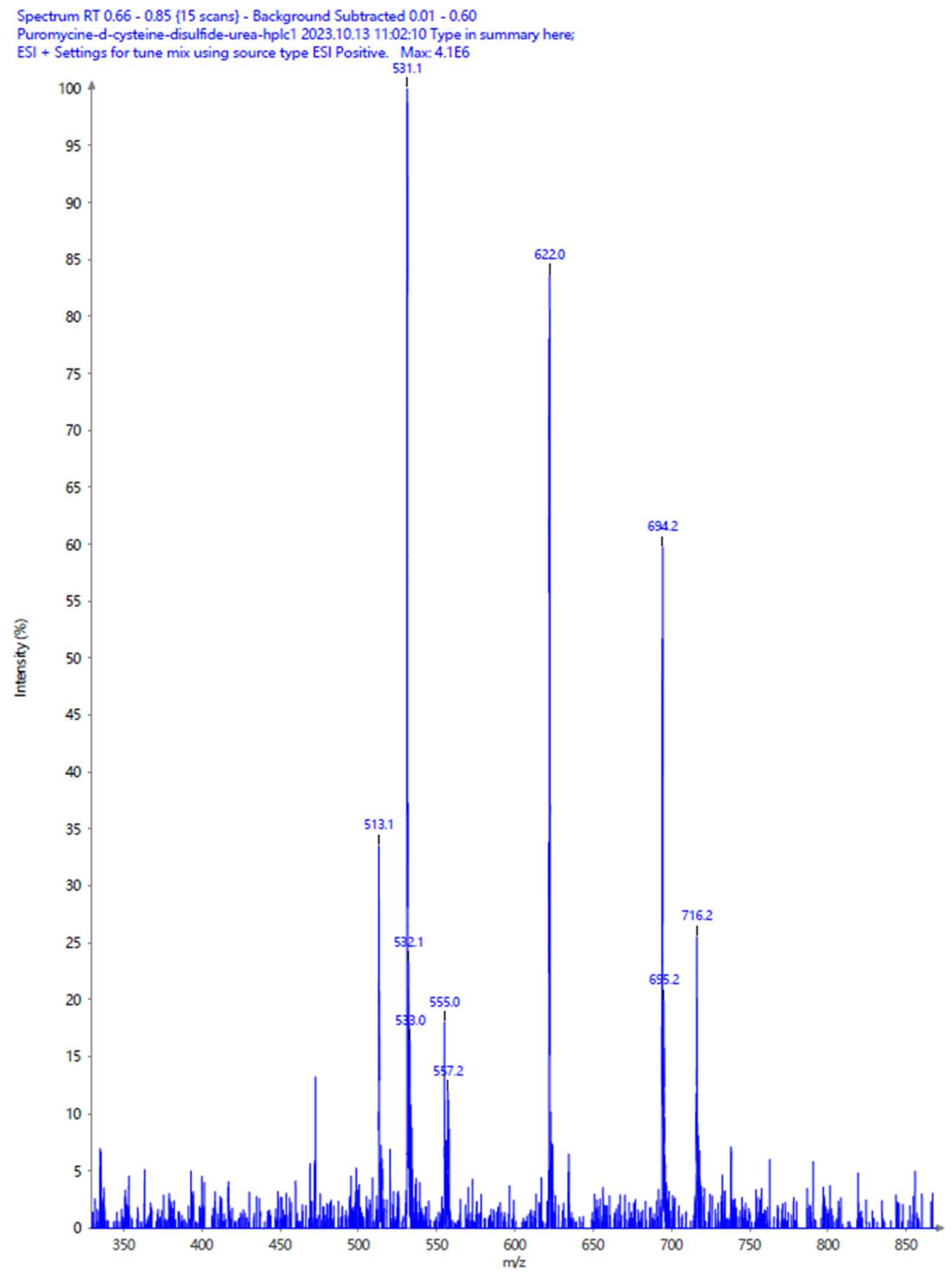


ESI-Mass spectrum for puromycin-cystamine-D-cysteine disulfide (**cys-Puro**) (m/z 694 for [M+H]$^+$).

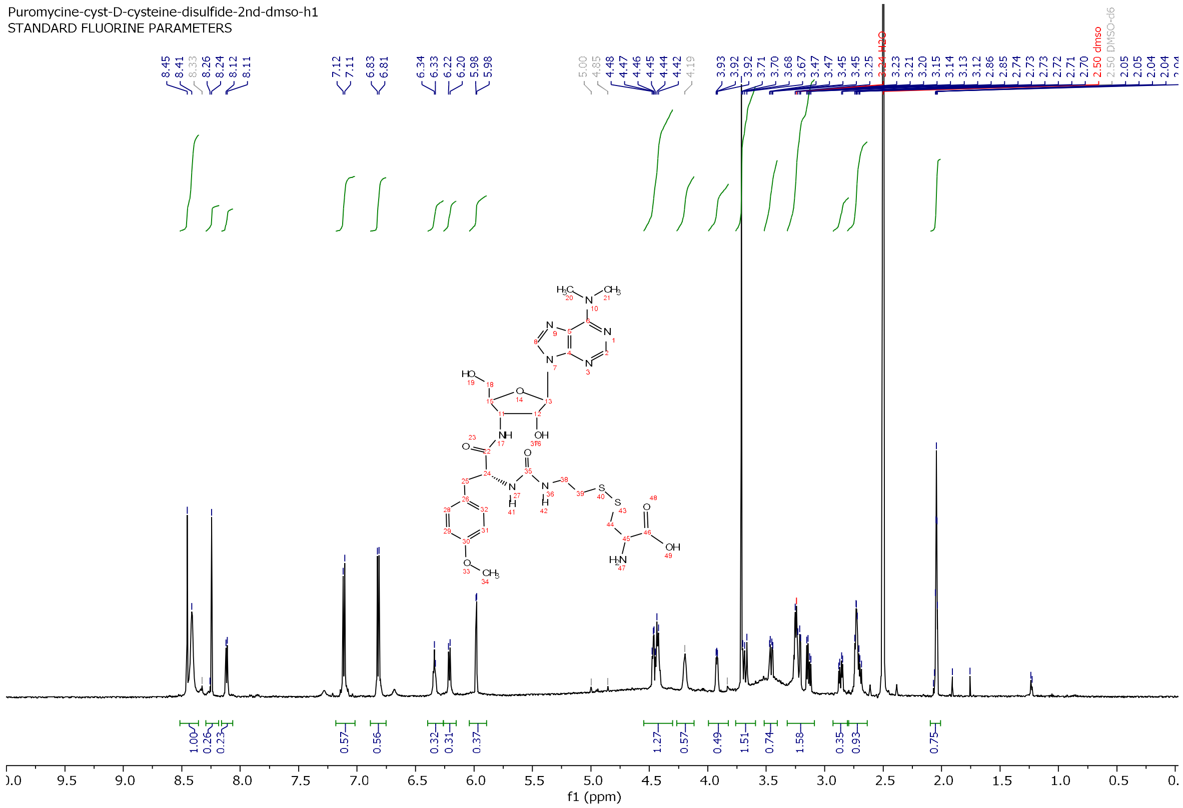


[1]H NMR spectrum of puromycin-cystamine-D-cysteine disulfide (**cys-Puro**).

**Rifamycin-cystamine-D-cysteine disulfide (cys-Rifa).**

**Synthesis of rifamycin-cystamine-D-cysteine disulfide (cys-Rifa).**

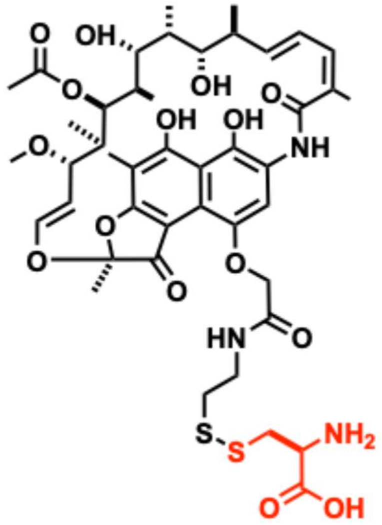


Synthesis of rifamycin-cystamine-D-cysteine disulfide (**cys-Rifa**) was carried out as outlined in the synthetic scheme below. Rifamycin B (25.1 mg, 0.033 mmol) was dissolved in anhydrous THF (3.0 mL). To this solution, 2-(pyridine-2-yldisulfaneyl)ethan-1-amine (11 mg, 0.05 mmol), DIC (10.4 mg, 0.05 mmol), and triethylamine (10 µL) were added. The reaction mixture was stirred at room temperature overnight. The following day, ESI–MS confirmed consumption of the starting material and formation of the intermediate pyridyl–disulfide conjugate (m/z 925 $[M+H]^+$). Volatiles were removed under reduced pressure using a rotary evaporator, and the residue was triturated with diethyl ether (2 × 5 mL). The ether layer was discarded, and the remaining solid was redissolved in $CH_3OH$ (2 mL). An aqueous solution of D-cysteine (25.0 mg, 0.2 mmol in 1.0 mL deionized $H_2O$) was then added, and the mixture was stirred for 2 h at room temperature. ESI–MS analysis indicated formation of the desired rifamycin-cystamine-D-cysteine (m/z 934 $[M+H]^+$).The reaction mixture was concentrated under reduced pressure to remove $CH_3OH$, and the residue was dissolved in $CH_3CN/H_2O$ (1:4, 10 mL). The solution was purified by preparative RP-HPLC on a C18 silica column. Fractions containing the desired compound, as determined by ESI–MS, were combined and concentrated to remove $CH_3CN$. The remaining aqueous solution was frozen at –80 °C and lyophilized to yield a colorless solid (20.1 mg).The compound was isolated as a trifluoroacetate salt and characterized by mass spectrometry. Purity of the final product was confirmed by analytical RP-HPLC.

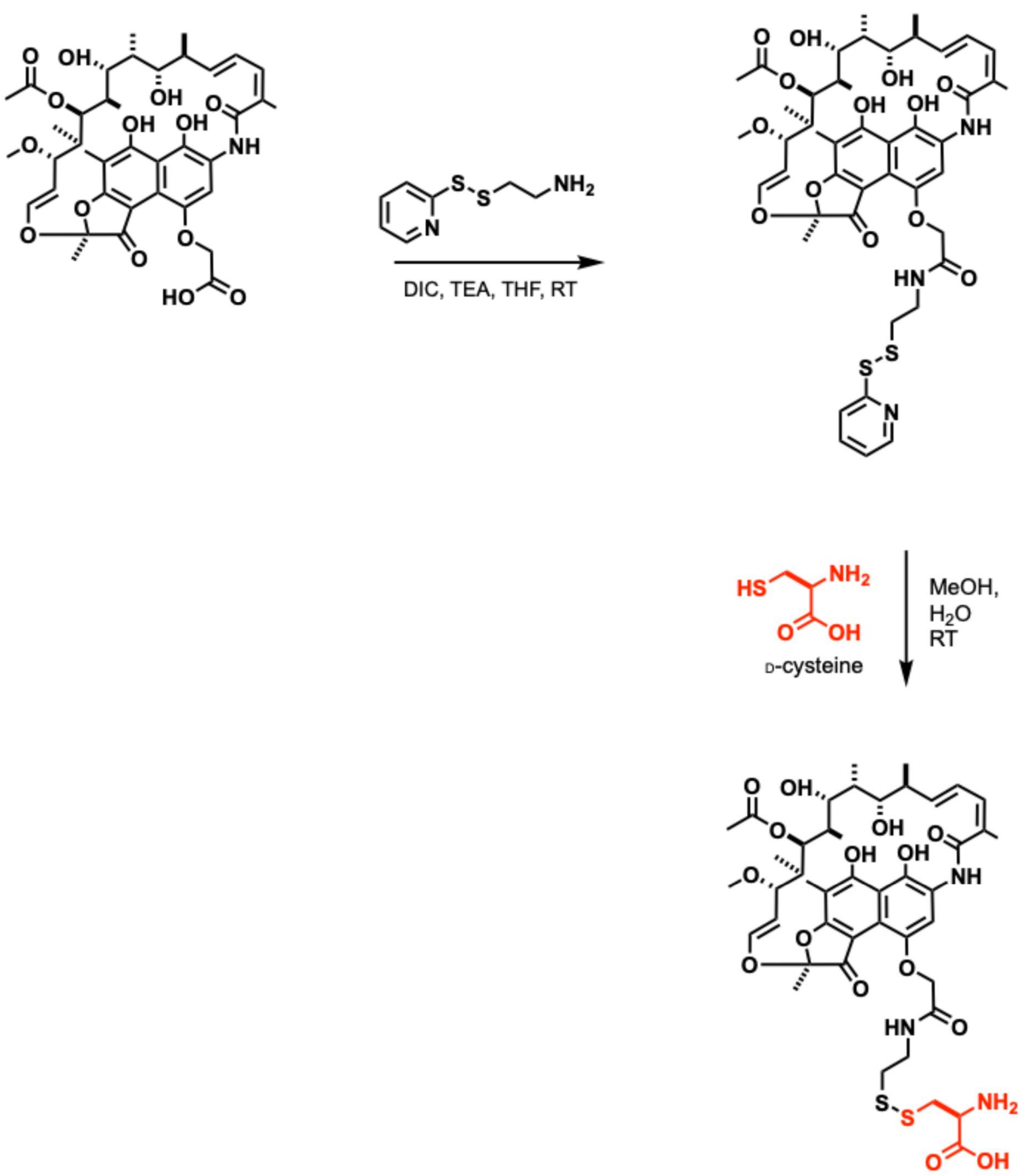


Synthesis scheme for rifamycin-cystamine-D-cysteine disulfide (**cys-Rifa**).

**Characterization of rifamycin-cystamine-D-cysteine (cys-Rifa).**

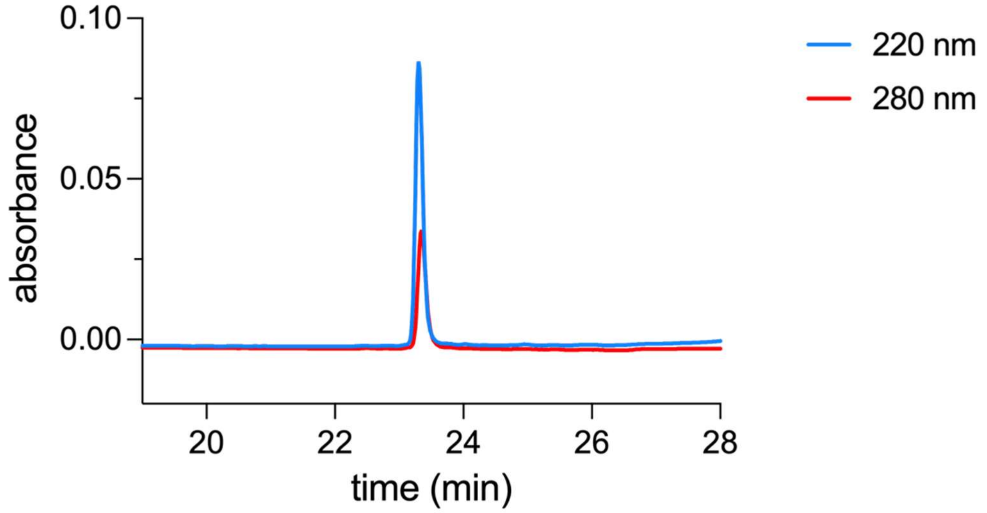


Analytical HPLC chromatogram rifamycin-cystamine-D-cysteine disulfide (**cys-Rifa**).

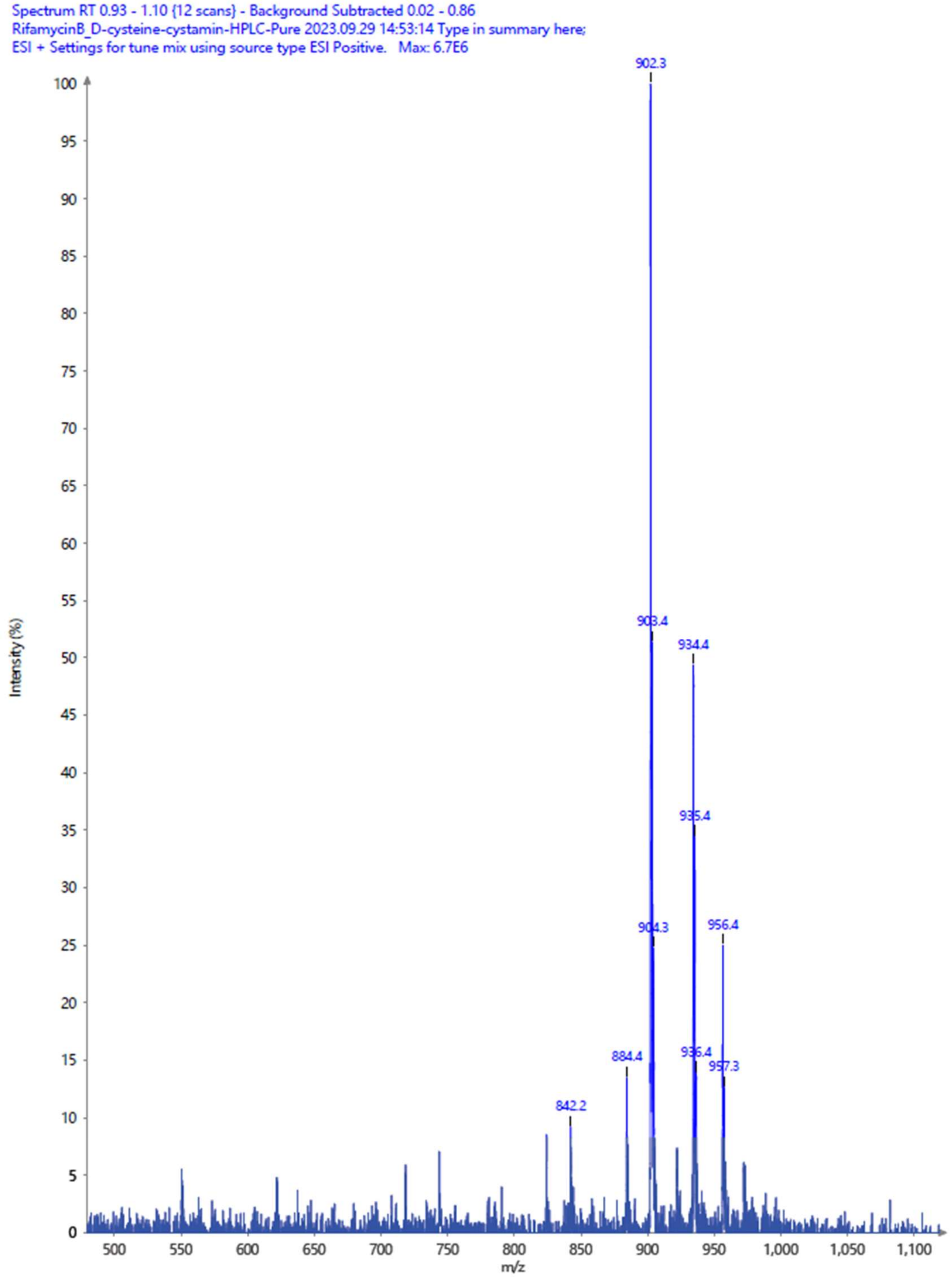


ESI-Mass spectrum for rifamycin-cystamine-D-cysteine disulfide (**cys-Rifa**) (m/z 934 for $[M+H]^+$).

### REFERENCES


(1) Dragset, M. S.; Ioerger, T. R.; Loevenich, M.; Haug, M.; Sivakumar, N.; Marstad, A.; Cardona, P. J.; Klinkenberg, G.; Rubin, E. J.; Steigedal, M.; Flo, T. H. Global Assessment of Mycobacterium Avium Subsp. Hominissuis Genetic Requirement for Growth and Virulence. *mSystems* **2019**, *4* (6), 10.1128/msystems.00402-19. https://doi.org/10.1128/msystems.00402-19.

(2) Vilchèze, C.; Copeland, J.; Keiser, T. L.; Weisbrod, T.; Washington, J.; Jain, P.; Malek, A.; Weinrick, B.; Jacobs, W. R. Rational Design of Biosafety Level 2-Approved, Multidrug-Resistant Strains of Mycobacterium Tuberculosis through Nutrient Auxotrophy. *mBio* **2018**, *9* (3), 10.1128/mbio.00938-18. https://doi.org/10.1128/mbio.00938-18.

(3) Kelly, C. N.; Townsend, C. E.; Jain, A. N.; Naylor, M. R.; Pye, C. R.; Schwochert, J.; Lokey, R. S. Geometrically Diverse Lariat Peptide Scaffolds Reveal an Untapped Chemical Space of High Membrane Permeability. *J. Am. Chem. Soc.* **2021**, *143* (2), 705–714. https://doi.org/10.1021/jacs.0c06115.

(4) Shahzad, S. A.; Sarfraz, A.; Yar, M.; Khan, Z. A.; Naqvi, S. A. R.; Naz, S.; Khan, N. A.; Farooq, U.; Batool, R.; Ali, M. Synthesis, Evaluation of Thymidine Phosphorylase and Angiogenic Inhibitory Potential of Ciprofloxacin Analogues: Repositioning of Ciprofloxacin from Antibiotic to Future Anticancer Drugs. *Bioorganic Chem.* **2020**, *100*, 103876. https://doi.org/10.1016/j.bioorg.2020.103876.
